# SLAP controls mTORC2 integrity *via* UBE3C-mediated mLST8 ubiquitination to mediate its tumour suppressive function in colorectal cancer

**DOI:** 10.1101/2025.03.10.642361

**Authors:** Rudy Mevizou, Dana Naim, Florent Cauchois, Cécile Naudin, Georgia Greaves, Kevin Espie, Bastien Felipe, Valérie Simon, Yvan Boublik, Julie Nguyen, Serge Urbach, Serge Roche, Audrey Sirvent

## Abstract

The mechanistic target of rapamycin complex 2 (mTORC2) signaling pathway, which regulates cell growth and migration, exhibits oncogenic function in colorectal cancer (CRC). mTORC2 signaling is primarily activated by a complex assembly of mTOR, RICTOR, SIN1, and mLST8, but how dysregulation of this mechanism contributes to its oncogenic function remains elusive. Here, we show that the Src-Like Adaptor Protein (SLAP), a negative regulator of tyrosine kinase signaling receptors, controls mTORC2 integrity to mediate its tumor-suppressive function in CRC. Mechanistically, SLAP interacts with mLST8 and promotes non-degradative ubiquitination, thereby reducing mTORC2 integrity and mTORC2-AKT signaling. The E3 ubiquitin ligase UBE3C was identified as a novel SLAP interactor involved in this ubiquitination process. Functionally, SLAP inhibition of CRC cell growth and invasion was dependent upon mTORC2 signaling inhibition. In immunodeficient mice CRC xenografts, SLAP depletion enhanced mTORC2 activity and sensitized CRC cells to mTOR catalytic inhibitors. Thus, our study uncovers a novel SLAP-mediated ubiquitination mechanism of mTORC2 dysregulation with potential therapeutic utility in CRC.

## INTRODUCTION

The mechanistic target of rapamycin (mTOR) is a key regulator of eukaryotic cell growth and metabolism induced by growth factors and nutrient availability^1,2^. mTOR is a serine/threonine protein of the phosphoinositide 3-kinase (PI3K)-related protein kinases (PIKK) family that mediates its cellular functions through two distinct complexes, mTOR complex 1 (mTORC1) and mTOR complex 2 (mTORC2)^1,2^. mTOR and mLST8 are core components of both complexes, while RAPTOR is a specific constituent of mTORC1 and SIN1 and RICTOR of mTORC2. mTORC1 primarily controls protein synthesis and anabolic metabolism, whereas mTORC2 is involved in cell survival, proliferation and cytoskeletal dynamics^1,2^. Although the regulation of mTORC1 has been extensively studied, much less is known about the regulation of mTORC2^1,3^. It is well established that mTORC2 is activated by a PI3K-dependent growth factor receptor signaling and mediates its cellular function by activating AGC (PKA, PKC and PKG) protein kinases, in particular by phosphorylating AKT at Ser473 to activate cell survival and migration^1,3^. mTORC2 activity is frequently dysregulated in human cancers, particularly in colorectal cancer (CRC), where it plays an important oncogenic role^4–7^. Notably, mTORC2 overactivation is involved in CRC cell proliferation, dissemination and therapeutic resistance^4,5,7,8^. Consequently, mTORC2 has been defined as an attractive therapeutic target in CRC. However, developed mTOR inhibitors have shown variable efficacy in CRC due to lack of specificity or induction of resistance mechanisms that overcome mTOR kinase inhibition^9–12^. Deciphering the underlying mechanism of mTORC2 oncogenic activation may lead to more effective mTORC2-based therapy.

mTORC2 signaling is primarily activated by complex assembly^1,13^. Ubiquitination has emerged as a key regulatory mechanism of this molecular process. For example, the co-chaperone TELO2-TTI1-TTI2 (TTT) complex defines a key control mechanism for the stabilization of PIKKs during translation, including mTORC2^3,13^. Its degradation by the ubiquitin ligase SCFFbxo9 has been reported to regulate mTORC2 abundance and survival AKT signaling in multiple myeloma^14^. Similarly, specific mTORC2 components (e.g. RICTOR and mTOR) have been subjected to ubiquitination-dependent degradation, affecting complex integrity and signaling^6,15^. Interestingly, recent findings have uncovered a non-degradative ubiquitylation mechanism of mTORC2 disassembly, *via* inhibition of the specific scaffolding function of mLST8 in this complex. mLST8 ubiquitination on Ub-K63, is mediated by TRAF2 and is overridden by the growth factor-responsive deubiquitinase OTUD7B^16^. A similar mechanism has been reported for RICTOR *via* the deubiquitinase USP9X^17^, suggesting a central mechanism of mTORC2 regulation. However, the contribution of the mechanism to the oncogenic function of mTORC2, particularly in CRC, is poorly characterized.

The Src-Like-Adaptor Protein (SLAP) belongs to the family of small adaptor proteins and is a negative regulator of tyrosine kinase (TK) signaling receptors, particularly those activated by immune antigens and growth factors^18,19^. Functionally, SLAP regulates the development and activation of thymocytes, where it is abundantly expressed^20,21^. It also negatively regulates Receptor TK (RTK) signaling leading to inhibition of fibroblast cell proliferation and migration^22–24^. Mechanistically, SLAP interacts with the E3 ligase CBL to induce receptor degradation^25–27^. SLAP shares high sequence homology with the N-terminus of TK SRC, including a myristoylation site and an SH3 and SH2 domain, hence its name^18^. Because of this homology, SLAP also inhibits SRC-like signaling possibly by preventing SRC-RTK association and/or promoting degradation of SRC-like substrates *via* CBL, such as the LCK substrate ZAP70 during TCR signaling^22,23,28^. In a previous study, we reported that SLAP is also abundantly expressed in colonic epithelium and its expression level is significantly downregulated in CRC^29^. We also uncovered a tumor suppressor function of SLAP in CRC involving degradation of the RTK and SRC oncogenic substrate EPHA2. This activity did not require CBL, but required interaction with the ubiquitination factor UBE4A, revealing the diversity of ubiquitination factors used by SLAP to mediate its inhibitory function^26^. Whether SLAP targets additional oncogenic signaling pathways to control CRC development remains to be determined.

In this study, we searched for additional SLAP signaling targets in CRC cells by proteomics. We show that mTORC2 is a major interactor of SLAP mediated by mLST8 binding. SLAP acts as an mTORC2 signaling inhibitor by promoting complex disassembly *via* non-degradative mLST8 ubiquitination and the novel SLAP interactor and E3 ubiquitin ligase UBE3C. Functionally, SLAP anti-oncogenic effect in CRC cells is mediated by mTORC2 inhibition. Collectively, our study identifies a novel dysregulation mechanism of mTORC2 complex integrity induced by SLAP downregulation with potential therapeutic utility in CRC.

## MATERIALS AND METHODS

Antibodies: anti-SLAP (C-19) was from Santa Cruz; anti-EPHA2 (D4A2), anti-mTOR (7C10), anti-LST8 (#3227), anti-Raptor (24C12), anti-S6K1 (#9202), anti-pT389S6K1 (108D2), anti-AKT (#9272), anti-pS473AKT (D9AE), anti-Rictor (53A2 et D16H9), anti-NDRG1 (D8G9), anti-pT346 NDRG1 (D98G11), anti-ULK1 (8054S) and anti-pS555 ULK1 (14202T) were from Cell Signaling Technology; anti-Sin1 (D7G1A), anti-TRAF2 (4724), anti-Rictor (A300-459A and A300-458A) and anti-LST8 (A300-679A) were from Bethyl Labs; anti-Flag M2, anti-Flag M2 agarose and anti-Actin were from Sigma Aldrich; anti-Flag magnetic beads was from Termofisher (#A36797); anti-Telo2 (15975-1-AP) was from Protein Tech; anti-pan Ubiquitin was from Zymed; anti-Rictor (NBP1-51645SS) used for PLA was from NOVUS.

Drugs: mTOR inhibitors KU-0063494, Temsilorimus (TS) and AZD2014 were from Selleckchem (USA); PP242/Torkinib (#T2414) was from TargetMOL.

Vectors: pcDNA3 and retroviral pMX-pS-CESAR vectors expressing SLAP-Flag, SLAP SH3*(P73L)-Flag, SLAP SH2*(R111E)-Flag and SLAP SH3*SH2*-Flag (a gift of A. Weiss, University of California, San Francisco, USA) and SLAP N32-Flag were described in ^29^. pMXs-DsRed Express was a gift from Shinya Yamanaka (Addgene plasmid # 22724; www.addgene.org). Human RICTOR, RAPTOR, mTOR, SIN1, LST8, TTI1 and TEL2 cDNA was obtained from an ORFeome library (Montpellier genomic platform, GENOMIX, www.mgc.cnrs.fr) and subcloned in HA and Flag-tagged form in pcDNA3 and retroviral pMX-pS-CESAR expression vectors. Pol, GAG, Env expression vectors were described in ^30^. pRBG4 Ubiquitin-myc-6His WT K48R and K63R mutants were from J. Pierre (IGR, Université Paris-XI, France). pcDNA3-LST8 mutants and pRBG4 Ub-myc-6His K29R and K33R mutants were obtained using the QuickChange Site-Directed Mutagenesis Kit (Agilent) using the following oligonucleotides: mLST8 K86R: (F) CAGCTACGACGGCGTCAACAGGAACATCGCGTCTGTGGGC, (R) GCCCACAGACGCGATGTTCCTGTTGACGCCGTCGTAGCTG; K215R: (F) CCCAGCTCATCCCCAAGACTAGGATCCCTGCCCACACGCGC, (R) GCGCGTGTGGGCAGGGATCCTAGTCTTGGGGATGAGCTGGG; K245R: (F) GCTCGGCTGATCAGACGTGCAGGATCTGGAGGACGTCCAACTTC, (R) GAAGTTGGACGTCCTCCAGATCCTGCACGTCTGATCAGCCGAGC; Ub K29R: (F) CGAAAATGTGAAGGCCAGGATCCAGGATAAGGAAGGC, (R) GCCTTCCTTATCCTGGATCCTGGCCTTCACATTTTCG; Ub K33R: (F) GTGAAGGCCAAGATCCAGGATAGGGAAGGCATTCCCCCCGACC, (R) GGTCGGGGGGAATGCCTTCCCTATCCTGGATCTTGGCCTTCAC. siRNA: siSLA_h #1 (ON-TARGETplus Human SLA (6503) siRNA J-010815-07; Dharmacon), siSLA_h #2 (ON-TARGETplus Human SLA (6503) siRNA J-010815-08; Dharmacon), siLST8_h #1 (ON-TARGETplus Human MLST8 (64223) siRNA J-016093-05; Dharmacon), siLST8_h #2 (ON-TARGETplus Human MLST8 (64223) siRNA J-016093-06; Dharmacon), siLST8_h #3 (ON-TARGETplus Human MLST8 (64223) siRNA J-016093-07; Dharmacon), siLST8_h #4 (ON-TARGETplus Human MLST8 (64223) siRNA J-016093-08; Dharmacon), siLST8_h #5(UGACGGAGCUGAGCAUCAA), siRictor_h #1; ON-TARGETplus Human RICTOR (253260) siRNA J-016984-05, Dharmacon), siRictor_h #2 (ON-TARGETplus Human RICTOR (253260) siRNA J-016984-06, Dharmacon), siRaptor_h#1(ON-TARGETplus Human RPTOR (57521) siRNA J-004107-05, Dharmacon), siRaptor_ h #2 (ON-TARGETplus Human RPTOR (57521) siRNA J-004107-05; Dharmacon), siUBE3C_h #1 (GCCAGACAUUACUACUUCCUA), siTRAF2_h #1 (GGACCAAGACAAGAUUGAA), siTRIM32_h #1 (GAUCAGGGGUGGUCAAAUA). shRNA vectors: control (CTRL, GACACTCGGTAGTCTATAC), shRICTOR (ACTTGTGAAGAATCGTATC) and shRAPTOR (GGCTAGTCTGTTTCGAAATTT) were subcloned in pSUPER.retro.neo.GFP. shSLAP (GACCTGGTGAACCACTATT) in pSiren-retroQ was described in ^30^.

### Cell cultures, retroviral infections and transfections

HCT116, HT29, SW480, DLD1, LS174T, LoVo, SW620 and HEK293T cell lines were obtained from ATCC (Rockville, MD). DLD1 cells stably expressing TEL2-degron (TEL2^AID^) (a gift of Dr D. Helmlinger, CRBM, Montpellier, France) were described in ^31^. Cells were cultured at 37°C and 5% CO_2_ in a humidified incubator in Dulbecco’s Modified Eagle’s Medium (DMEM) GlutaMAX (Invitrogen) supplemented with 10% fetal calf serum (FCS), 100 U/ml of penicillin and 100 µg/ml of streptomycin as described in ^29^. Retroviral production and cell infection were performed as described in ^29^. Stable cell lines were obtained by GFP and/or mRFP-fluorescence-activated cell sorting. Transient plasmid transfections in HEK293T cells were performed with the jetPEI reagent (Polyplus-transfection) according to the manufacturer’s instructions. For siRNA transfection, 2.10^5^ cells were seeded in 6-well plates and transfected with 20 nmol of siRNA and 9 µl of Lipofectamine RNAi Max according to the manufacturer’s protocol (ThermoFisher Scientific).

### Soft agar, invasion, colonospheres and tumoroid formation assays

Colony formation in soft-agar assays were performed from 1 000 to 2 000 cells per well that were seeded in 12-well plates in 1ml DMEM containing 10% FCS and 0.33% agar (Sigma Aldrich)on a layer of 1ml of the same medium containing 0.7% agar as described in ^29,32^. After 18-21 days, colonies with > 50 cells were scored as positive. Cell invasion assay was performed in Fluoroblok invasion chambers (BD Bioscience) using 50 000 cells in the presence of 100 µl of 1-1.2 mg/ml Matrigel (BD Bioscience) as in ^29,32^. After 24-48h, cells were labelled with Calcein AM (Sigma Aldrich) and invasive cells were photographed using the EVOS FL Cell Imaging System (ThermoFisher Scientific). Quantification of the number of invasive cells per well was done with the Image J software (https://imagej.nih.gov/ij/). For colonosphere formation, 100 cells were seeded in ultra-low attachement plates (Corning) in 100 μl of DMEM/F12 (Life technologies) supplemented with 2mM L-glutamine, N-2 (Life technologies), EGF (20ng/ml, Bio-techne) and FGF (10ng ml-1 - Bio-techne) for 7 days. Tumoroids formation was performed as follows: 3 000 HCT116 cells were resuspended in advanced DMEM/F12 (Life Technologies) supplemented with 1% Penstrep, 2mM L-glutamine, N-2 (Life Technologies) and Matrigel (Corning) (1:2 ratio) prior to plating in 24-well plates. After polymerization of Matrigel, 0.5 mL DMEM/F12 supplemented with 20 ng/ml EGF and 10 ng/ml FGF (Biotechne) was added. Tumoroids were cultured at 37°C and 5% CO2 in a humidified incubator and treated or not with indicated drug (twice per week) for 7 days before size quantification by microscopy.

### RNA extraction and RT-quantitative PCR

mRNA was extracted from cell lines and tissue samples using the RNeasy plus mini kit (Qiagen) according to the manufacturer’s instructions. RNA (1µg) was reverse transcribed with the SuperScript VILO cDNA Synthesis Kit (Invitrogen). Quantitative PCR (qPCR) was performed with the SyBR Green Master Mix in a LightCycler 480 (Roche). Expression levels were normalized with the Tubulin or GAPDH human housekeeping gene. Primers used for qPCR are: human TRIM32: (F) TGT GGT TTG GTG TTA TGT GAG C, (R) TAA GTT CCC GCA GAC GAG TTA; human TRAF2: (F) GCT CAT GCT GAC CGA ATG TC, (R) GCC GTC ACA AGT TAA GGG GAA; human UBE3C: (F) AAA GCA GAT AAG GTC ACT CAG C, (R) CAA AAG GCA ACT CTG TCA GGA.

### Biochemistry

Immunoprecipitation (IP) and immunoblotting (WB) were performed as described in ^29^. Briefly, cells were lysed at 4°C with Triton-lysis buffer (20 mM Hepes pH7.5, 150 mM NaCl, 0.5% Triton X-100, 6 mM octyl β-D-glucopyranidose, 10 µg/ml aprotinin, 20 µM leupeptin, 1 mM NaF, 1 mM DTT and 100µM Na3VO4) unless specified. For WB, antibodies were used at a dilution of 1:1000. For mTORC2 co-immunoprecipitation experiments cells were lysed in CHAPS buffer [40 mM HEPES pH 7.5, 120 mM NaCl, 1 mM EDTA, 0.6% CHAPS 10 mM pyrophosphate, 10 mM glycerophosphate, 50 mM NaF, 1.5 mM Na3VO4, 1% Triton X-100, and EDTA-free protease inhibitors (Roche)] as described in ^33^. Cycloheximide (50 μg/ml) cell treatment were performed as in ^29^. For ubiquitination assays, cells were lysed at room temperature with Guanidine Buffer (6 M guanidinium-HCl, 0.1 M Na_2_HPO_4_/NaH_2_PO_4_, 0.01 M Tris/HCl, pH 8.0, 5 mM imidazole and 10 mM β-mercaptoethanol). Ubiquitinated proteins were purified using Ni-NTA agarose beads (Qiagen). Complexes were then washed in Guanidine buffer and then Urea buffer (8 M urea, 0.1 M Na_2_HPO_4_/NaH_2_PO_4_, 0.01 M Tris/HCl, pH 8.0, 10 mM β-mercaptoethanol, 0.1% Triton X-100) and analyzed by WB. GST pull-down assays were performed as described in ^29^. The expression of fusion proteins in *Escherichia coli* (BL21 strain) was induced by incubation with 0.2mM isopropyl β-D-thiogalactopyranoside (IPTG) at 25°C for 3h. The expressed proteins were purified as described in ^44^ and bound to Glutathione Sepharose 4B beads (GE Healthcare). 15 µg of GST alone, GST-SLAP N32 and GST-SLAP N3*2 bound to Glutathione Sepharose beads were incubated with transfected HEK293T cell-lysates at 4°C for 2h. Complexes were washed in PBS and analyzed by WB. 20-50 µg of whole cell lysates were loaded on SDS-PAGE gels and transferred onto Immobilon membranes (Millipore). Detection was performed using the ECL System (Amersham Biosciences).

### Proteomics

The Proteome Profiler Human Phospho-Kinase Array Kit was purchased from R&D Systems. CRC cells were lysed and 300 μg of protein lysates were used for WB as described in ^34^. Signals were quantified with the Amersham Imager 600 (GE Healthcare) from two independent biological replicates. SILAC-based interactomic analysis were performed as described in ^29^. Briefly, SW620 cells were cultured in SILAC DMEM (Pierce) without Lysine (Lys) and Arginine (Arg) and supplemented with 4mM L-glutamine, 10% dialyzed FBS (Invitrogen), 0.084g/l Arg and 0.146g/l Lys. Heavy (^13^C ^15^N -Arg and ^13^C ^15^N -Lys, from EurisoTop) or unlabeled amino acids (light Arg and Lys, from Sigma Aldrich) were used. After 3 weeks of metabolic labeling, cells were lysed in lysis buffer. Heavy and light lysates (30 mg of protein) were mixed and immunoprecipitated with anti-Flag M2 agarose overnight at 4°C. Peptides obtained after digestion of IP were analyzed using a Qexactive-HFX system coupled with a RSLC-U3000 nano HPLC. Samples were desalted and pre-concentrated on-line on a Pepmap® precolumn (0.3 mm x 10 mm). A gradient consisting of 6-25% B for 100 min, 25-40% B for 20 min, 40-90% B for 2 min (A = 0.1% formic acid; B = 0.1 % formic acid in 80% acetonitrile) at 300 nl/min was used to elute peptides from the capillary reverse-phase column (0.075 mm x 250 mm; Pepmap®, ThermoFisherScientific), fitted with a stainless steel emitter (Thermo Sientific). Spectra were acquired with the instrument operating in the data-dependent acquisition mode throughout the HPLC gradient. MS scans were acquired with resolution set at 60,000. Up to twelve of the most intense ions per cycle were fragmented and analyzed using a resolution of 30,0000. Peptide fragmentation was performed using nitrogen gas on the most abundant and at least doubly charged ions detected in the initial MS scan and an dynamic exclusion time of 20s. Analysis was performed with the MaxQuant software (version 2.0.3.0). All MS/MS spectra were searched using Andromeda against a decoy database consisting of a combination of the *Homo sapiens* reference proteome (release 2024_01 www.uniprot.org) and of 250 classical contaminants, containing forward and reverse entities. A maximum of two missed cleavages was allowed. The search was performed with oxidation (M) and acetyl (protein N-term) as variable modifications, and carbamidomethyl (C) as fixed modification. FDR was set at 0.05 for peptides and proteins, and the minimum peptide length was 7.

### PLA and cell imaging

Proximity Ligation Assay (PLA) was performed according to the manufacturing kit protocol (#NF.100.2, Navinci). Briefly, SW620 cells plated on glass coverslips for 48 hours were fixed with 4% paraformaldehyde, 0.5% triton during 20 min at room temperature. Glass slides were blocked in the Navinci® Blocking Buffer (1x) during 1 hour at 37°C in a humidity chamber and then incubated with primary antibodies anti-RICTOR (NOVUS NBP1-51645SS, 1:400, mouse) and anti-mTOR (CST#2983, 1:400 rabbit) in the Navinci® Primary Antibody Diluent (1X) overnight at 4 °C. After the washing steps, glass coverslips were incubated 1 hour at 37°C in a humidity chamber with the Navenibodies diluted (1:40 each) in Navenibody Diluent. After three washes, three enzymatic reactions followed. The enzyme (A,B,C) was diluted (1:40) in the associated buffer (A,B,C) (1:5) and glass coverslips were incubated respectively 60 min, 30 min and 90 min at 37°C in a humidity chamber. Finally, after washes glass coverslips were mounted with the Duolink® In Situ Mounting Medium with DAPI (#DUO82040) and observed with an upright fluorescent microscope. Images were acquired using Zeiss Axioimager Z2 microscope (20-80 cells/field, 7-15 fields per coverslip), with objective plan-apochromat 63x/ 1.40 oil, then PLA signal area analysis was processed using Fiji software (https://fiji.sc/).

### Animal experiments

*In vivo* experiments were performed in compliance with the French guidelines for experimental animal studies (Direction des services vétérinaires, ministère de l’agriculture, France, APAFIS #35916-2022031511382008 v6, local animal house agreement number F3417216). 2 10^6^ SW620 cells (or derivatives) were subcutaneously injected in the flank of 5-week-old female athymic nude mice (Envigo). Tumor volumes were measured as the indicated intervals using calipers for 24 days. For drug treatment, HCT116 cells (or derivative) were first cultured during 2 weeks in suspension to form colonospheres in low attachment T150 flasks (Sigma); 150,000 dissociated cells (using accumax solution, Invitrogen) mixed Matrigel (Corning, ratio 1:1) were injected in 7-old female athymic nude mice (Charles River Laboratories, France). When tumor volume reached 50mm3, mice were randomized and treated with vehicle (0.5% methylcellulose/0.05% Tween80) or Torkinib (PP 242; 40 mg/kg) by oral gavage daily, 5 days/week for 21 days. Tumor xenografts were excised, weighed and cryopreserved or processed for subsequent IHC or protein tissue analysis by WB as described in ^29,32^. *In vivo* siRNA administration in *APC^Δ^*^14^*^/+^* mice was described in ^30^. Briefly, murine Slap AGAUUGGUAGCUUCAUGAU or control CGUACGCGGAAUACUUCGATT siRNAs (10 µg) were mixed and complexed at room temperature for 30 min with an equal volume of cationic liposomes (60 nmol, generous gift from V. Escriou, Paris Descartes University, France)^45^ in 0.9% NaCl incubated. Then, 3 week/old *APC^Δ^*^14^*^/+^* mice (a gift of C Perret, Cochin Institute, Paris) received i.p. injections of siRNA twice a week for 6 weeks. Five days after the last injection, intestinal tumors were counted, measured, removed and cryopreserved. equal volumes of cationic liposomes (60 nmol) and siRNA (10µg) in 0.9% NaCl were mixed and incubated at room temperature for 30min. Then, 3 week/old *APC^Δ^*^14^*^/+^* mice (a gift of C Perret, Institut Cochin, Paris) received i.p. injections of siRNA twice a week for 6 weeks. Five days after the last injection, intestinal tumors were counted, measured, removed and cryopreserved.

### Immunohistochemical analysis

For bright field immunochemistry, sections were treated with BIOXALL (Vector Laboratories) for 10 minutes before blocking and incubation with primary antibodies (anti-pSer473 AKT, D9AE 1:50 according to CST protocol). Secondary staining was detected with DAB (Vector Laboratories), and sections were counterstained with hematoxylin (Sigma-Aldrich) and mounted with mounting medium (Pertex). For histological examination, tissue sections were deparaffinized and stained with hematoxylin, eosin, and alcian blue.

### Statistical analysis

All analysis were performed using GraphPad Prism (9.3.1). Data are presented as the mean ± SEM from at least 3 independent experiments as stated. The two-tailed *t* test was used for between-group comparisons using the Mann-Whitney test was used. The statistical significance level is illustrated with p values: *p≤0.05, **p≤0.01, ***p≤0.001 (t-test).

## RESULTS

### SLAP interacts with mTORC2

To identify novel SLAP targets in CRC cells, we performed SILAC-based interatomic analysis of SW620 CRC cells overexpressing a SLAP-Flag construct (Fig. S1A). We found the mTORC2 components, namely mTOR, RICTOR, SIN1/MAPKAP1(SIN1), and mLST8 (retrieved in ≥2/3 independent experiments with ≥3 peptides), showing that mTORC2 is a novel SLAP target in CRC cells (Fig. S1 and Table S1). The PIKKs co-chaperones TELO2 and TTI1 were also identified, suggesting that SLAP may also bind to mTORC2 during complex formation (Fig. S1 and Table S1). SLAP interaction with all mTORC2 components was next confirmed biochemically from cell lysates of HT29 and SW620 cells overexpressing SLAP (Fig. 1A). The effect of SLAP on mTORC2 phospho-signaling in CRC cells was next assessed with a phospho-kinase arrays (Fig. S2A). The phosphorylation level of specific mTORC2 substrates (pS473 AKT, pS9 GK3β), but not of mTORC1 (pT398 S6K and pT412/S424 S6K), was reduced upon SLAP expression, whereas their levels were increased upon shRNA-mediated SLAP silencing in HCT116 CRC cells, which retain a substantial level of endogenous SLAP. This mTORC2 signaling inhibition by SLAP was confirmed by western blotting on specific substrates, including AKT and NDRG1, while the phosphorylation levels of the mTORC1 specific substrates S6K and ULK1 were unchanged^1^ (Fig. 1B). Consistent with this, mTORC2-AKT signaling induced by growth factor stimulation in SW620 and HCT116 cell-lines was modulated by SLAP expression, in contrast to mTORC1-S6K phospho-signaling (Fig. S2B and S2C). Thus, we concluded that SLAP specifically inhibits mTORC2 activity in CRC cells.

**Fig. 1:**
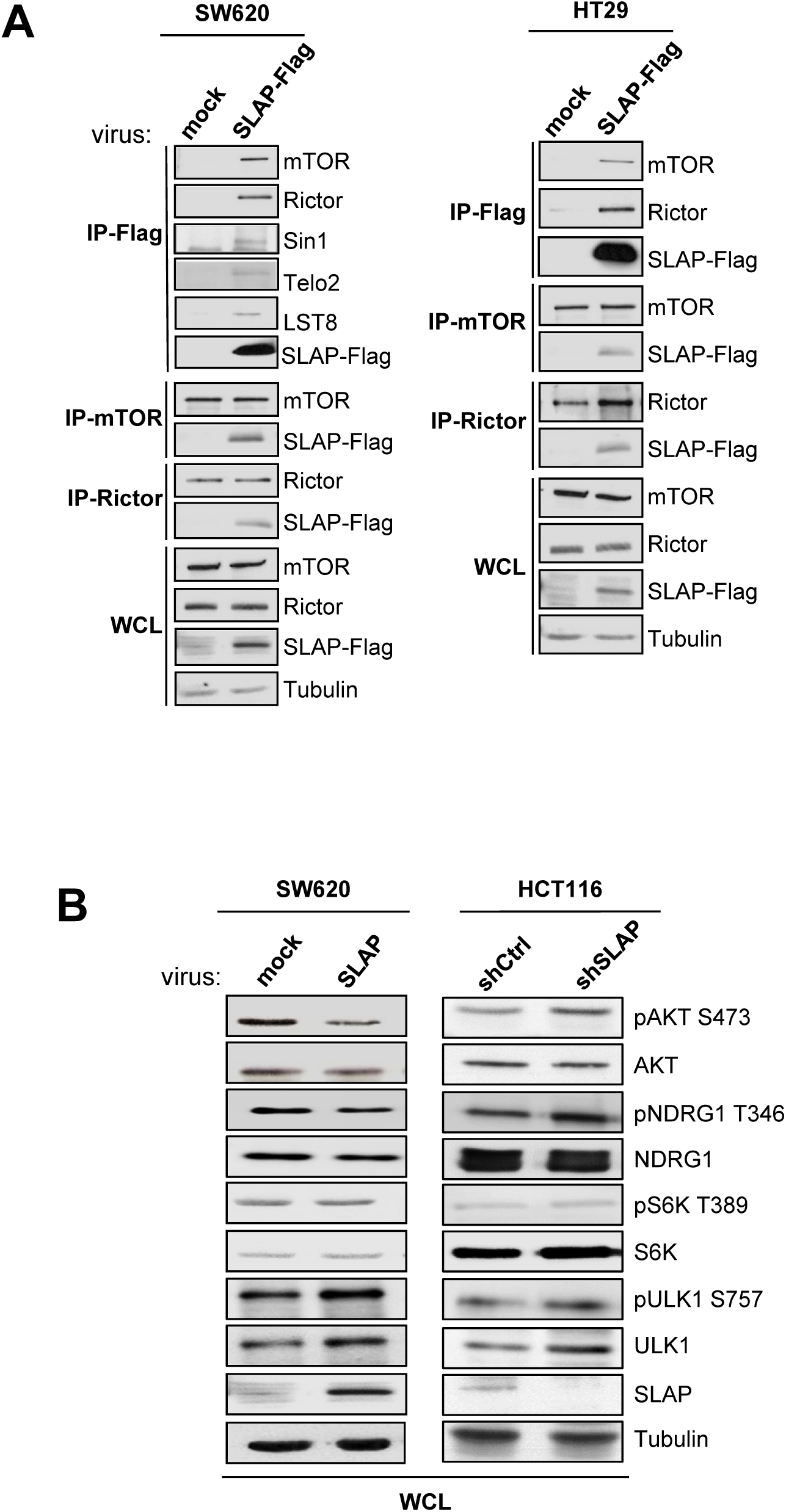
SLAP Interacts with mTORC2. **A** Biochemical validation of SLAP-mTORC2 complex formation in indicated CRC cells. **B.** Western blot (WB) analysis of SLAP-dependent phosphorylation of selected mTORC1/2 substrates in indicated CRC cells. Shown are representative WB from 3 independent experiments.

### mTORC2 inhibition mediates SLAP anti-oncogenic function

We next investigated the contribution of mTORC2 signaling to the anti-oncogenic function of SLAP in CRC cells. SLAP depletion in HCT116 cells leads to an increase in anchorage-independent growth in soft agar and cell invasion in Matrigel^29^ (Fig. 2A). This transformation-promoting effect was reversed by mTORC2 inhibition mediated by shRNA-mediated RICTOR knockdown. In contrast, mTORC2 inhibition had no significant effect in control cells, demonstrating the SLAP-dependent transforming effect. In addition, mTORC1 inhibition induced by shRNA-mediated RAPTOR depletion has no clear effect (Fig. 2A), confirming the mTORC2-dependent SLAP effect. Conversely, the inhibition of these transforming properties in HT29 cells by SLAP overexpression was overcome by RICTOR overexpression (Fig. 2B). RICTOR also restored mTORC2 activity as shown by pS473 AKT levels (Fig. 2B), consistent with an mTORC2-dependent rescue effect of RICTOR. Finally, anchorage-independent cell growth and invasive properties of the metastatic CRC cell line SW620 were inhibited by RICTOR, but not by RAPTOR depletion, revealing a specific mTORC2 oncogenic function in these cells (Fig. S3A and S3B). However, this mTORC2 function was not evident upon SLAP overexpression (Fig. S3A and S3B), confirming that the mTORC2 oncogenic function can be inhibited by SLAP expression in CRC cells.

**Fig. 2:**
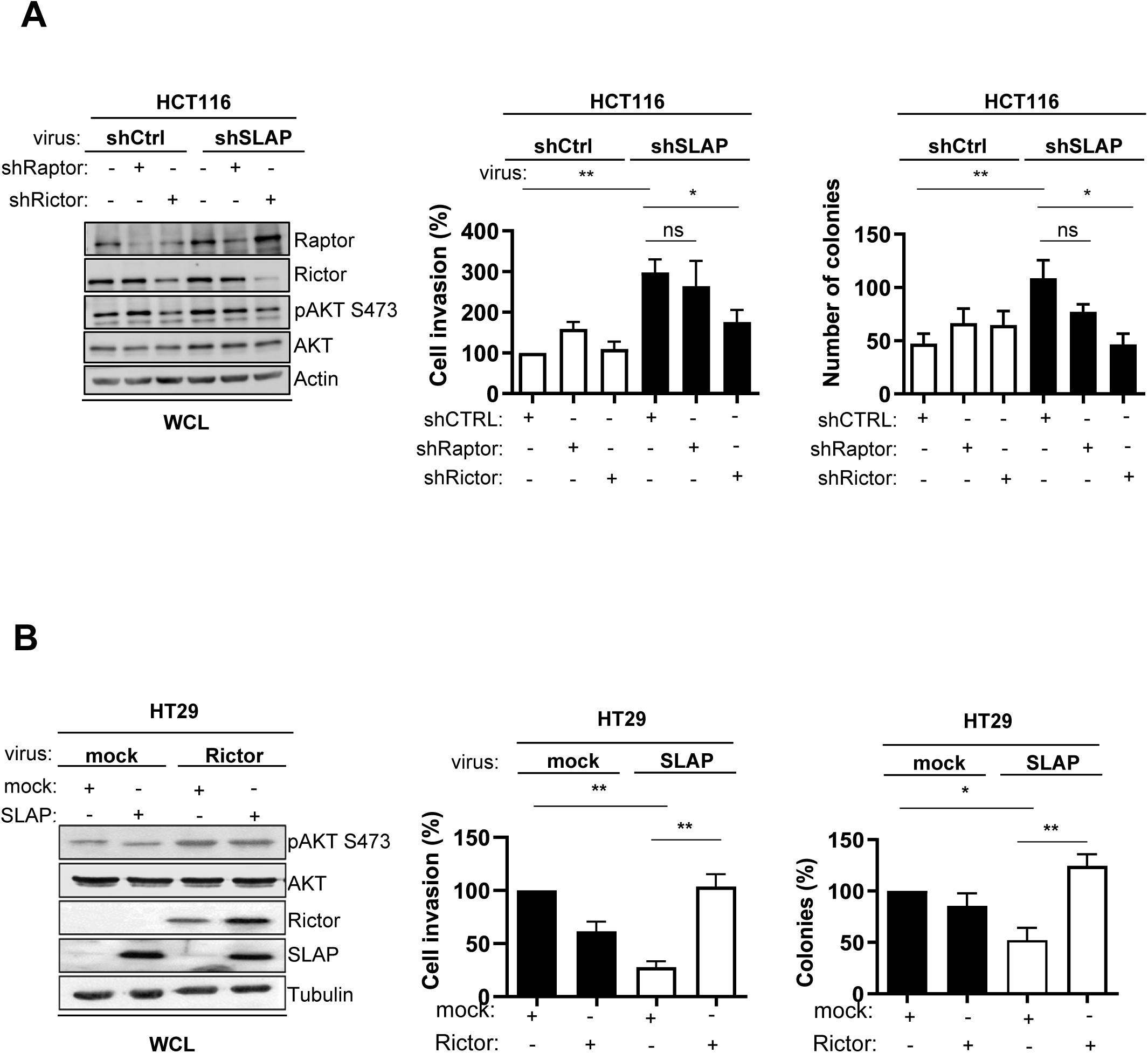
SLAP anti-oncogenic function is mediated by mTORC2 inhibition. **A.** HCT116 CRC cell transforming properties induced by SLAP depletion are dependent upon RICTOR but not RAPTOR expression. Left: WB of protein level in indicated HCT116 cells transduced with indicated retroviruses. Right: cell invasion in Matrigel (control: 100%) and colonies formation in soft agar (number) is shown (mean ± SEM, n=3-5). **B.** SLAP-dependent inhibition of HT29 cell transforming properties is overcome by RICTOR overexpression. Left: WB of indicated protein levels in HT29 cells transduced with indicated retroviruses. Right: cell invasion in Matrigel colony formation in soft agar (control: 100%) is shown. Is shown the mean ± SEM, n=3-5, *p≤0.05, **p≤0.01 (t-test).

### SLAP targets mLST8 to inhibit mTORC2 complex stability

To address the mechanism by which SLAP inhibits mTORC2 activity, we first examined the interaction between SLAP and each component of the complex, which were expressed in HEK293T cells. TELO2 and mLST8 were identified as the major SLAP interactors (Fig. S4A). Structure-interaction analysis of SLAP mutants harboring inactive mutations in the SH2 and/or SH3 domains (SH2*, SH3*, SH3*SH2*) or a deletion of the C-terminal domain (N32) revealed a major contribution of SLAP-SH3 and -SH2 in the association with the mTORC2 complex in SW620 cells (Fig. S4B). *In vitro* pull-down assays, using purified GST-SLAP N32 with lysates from HEK293T cells expressing TELO2 or mLST8, confirmed specific interactions with these mTORC2 components. mLST8 interaction was reduced with GST-SLAPN3*2, in contrast to TELO2, showing a SLAP-SH3-dependent interaction with mLST8 (Fig. S4C). Having demonstrated SLAP-SH3-dependent mTORC2 association in CRC cells, we hypothesized that mLST8 is involved in this molecular process. Consistent with this, siRNA-mediated depletion of mLST8 reduced SLAP association with RICTOR in both SW620 and HT29 cells (Fig. 3A). Functionally, mLST8 overexpression in HT29 and SW620 cells restored cell growth and invasion, which has been inhibited by SLAP expression (Fig. 3B and 3C), while siRNA-mediated mLT8 depletion reversed the transformed character of HCT116 cells that has been increased upon SLAP silencing. The depletion of mLST8 also reduced mTORC2 activity, as shown by pS473 AKT levels, consistent with an mTORC2-dependent effect of mLST8 depletion on these cellular responses (Fig. 3D). In contrast, TELO2 depletion did not reduce SLAP-RICTOR association in CRC cells (Fig. S5A), indicating that it is not involved in SLAP-mTORC2 complex formation. Furthermore, TELO2 overexpression in HT29 cells did not affect the SLAP-inhibitory effect on cell growth and invasion (Fig. S5B and S5C). Thus, we conclude that mLST8, but not TELO2, plays an important role in this SLAP-mTORC2 signaling process.

**Fig. 3:**
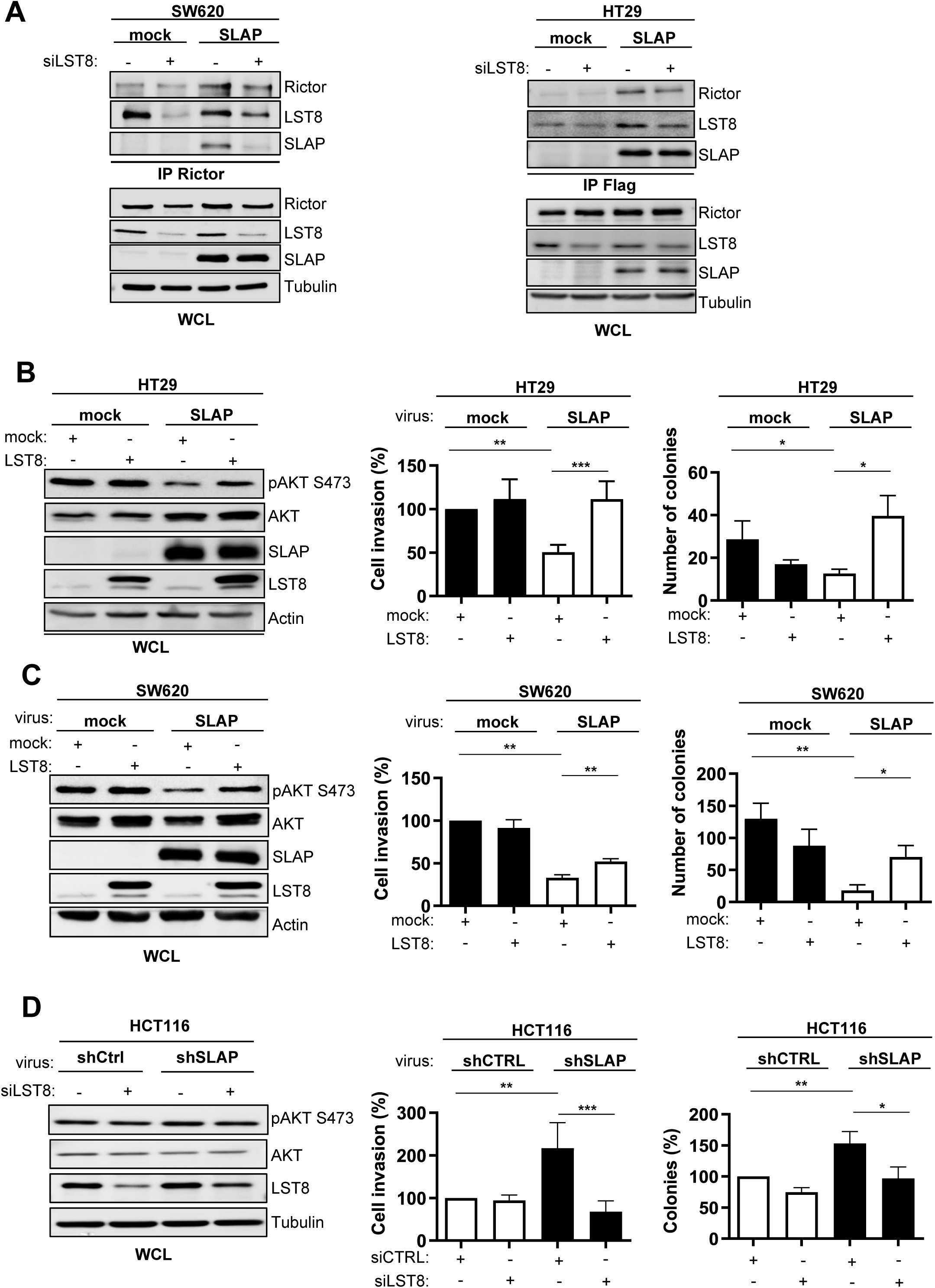
SLAP associates with mLST8 to mediate its anti-oncogenic function. **A.** SLAP-RICTOR interaction is mediated by mLST8 expression in CRC cells. co-immunoprecipitation of SLAP-FLAG with RICTOR in indicated CRC cells that were transfected with a control siRNA or a siRNA targeting LST8 (siLST8) as indicated. The level of SLAP, RICTOR and mLST8 is shown. **B** and **C**: SLAP inhibition of HT29 (**B**) and SW620 (**C**) cell transforming properties are overcome by mLST8 overexpression. Left: WB of indicated protein levels in HT29 cells transduced with indicated retroviruses. Right: cell invasion in Matrigel (control: 100%) and colonies formation (number) in soft agar is shown. **D**. Enhanced HCT116 cell transforming properties induced by SLAP depletion are mediated by mLST8 expression. Left: WB of protein level in indicated HCT116 cells transduced with indicated retroviruses and siRNA. cell invasion in Matrigel and colony formation in soft agar (control: 100%) are shown the mean ± SEM, n=3-5, *p≤0.05, **p≤0.01, ***p≤0.001 (t-test).

Since SLAP acts by promoting the degradation of signaling proteins, we evaluated the effect of SLAP on mLST8 protein stability. However, we did not observe any significant effect of SLAP on mLST8 levels, nor on additional mTORC2 components (Fig. S6A). Inhibition of protein synthesis by cycloheximide treatment of SW620 cells did not show any SLAP effect on mLST8 protein turnover (Fig S6B). Thus, these data do not support a degradation mechanism involved in the regulation of mTORC2 by SLAP. We next examined the effect of SLAP on the integrity of the mTORC2 complex, which is required for activity. Although mLST8 is a common component of mTORC1 and 2, mLST8 is not required for mTORC1 activity, whereas it plays a key role in mTORC2 assembly^3^. Co-immunoprecipitation assays revealed that SLAP expression decreased mLST8 association with RICTOR in SW620 cells, whereas SLAP depletion in HCT116 cells increased this interaction (Fig. 4A). Similar results were obtained for mTOR-RICTOR association (a readout of mTORC2 assembly) in SW620 cells by proximity ligation assay (PLA), in which protein pairs separated by a distance of less than approximately 40 nm were detected using fluorescent probes (Fig 4B).

**Fig. 4:**
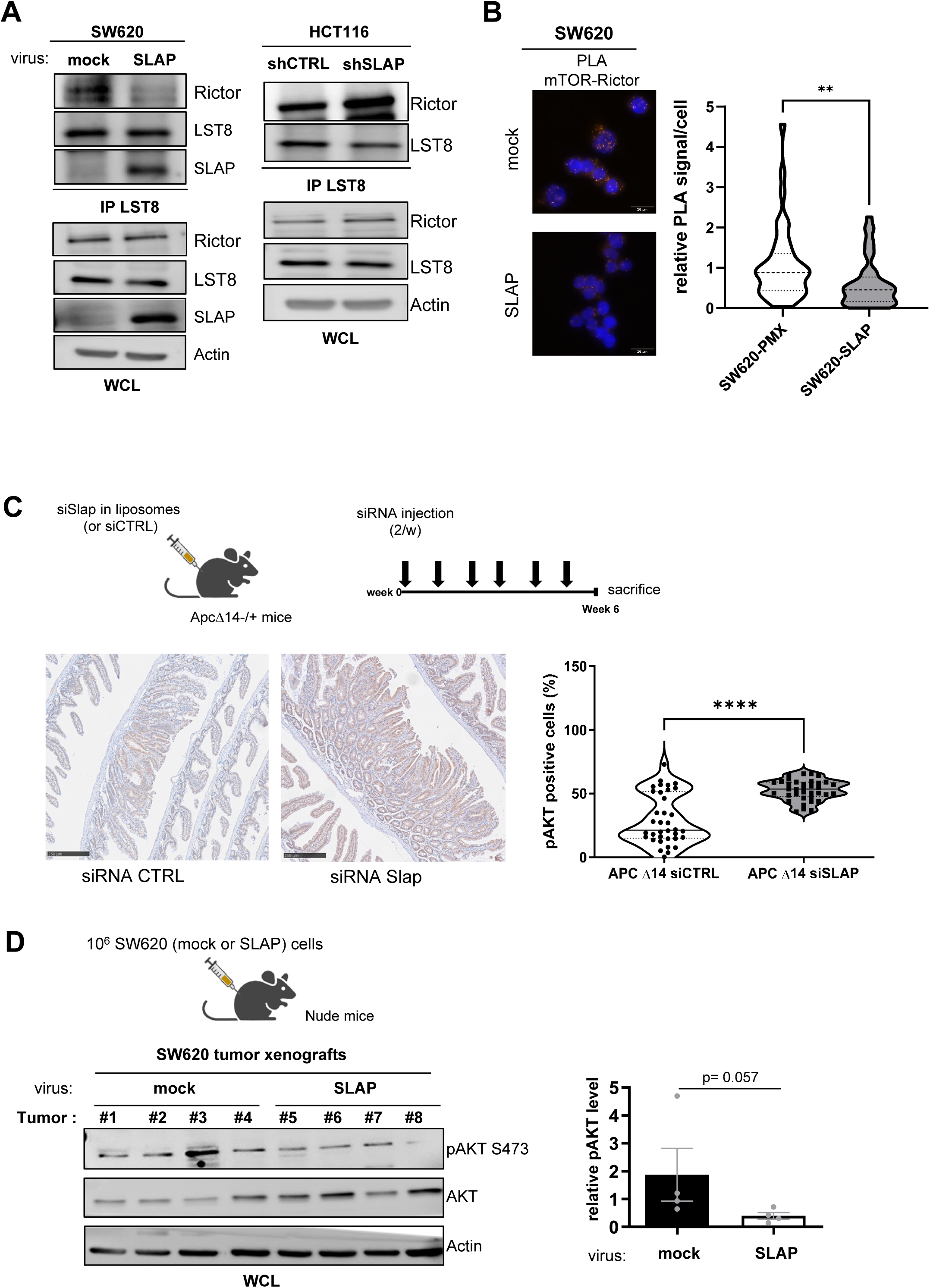
SLAP controls mTORC2 activity by regulating complex integrity. **A.** mLST8-RICTOR interaction, necessary for mTORC2 activity, is controlled by SLAP expression in CRC cells. Association of mLST8 with RICTOR as assessed by co-immunoprecipitation from indicated CRC cells that were transduced with indicated retrovirus. The level of SLAP, RICTOR and mLST8 is shown. The level of pS473 AKT is also assessed as a readout of mTORC2 activity. **B.** The effect of SLAP expression on mTORC2 assembly in SW620 cells. A representative example (left) and its quantification (right) of endogenous mTOR-RICTOR interaction in SW620 cells expressing or not SLAP by PLA. Is shown a violin representation of deduced average PLA signal (20-80 cells per field, 7-15 fields analyzed/replicate, n=4); **p<0.01 (Mann Whitney test). **C.** SLAP silencing increase mTORC2 activity in murine intestinal adenoma. Top: schematic of siRNA-mediated Slap silencing in ApcΔ14−/+ mice. Bottom: a representative example (left) and quantification (right) of pS473 Akt positive cells in intestinal adenoma of mice treated with indicated siRNA. Is shown the mean ± SEM of 5-7 area/mice, n=5 mice per group; ***p<0.001 Mann Whitney test. **D.** SLAP controls mTORC2 activity in SW620 tumor xenografts in nude mice. Top: schematic of CRC tumor xenograft development; (bottom): WB analysis (left) and the quantification (right) of pS473 AKT levels (pAKT/AKT level) in indicated tumors (mean ± SEM, n=4, t-test, p is indicated).

We next investigated the effect of SLAP expression on mTORC2 signaling *in vivo*, by focusing on pS473 AKT level as a readout of mTORC2 activity in experimental CRC models. In *Apc^Δ^*^14^*^/+^* transgenic mice, which develop Wnt pathway-driven intestinal tumors, we previously reported that siRNA-mediated inhibition of *Slap* expression *in vivo* increased the number and size of tumors ^29^. Interestingly, immunohistochemical (IHC) analysis revealed increased pS473 AKT levels in intestinal adenoma of mice treated with siRNA against Slap (Fig 4C). Finally, in human CRC xenografts in nude mice, we reported that SLAP overexpression reduce tumor development in SW620 CRC tumors^29^. Western blotting analysis from these tumor lysates revealed a reduction of pS473 AKT tumor levels by SLAP overexpression (Fig. 4D). Taken together, these data indicate that SLAP inhibits mTORC2 activity by promoting complex disassembly *via* its interaction with mLST8.

### A SLAP-UBE3C complex induces mLST8 ubiquitination and mTORC2 disassembly

To address the underlying mechanism of mTORC2 disassembly, we turned to an mLST8 ubiquitination mechanism activated by SLAP. mLST8 expression in HT29 cells already displays a high level of ubiquitination, while SLAP expression further induced a two-fold increase (Fig. 5A). Similar results were obtained in HEK293T cells (Fig 5A, right panel). We took advantage of this cell system to dissect this ubiquitination process activated by SLAP. First, we found that K/R mutation of Ubiquitin (Ubi-His-Myc) on K29 and K33 reduced the SLAP-mediated Ubiquitination effect on mLST8, in contrast to K48 and K63 mutations (Fig S7). This data indicates that SLAP preferentially induces mLST8 ubiquitination branching on K29 and K33, consistent with a non-degradative mechanism of LST8 ubiquitination (Fig. S7). By mutagenesis analysis (K to A mutation) of mLST8 ubiquitination sites retrieved from the proteomic database (phosphosite.org), we found that SLAP-induced mLST8 ubiquitination involves K86, and possibly K215 and K245, where the triple mutant (mLST8 K86A/K215A/K245A) was poorly ubiquitinated in this assay (Fig. 5B). These data are consistent with a non-degradative ubiquitination mechanism of SLAP-mediated mTORC2 disassembly in CRC cells.

**Fig. 5:**
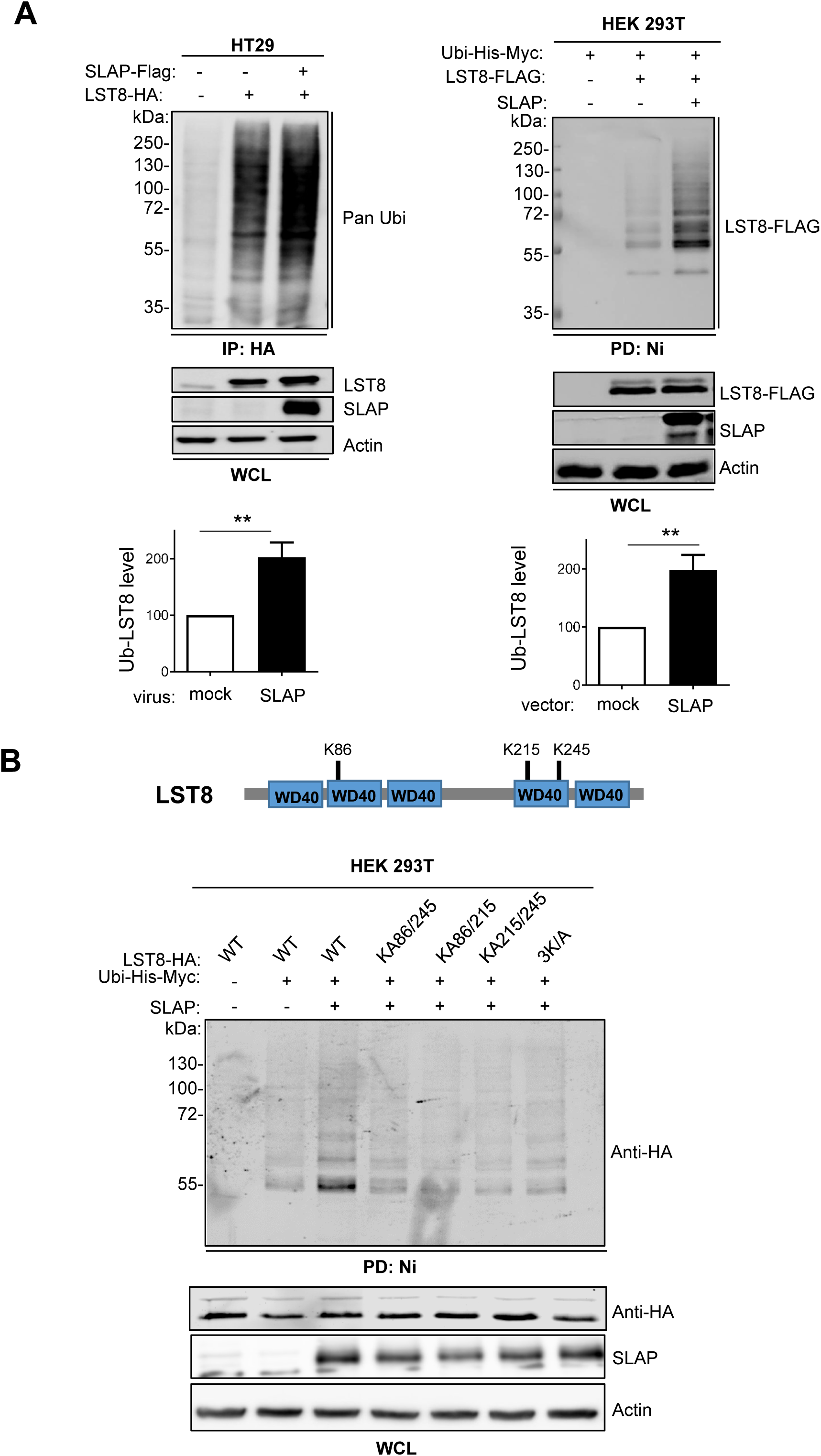
SLAP regulates mTORC2 assembly *via* LST8 ubiquitination. **A**. SLAP expression increases HA-LST8 ubiquitination in HEK293T and HT29 cells. WB analysis (top) and the quantification (bottom) of HA-mLST8 ubiquitination from cells overexpressing or not SLAP, as shown. Is shown the mean ± SEM, n=3-5, **p≤0.01, (t-test). **B**. Mutagenesis analysis of SLAP-dependent HA-mLST8 ubiquitination in HEK293T cells. Is shown the schematic (top) and the mutagenesis analysis (bottom) of SLAP-dependent mLST8 ubiquitination sites as assessed by WB analysis (representative example of 3 independent experiments).

Next, we searched for ubiquitination factors involved in this SLAP signaling mechanism. Since TRAF2 was originally reported to be involved in mTORC2 disassembly^16^, we tested whether this ubiquitination factor was involved in this SLAP effect. However, in contrast to mLST8, we did not detect any association between TRAF2 and SLAP in SW620 and HT29 cells (Fig. S8A), consistent with our proteomic analysis, which also failed to detect TRAF2 as a SLAP interactor (Table S1). Functionally, we did not observe a clear rescue effect of siRNA-mediated TRAF2 depletion on SLAP-induced SW620 cell invasion (Fig. S8B and S8C). These data are inconsistent with a major role for TRAF2 in SLAP signaling in CRC cells. In addition to mTORC2, our interactomic analysis uncovered several novel ubiquitination factors as SLAP interactors in SW620 cells (Fig. S1 and Table S1), including LTN1, UBE3C, UBR5, TRIM25 and possibly TRIM32 (retrieved in 1/3 experiments). We focused on UBE3C, because of its reported capacity to induce K29 and K33 ubiquitination branches^36–38^. We found that siRNA-mediated depletion of UBE3C specifically reversed SLAP-induced mLST8 ubiquitination in HT29 cells, while it had no significant inhibitory effect at the basal level (Fig. 6A). In contrast, depletion of TRIM32, which reportedly induce K48 and K63 ubiquitination^39,40^, had no such inhibitory effect on this SLAP response (Fig. 6A). Thus, these data implicate UBE3C as an E3 ligase involved in this SLAP ubiquitination process on mLST8. Furthermore, UBE3C was associated with SLAP in HT29 and SW620 cells (Fig. 6B). SLAP expression also increased the association between UBE3C and mLST8 in HT29 cells (Fig. 6C). Consequently, UBE3C silencing enhanced mTORC2 assembly, which was inhibited by SLAP expression, as shown by mLST8-RICTOR association, and activity, as shown by pS473 AKT level, in SW620 cells (Fig. 6D). Functionally, siRNA-mediated UBE3C depletion specifically rescued the SW620 cell invasion and anchorage-independent cell growth, which had been inhibited by SLAP (Fig. 6E). In contrast, TRIM32 silencing has no such effect, demonstrating UBE3C specificity (Fig. S9B). Collectively, these data reveal a SLAP-UBE3C ubiquitination mechanism in the control of mTORC2 assembly in CRC cells.

**Fig. 6:**
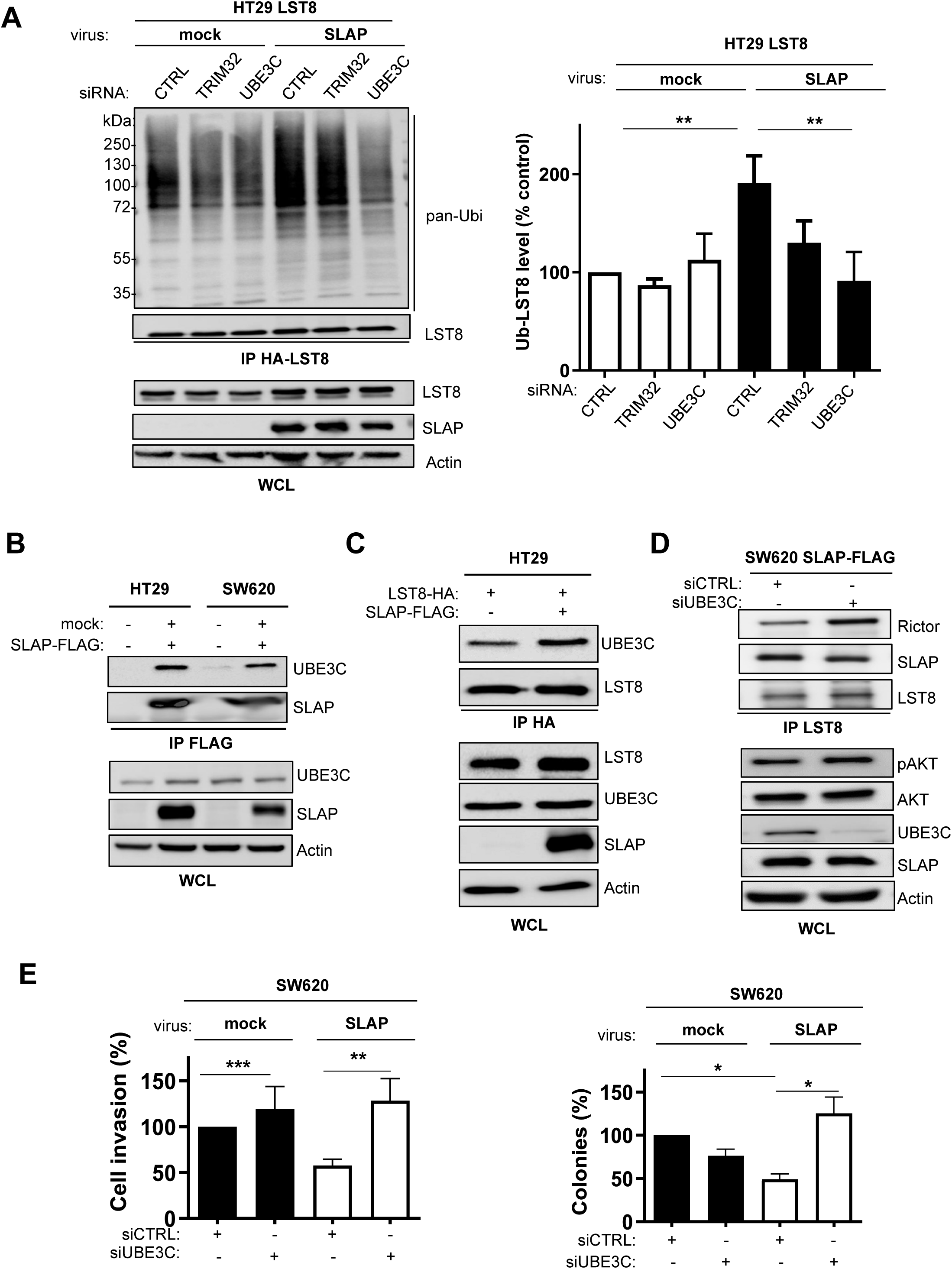
SLAP interacts with UBE3C to facilitate LST8 ubiquitination and mTORC2 inhibition. **A.** SLAP-dependent HA-mLST8 ubiquitination in HT29 cells is dependent upon UBE3C expression. Is shown the ubiquitination level of purified HA-mLST8 from lysates of HT29 cells expressing SLAP or not (control) and transfected with indicated siRNA targeting SLAP-associated ubiquitination factors. Left: a representative example; right: quantification (control: 100%); mean ± SEM, n=3-5, **p≤0.01 (t-test). **B.** SLAP association with UBE3C in HT29 and SW620 CRC cells. **C.** SLAP expression enhances HA-mLST8-UBE3C association in HT29 cells. **D**. UBE3C depletion enhances mLST8-RICTOR association in SLAP overexpressing HT29 cells. **D.** WB analysis of the level of SLAP, UBE3C, and pS473 AKT, as a readout of mTORC2 activity. **E.** SLAP inhibition of HT29 cell transforming properties is overcome by UBE3C depletion. Colony formation in soft agar and cell invasion in Matrigel (control: 100%) is shown. Are shown the mean ± SEM, n=3-5, *p≤0.05, **p≤0.01, ***p≤0.001 (t-test).

### SLAP influences CRC cell response to mTOR inhibitors

Having shown that SLAP defines a novel control mechanism of mTORC2 oncogenic function in CRC cells, we investigated whether it also influences CRC cell response to mTORC2 inhibitors. Since there are no specific mTORC2 inhibitors in clinical development, we used the mTOR catalytic inhibitor (mTORCi) KU-0063494, which targets both mTORC1 and mTORC2^41,42^. We also examined the effect of the mTORC1 specific inhibitor Temsilorimus (TS)^43^ to evaluate the contribution of mTORC1 activity in this response. We found that anchorage-independent cell growth and cell invasion of HT29 cells was strongly inhibited by KU-0063494 treatment (100 nM) in contrast to TS (1 μM), indicating the mTORC2-dependent inhibitory effect (Fig. 7A). This mTORCi inhibitory response was not evident upon expression of SLAP, which already inhibits mTORC2 activity (Fig 7A). Note that TS had a paradoxical promoting effect on pS473 AKT level in SLAP-expressing cells (Fig 7A, top panel), which was attributed to a robust increase in mTORC2 activity, likely due to feedback mechanisms between mTORC1 and mTORC2 activity^41,44^. Similar results were observed in SW620 cells, further validating our findings (Fig. S10A and S10B). Conversely, KU-0063494 strongly inhibited the transforming effects induced by SLAP depletion in HCT116 cells, while TS had no effect, which confirms the mTORC2-dependent response (Fig 7B). In contrast, KU-0063494 had no inhibitory effect in control cells (i.e., shCTRL HCT116 cells), likely due to substantial SLAP-dependent mTORC2 inhibition in these cells (Fig. 7B). In fact, these mTOR inhibitors resulted in a paradoxical promoting effects, as exemplified on anchorage-independent cell growth (Fig 7B), and for TS on pS473 AKT levels (Fig 7B), possibly due to feedback mechanisms between mTORC1 and mTORC2 activity^41^. Consistent with these data, we obtained a good correlation between SLAP expression, pS473 AKT levels and the anti-transforming response to KU-0063494 in a panel of 6 CRC cell lines (Fig. S10C). Finally, we investigated whether SLAP downregulation sensitizes CRC to mTORCi *in vivo*. We turned to the mTORC catalytic inhibitor Torkinib because of its reported effective anti-tumor response in immunodeficient mouse CRC xenograft models^45^. We first confirmed that Torkinib, inhibited SLAP-dependent growth of tumoroids derived from HCT116 cells, as an ex-vivo model of tumorigenesis (Fig. S11). Note that similar results were obtained with the mTORCi AZD2014 (Fig. S11), confirming the mTORC-dependent SLAP effect on this *ex-vivo* cancer model. Next, shCTRL or shSLAP HCT116 cells were subcutaneously inoculated into nude mice, followed by daily treatment with Torkinib (or vehicle) when tumors reached a volume of 50 mm^3^. Although Torkinib did not affect tumor development in control cells, it abrogated the increased tumor volume induced by SLAP depletion (Fig. 8A and 8B). This specific anti-tumor effect was associated with a reduction of CRC cell proliferation and increased cell apoptosis (Fig. S12A and S12B). Similar results were obtained on mTORC2 activity, consistent with the SLAP-dependent mTORC2 tumor function that was targeted by Torkinib (Fig. 8C and Fig. S12C). Thus, we conclude that SLAP acts as an mTORC2 signaling inhibitor in CRC cells and influences the tumor cell response to mTOR catalytic inhibitors (Fig S13).

**Fig. 7:**
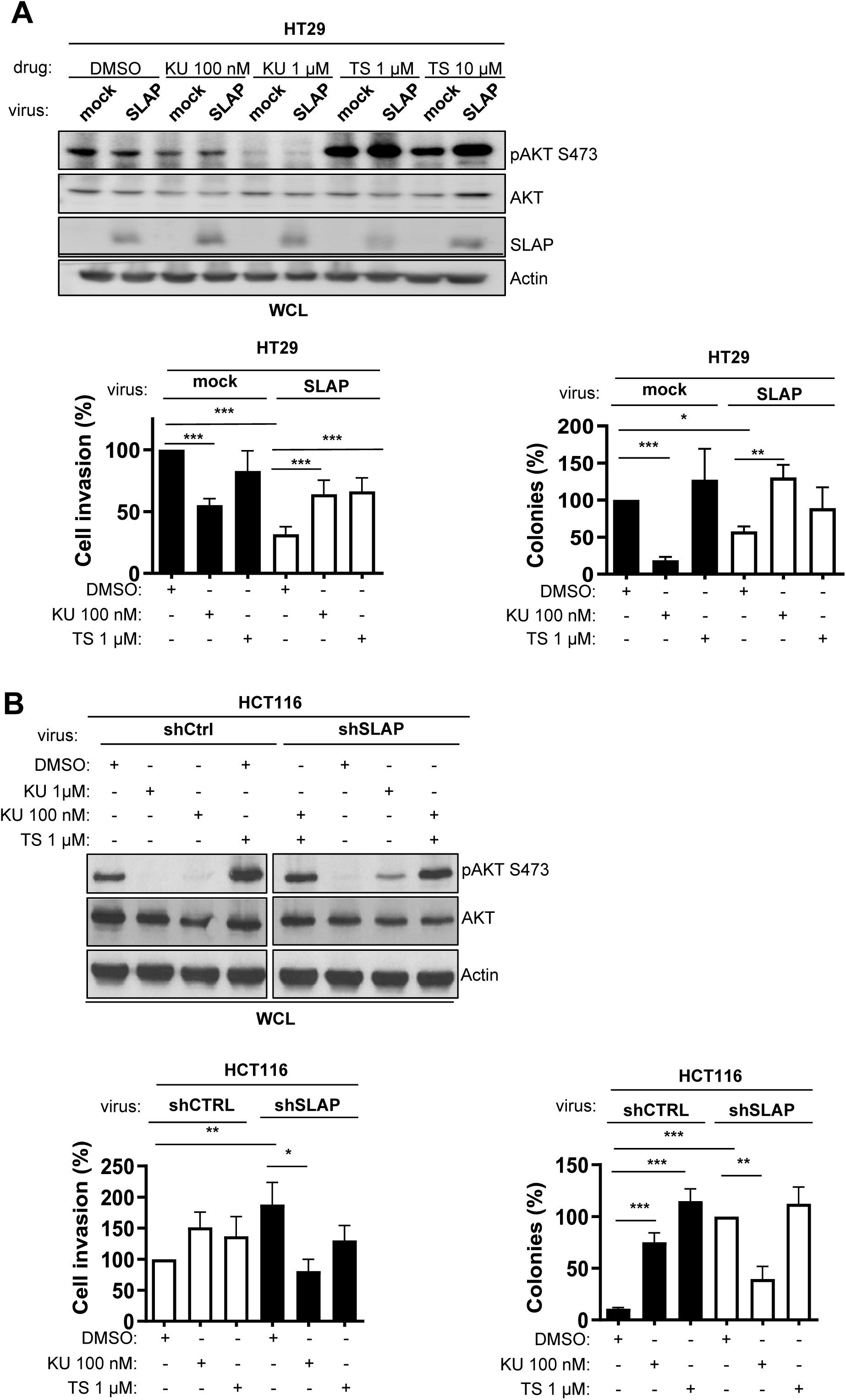
SLAP influences CRC cell response to mTORCi in vitro. **A.** SLAP-dependent inhibition of HT29 cell transforming properties induced by the mTOR catalytic inhibitor KU-0063494 but not the rapamycin analog Temsilorimus. Top: WB analysis of the effect of mTOR inhibitors on pS473 AKT levels in HT29 cells expressing or not SLAP. Bottom: quantification of cell invasion in Matrigel and colony formation in soft agar (control: 100%). **B**. Enhanced HCT116 cell transforming properties induced by SLAP depletion are sensitive to the mTOR catalytic inhibitor KU-006349 but not TS. Top: WB analysis of the effect of mTOR inhibitors on pS473 AKT levels. Bottom: quantification of cell invasion in Matrigel (control: 100%) and colony formation in soft agar (shRNA SLAP: 100%). Are shown the mean ± SEM, n=3-5, *p≤0.05, **p≤0.01, ***p≤0.001 (t-test).

**Fig. 8:**
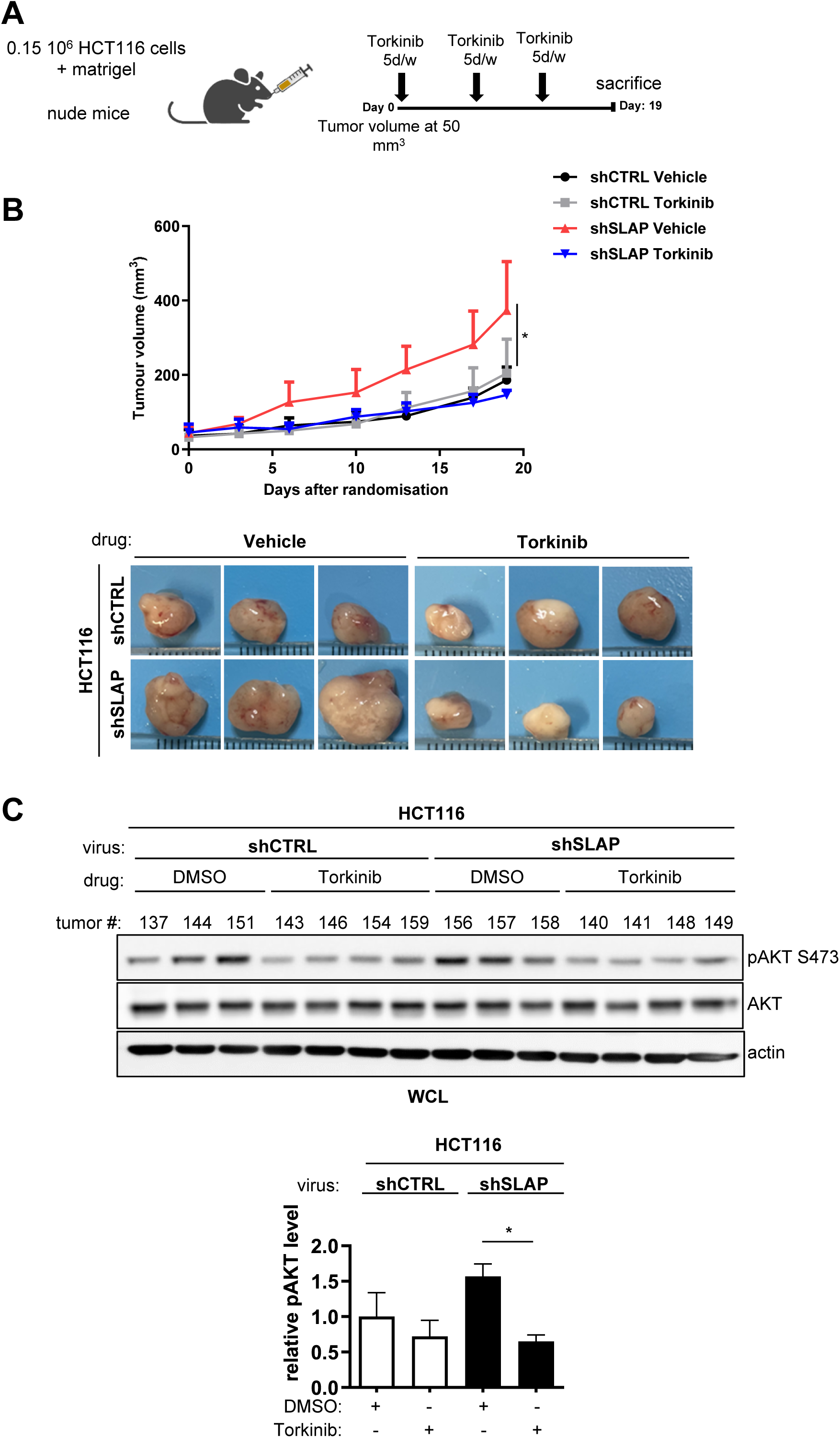
SLAP influences CRC cell response to mTORCi *in vivo*. SLAP-dependent anti-tumor activity of the mTORC catalytic inhibitor Torkinib in CRC xenografts. **A**. Schematic of mice treatment with mTORCi. **B**. Tumor volume induced by control and SLAP-depleted HCT116 cells inoculated into immunodeficient mice and treated daily with Torkinib and a vehicle as control. Top: quantification over time; bottom: representative examples of CRC xenografts. **C.** top: WB analysis of pS473 AKT levels from indicated CRC xenografts. Bottom: quantification. Are shown the mean ± SEM, n=5 mice/cohort, *p≤0.05 (t-test).

## DISCUSSION

Tyrosine kinase signaling receptors are under the control of small adaptor proteins, the prototype of which is suppressors of cytokine signaling protein SOCS, which plays an important role in cancer^46^. SLAP belongs to this family and negatively regulates immune cell receptor signaling through proteasomal degradation of specific signaling components^18,47^. Previously, we reported an unexpected tumor suppressor function of SLAP in the colon and identified the cell adhesion receptor EPHA2 as an important target in this neoplastic process^29^. Our present study demonstrates that SLAP additionally acts as an mTORC2 inhibitor and uses this mechanism to mediate its anti-oncogenic function in CRC. Since EPHA2 has been established as an upstream receptor of AKT signaling^48^, our results show that SLAP inhibits an AKT pathway in CRC by targeting multiple components of the signaling cascade. In fact, EPHA2 has been reported as both an upstream activator and an AKT substrate, creating a feed-forward loop in cancer signaling^49^; therefore, our data support a model in which SLAP inhibits mTORC2 activity both by facilitating complex degradation and by reducing EPHA2/PI3K-dependent mTORC2 activation. However, the mechanistic details of this signaling mechanism are not fully characterized. In particular, while the targeting of EPHA2 is mediated by SLAP-SH2 and the phosphorylation of EPHA2 at Tyr594 by SRC, the targeting of mTORC2 remains unclear because this complex has not been established as a TK substrate. Nevertheless, the SLAP-mTORC2 association implicates SLAP-SH2 in CRC cells. Therefore, it is possible that SLAP targets mTORC2 when in a complex with EPHA2 or another TK signaling receptor to be identified. The exact mechanism of mTORC2 regulation by SLAP requires further investigation.

By enabling mTORC2 assembly induced by SLAP downregulation, our study reveals a novel mechanism of mTORC2 dysregulation in cancer. This finding is highly consistent with previous reports showing mTORC2 oncogenic induction upon mLST8 mutation, which facilitates mTORC2 formation^16,50^. Therefore, dysregulation of complex assembly may be an important mechanism of mTOR oncogenic activation. Mechanistically, ubiquitination of LST8 plays a key role in this molecular process. Our results support a model in which SLAP associates with mTORC2 through a SLAP-SH3-dependent mLST8 interaction to recruit UBE3C and facilitate non-degradative LST8 polyubiquitination. While TRAF2 can promote mTORC2 disassembly *via* K63-linked polyubiquitination of LST8 at K305 and K313^16^, our work suggests that additional UBE3C-dependent polyubiquitination of K86, K215 and K245 is also involved. Interestingly, these residues are located in WD40 domains involved in SIN1-mLST8-RICTOR complex formation^13^, predicting that their polyubiquitination would destabilize these interactions. Consistent with this finding, a recent study reveals a similar ubiquitination mechanism at K86 and K261 mediated by the oxidative sensor KEAP1^50^, revealing the diversity of ubiquitination-dependent mLST8 scaffold inhibition. From this model, several mechanisms of mTORC2 dysregulation in cancer can be proposed, including mLST8 mutations resistant to ubiquitination, as reported in melanoma, or dysregulation of ubiquitination mechanisms involved in this process. Oncogenic mLST8 mutations have been rarely detected in CRC. Rather, mTORC2 overactivation was associated with overexpression of multiple components of the complex, including mTOR and RICTOR^5^. Our results indicate that overexpression of RICTOR or mLST8 is sufficient to bypass SLAP-dependent mTORC2 inhibition. Thus, we propose that the combination of SLAP downregulation with RICTOR overexpression may contribute to the deregulation of mTORC2 assembly in CRC.

Clinically developed mTORCi have shown variable efficacy in CRC^9–12^. The newly reported mechanism of mTORC2 dysregulation by SLAP downregulation may help predict the therapeutic response to these ATP site inhibitors. For example, overactivation of RTK signaling, such as PI3K or RAS pathways, has been reported to influence CRC response to mTORCi^9–12^. Our results add SLAP tumor expression as a potential predictor of CRC response to mTORCi. The fact that SLAP controls CRC metastasis in nude mice models and the cell invasive function of mTORC2 in vitro suggests that SLAP may also regulate the invasive function of mTORC2 *in vivo*. Whether SLAP expression influences mTORCi activity in metastatic CRC, alone or in combination with chemotherapy, deserves further investigation. Similarly, the PIKK co-chaperones TTI were retrieved from our SLAP proteomic analysis in CRC cells. Whether SLAP also affects PIKK assembly and influences CRC response to PIKK inhibitors is another question to be addressed in the future.

Collectively, our data reveal a central mechanism of mTORC2 oncogenic induction in CRC mediated by dysregulation of complex integrity *via* downregulation of the tumor suppressor adaptor SLAP, with potential implications in mTORCi-targeted therapy (Fig. S13). By showing that SLAP mediates its signaling function by recruiting diverse ubiquitination factors (i.e. CBL, UBE4A and UBE3C) and possibly additional ones (i.e. LTN1, UBR5, TRIM25…) as revealed by our proteomic analysis, this study sheds light on a more complex signaling mechanism by which SLAP regulates TK signaling receptors than initially reported. Finally, since SLAP expression is altered in other cancers (e.g., lung cancer, and leukemia), this uncovered SLAP activity may also have important function in these cancers.

## ACKNOWLEDGEMENTS

We acknowledge Montpellier Biocampus facilities: Montpellier Ressources Imagerie (MRI) platform, the histology and animal experimentation platforms RHEM and RAM, the PCEA and IGMM mouse facilities ZEFI, and the Proteomic Core Platform of FPP for proteomic analyses. This work was supported by La Ligue Nationale Contre le Cancer (LNCC), La Foundation pour la Recherche Médicale, Montpellier SIRIC Grant «INCa-DGOS-Inserm 6045», CNRS, and the University of Montpellier. RHEM facility supported by SIRIC Montpellier Cancer Grant INCa_Inserm_DGOS_12553, the European regional development foundation and the Occitanian region (FEDER-FSE 2014-2020 Languedoc Roussillon), la Ligue Contre le Cancer for processing our animal tissues and histology technics. RM and DN were supported by the LNCC, ARC and Montpellier University. SR is an INSERM investigator.

## AUTHORS CONTRIBUTION

All authors contributed extensively to the work presented in this paper. Experimental analysis and Data acquisition: RM, DN, FC, CN, GG, BF, KE, VS, YB, JN and AS. MS analysis: SU. Project supervision: SR and AS. Funding acquisition and Writing of the paper: SR.

## CONFLICT OF INTEREST

The authors declare no conflict of interest.

## Data availability statement

The data that support the findings of this study are available from the corresponding authors upon reasonable request.

## Notes

### Competing Interest Statement

The authors have declared no competing interest.

