## Supplementary figures & table S1 for "SLAP controls mTORC2 integrity *via* UBE3C-mediated mLST8 ubiquitination to mediate its tumour suppressive function in colorectal cancer"

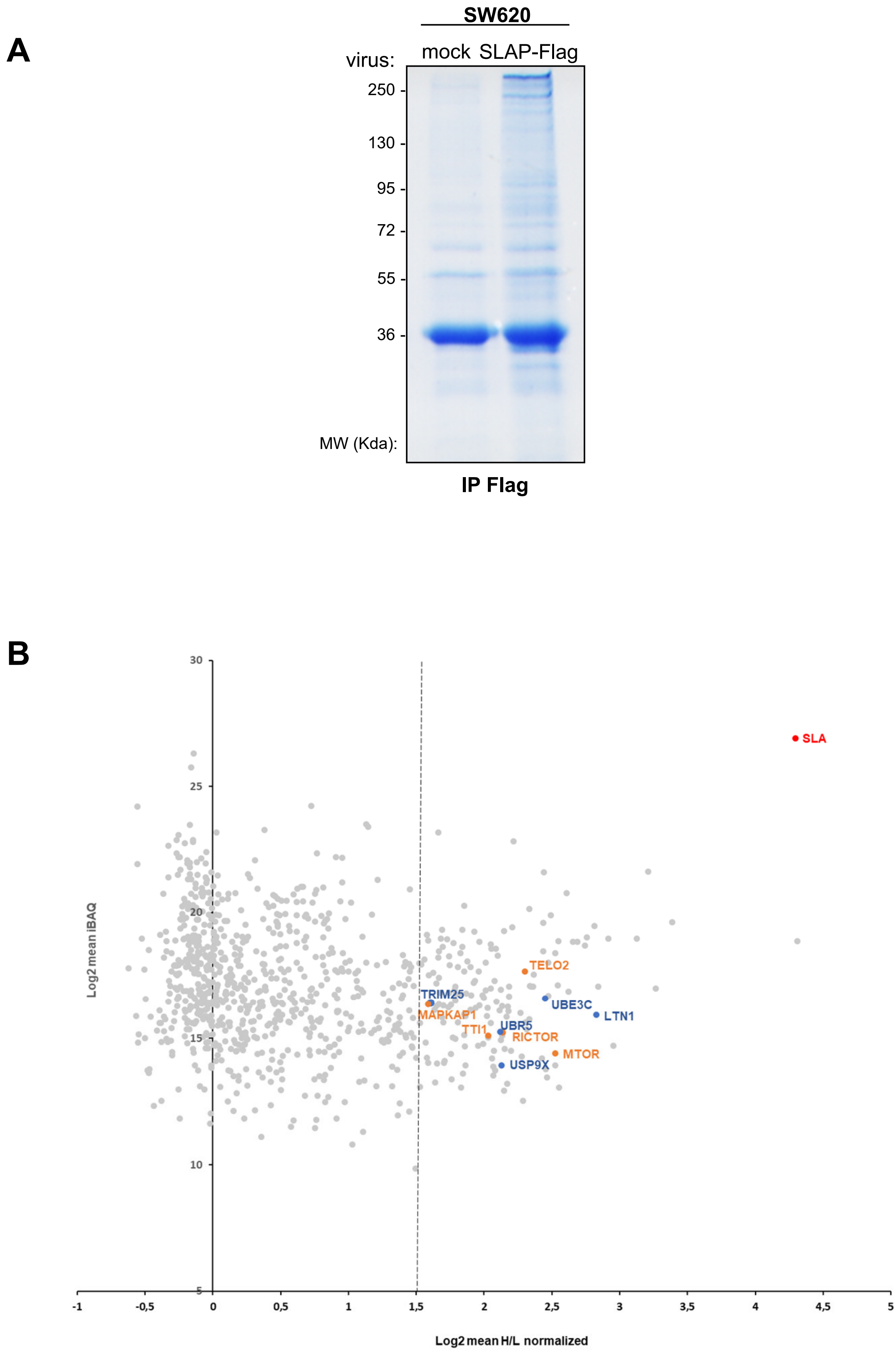


**Figure S1: SILAC-based SLAP interactomics in SW620 cells. A:** brillant blue staining of SDS-PAGE gel from SLAP-Flag immunoprecipitation in SW620 cells. **B:** graph showing the relative quantification (log2 mean intensity relative to the log2 mean H/L normalized ratio) of SLAP interactors (identified ≥2/3 independent experiments with ≥3 peptides). SLAP (red), mTORC2 components (orange) and ubiquitination factors (blue) are highlighted.

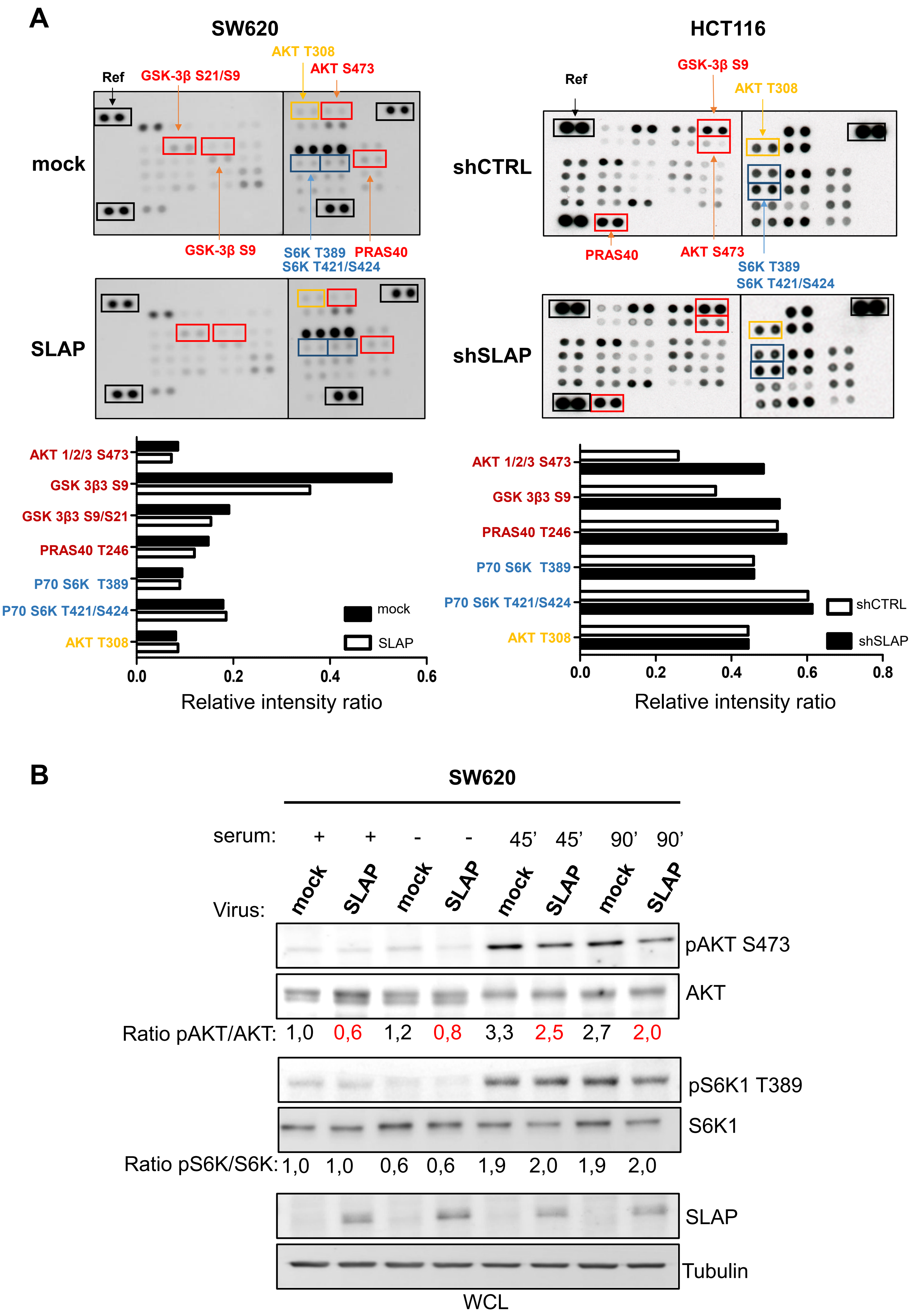
A
SW620
HCT116
Ref
GSK-3β S21/S9
AKT T308
AKT S473
mock
GSK-3β S9
S6K T389
PRAS40
S6K T421/S424
SLAP
GSK-3β S9
AKT T308
shCTRL
PRAS40
AKT S473
S6K T389
S6K T421/S424
shSLAP
AKT 1/2/3 S473
GSK 3β3 S9
GSK 3β3 S9/S21
PRAS40 T246
P70 S6K T389
P70 S6K T421/S424
AKT T308
mock
SLAP
0.0
0.2
0.4
0.6
Relative intensity ratio
AKT 1/2/3 S473
GSK 3β3 S9
PRAS40 T246
P70 S6K T389
P70 S6K T421/S424
AKT T308
shCTRL
shSLAP
0.0
0.2
0.4
0.6
0.8
Relative intensity ratio
B
SW620
serum: + + - - 45' 45' 90' 90'
Virus: mock SLAP mock SLAP mock SLAP mock SLAP
pAKT S473
AKT
Ratio pAKT/AKT: 1,0 0,6 1,2 0,8 3,3 2,5 2,7 2,0
pS6K1 T389
S6K1
Ratio pS6K/S6K: 1,0 1,0 0,6 0,6 1,9 2,0 1,9 2,0
SLAP
Tubulin
WCL

C

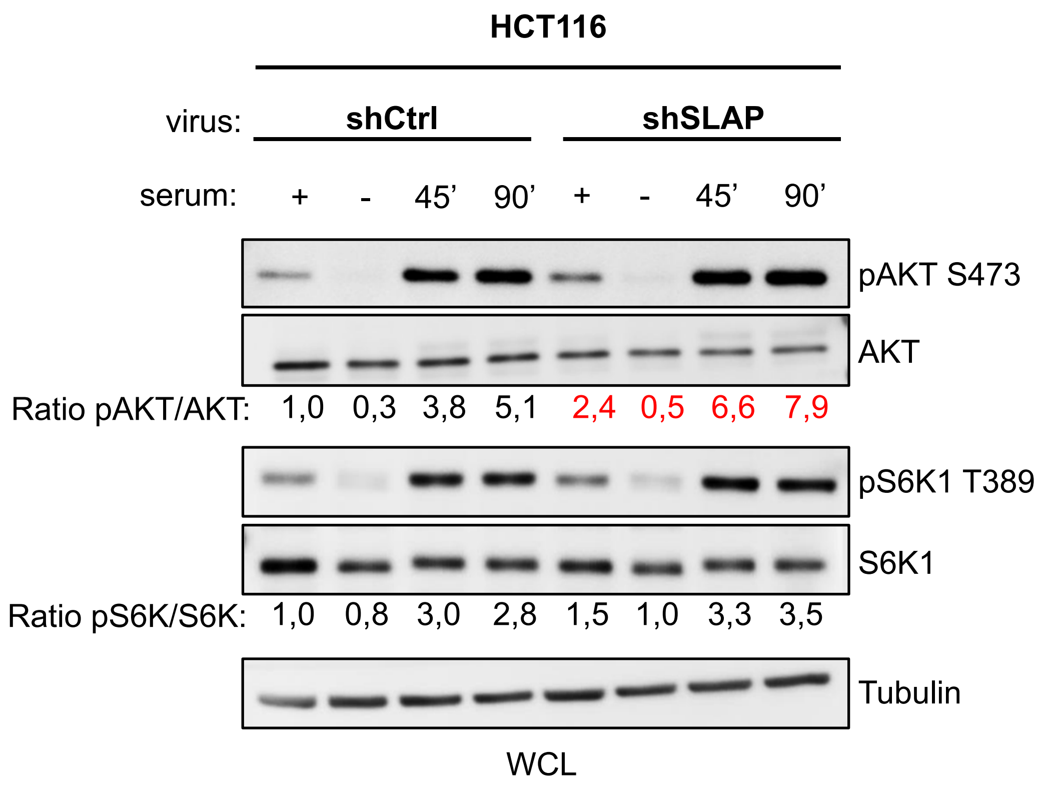


**Figure S2: SLAP regulates mTORC2 phospho-signaling in CRC cells**. **A**: phospho-kinase array of indicated cell-lysates; **B** (SW620) and **C** (HCT116): WB analysis of selected mTOR substrates phosphorylation in quiescent CRC cells expressing or not SLAP that were stimulated with 10% serum at indicated time points. The relative phosphorylation level is shown (ratio).

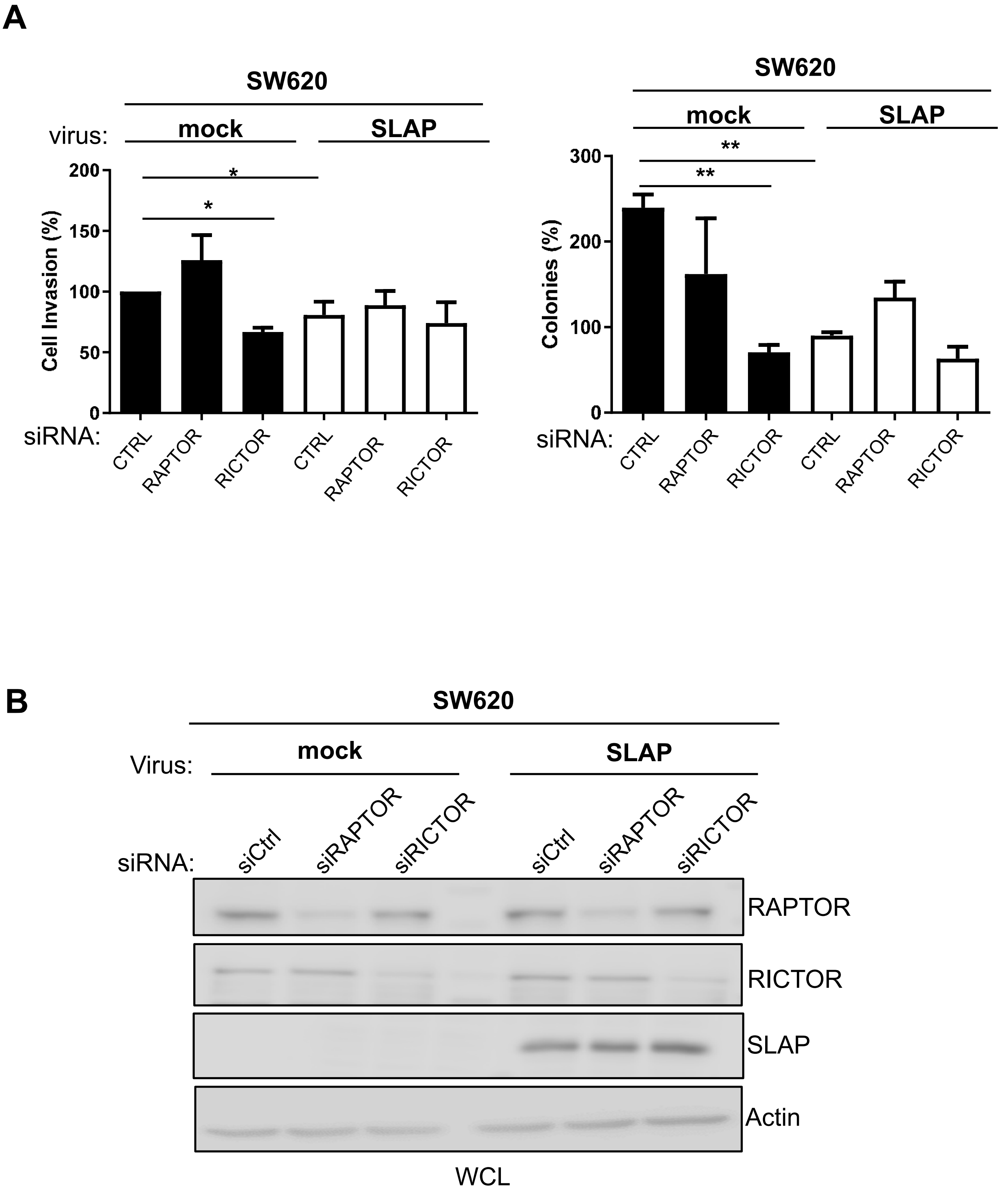


**Figure S3: SLAP-dependent mTORC2 transforming function in CRC cells. A**: RICTOR-dependent anchorage-independent cell growth and invasion in matrigel of SW620 cells expressing or not SLAP. **B** :WB of indicated protein levels. Mean ± SEM, n=4-6; *p<0.05; **p<0.01 Student's t test

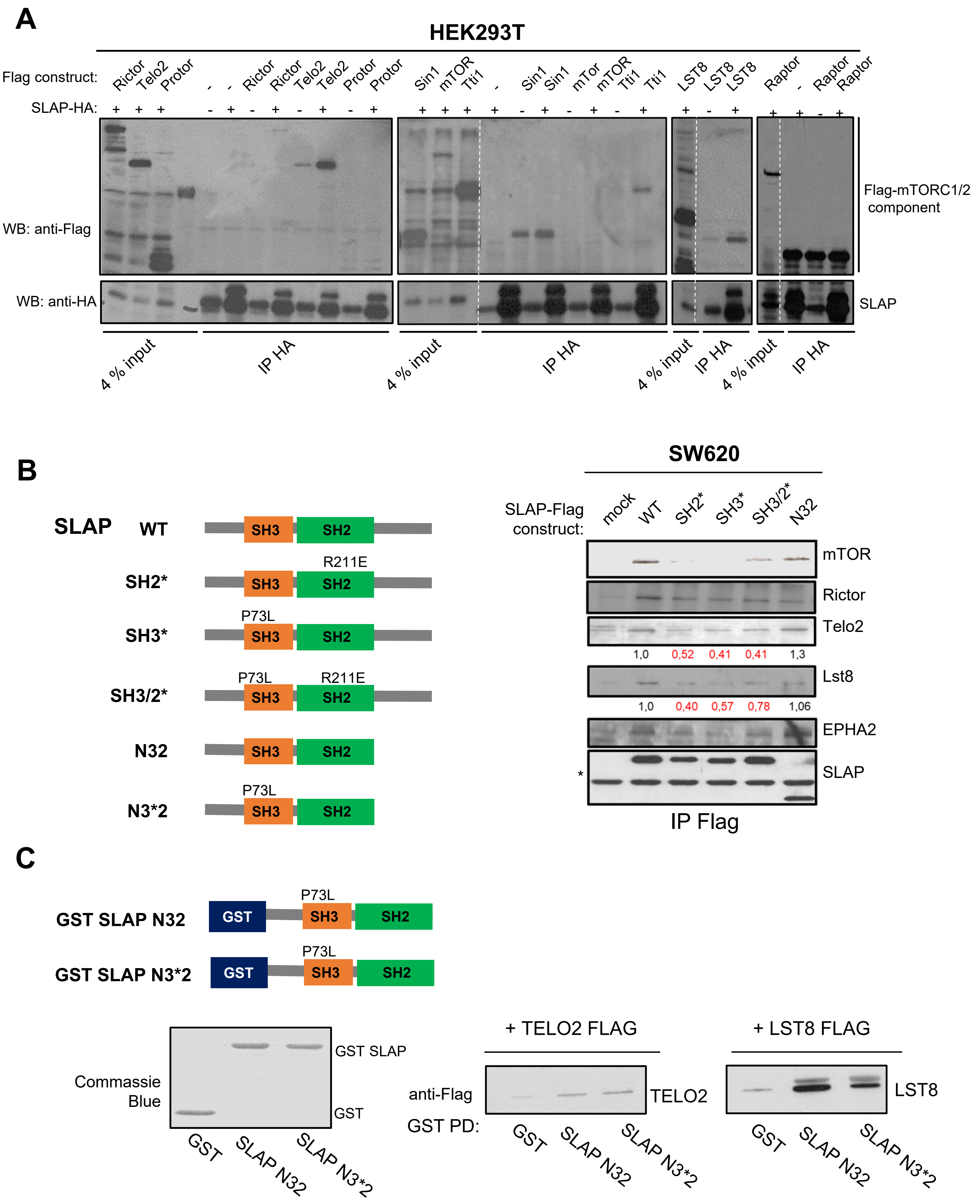


**Figure S4: SLAP interaction with TELO2 and LST8. A**: SLAP co-immunoprecipitation with mTORC1 and 2 components that were expressed in HEK293T cells. **B**: Mutagenesis analysis of SLAP interaction with mTORC2 components in SW620 cells transduced with indicated SLAP-Flag constructs. EPHA2 was used as a positive control **C**: GST-SLAP pull-down (PD) using HEK293T cell-lysates transfected with TELO2 and LST8 constructs as shown. Dotted lines in A indicate where the image was cropped to remove irrelevant lanes. Raw data for the full image are provided in the Source Data file. *: unspecific bans (light chain of ip Flag antibody).

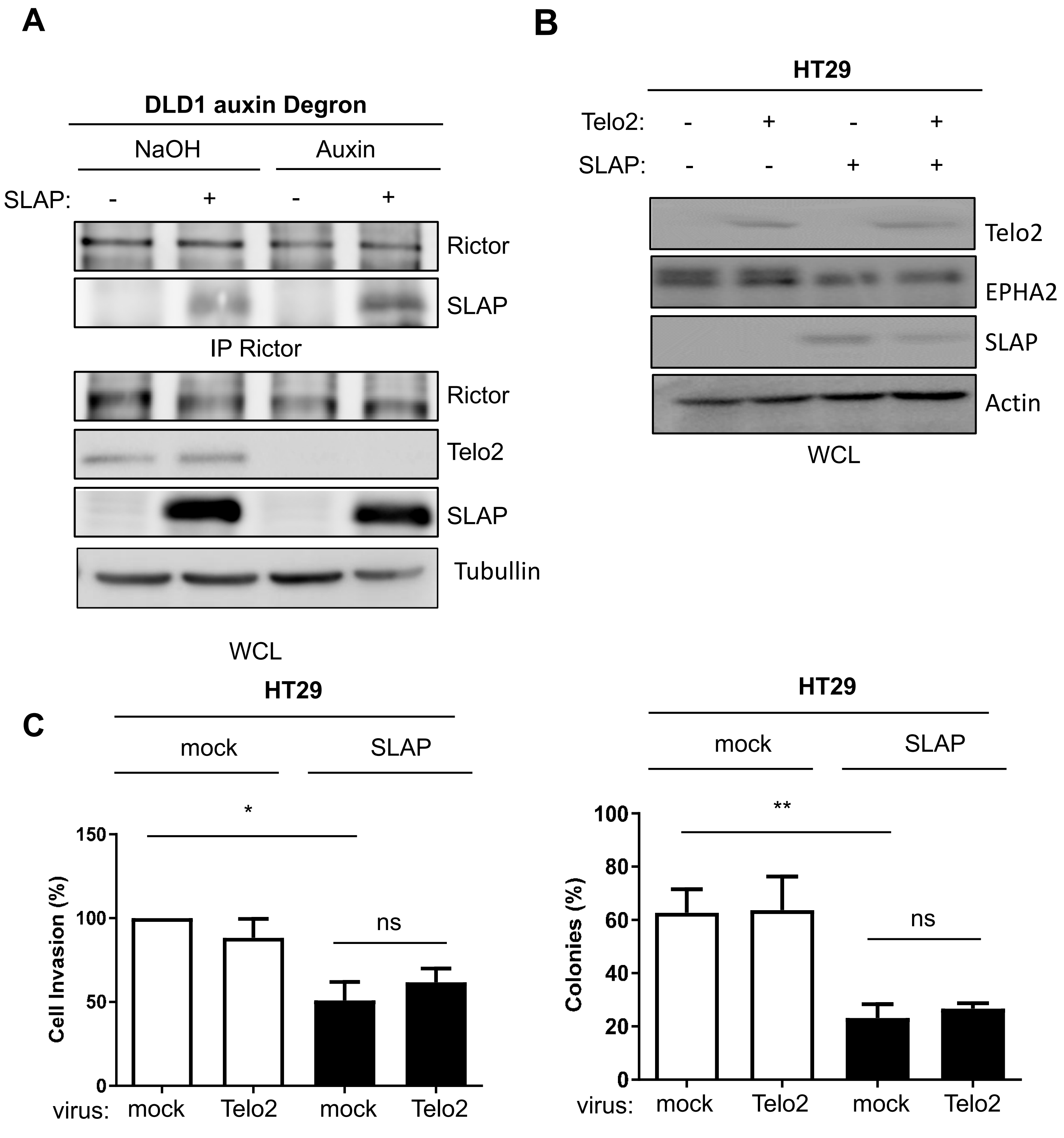


**Figure S5: TELO2 is dispensable for SLAP anti-oncogenic effect. A**: Telo2 degradation via a auxin-induced degron system does not affect SLAP association with RICTOR in DLD1 CRC cells. **B:** WB of Telo2 overexpression in control and SLAP overexpressing HT29 cells. The level of the SLAP target EPHA2 is shown. **C**: Telo2 overexpression does not overcome SLAP anti-oncogenic properties in HT29 cells. The Mean ± SEM, n=3-5; ns p>0.05; *p<0.05; **p<0.01; Student's t test.

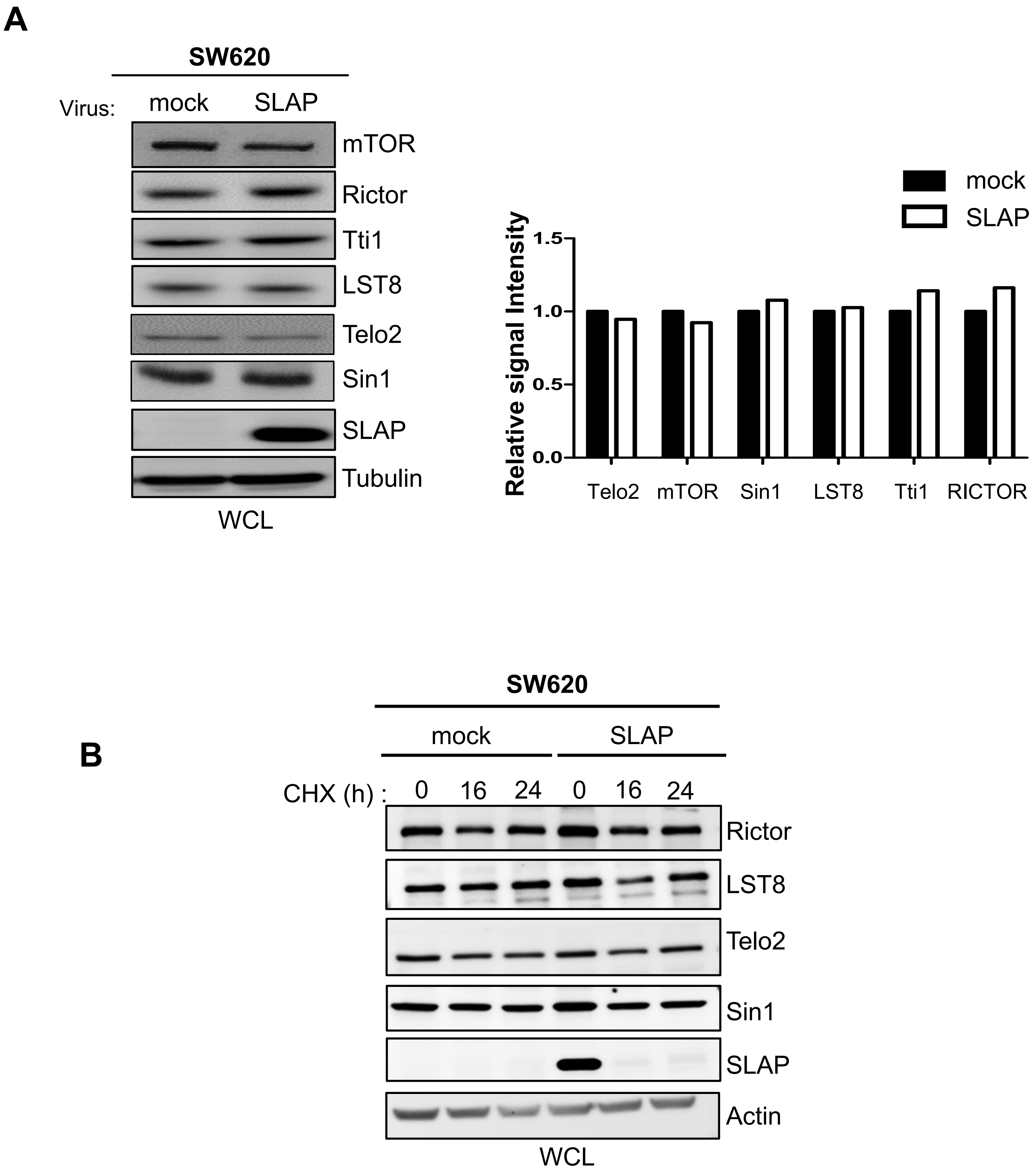


**Figure S6: SLAP does not affect the protein stability of mTORC2 components. A**: SLAP does not affect mTORC2 proteins levels in CRC cells. Is shown a representative example (left) and quantification (mean, n=2) (right). **B:** SLAP does not affect mTORC2 protein turnover, as shown from cell treatment with cycloheximide at indicated times (CHX; 100μg/ml).

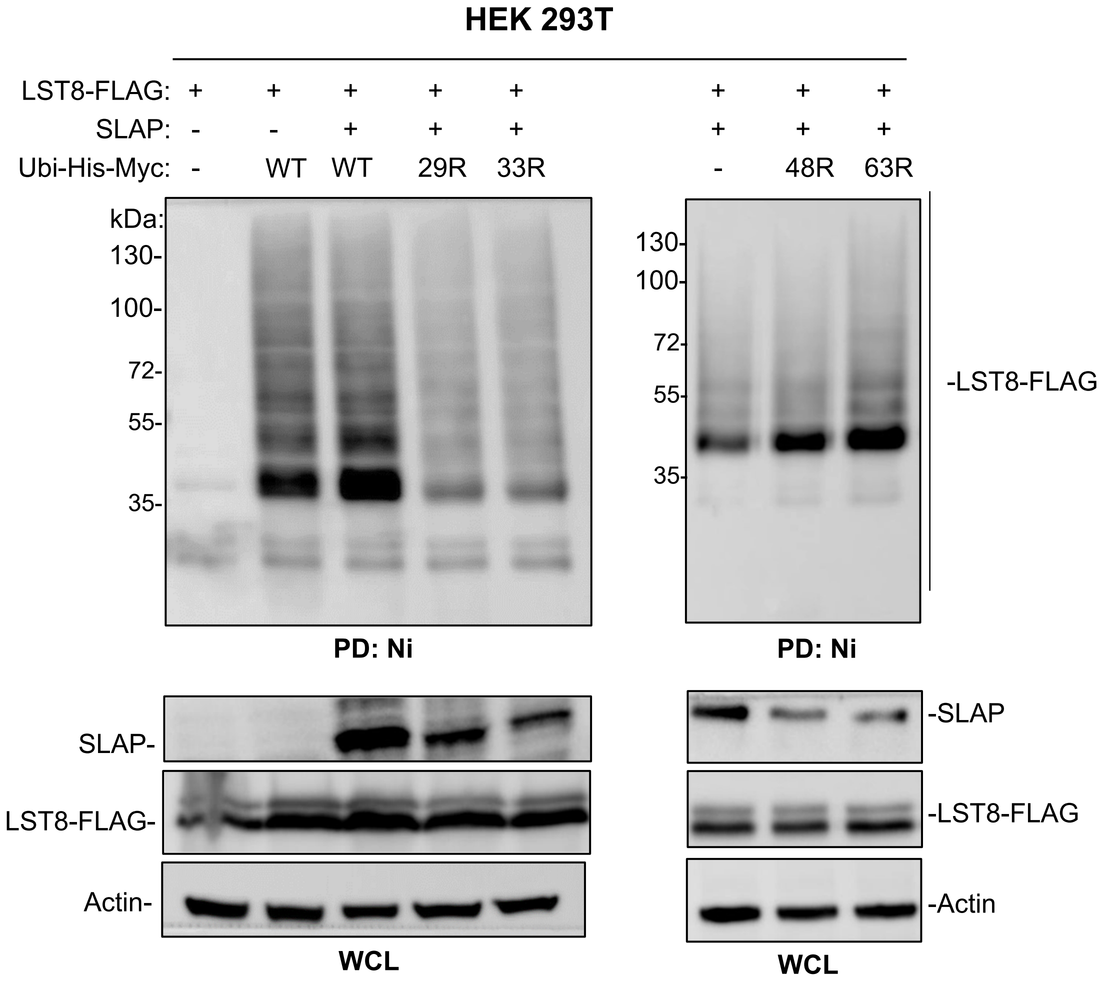


**Figure S7: SLAP-mediated LST8 ubiquitination involves branching on Ubiquitin K29 and K33.** SLAP-dependent LST8 ubiquitination assay in HEK 293T cells using SLAP WT or indicated KR Ubiquitin (Ub-His-Myc) mutants.

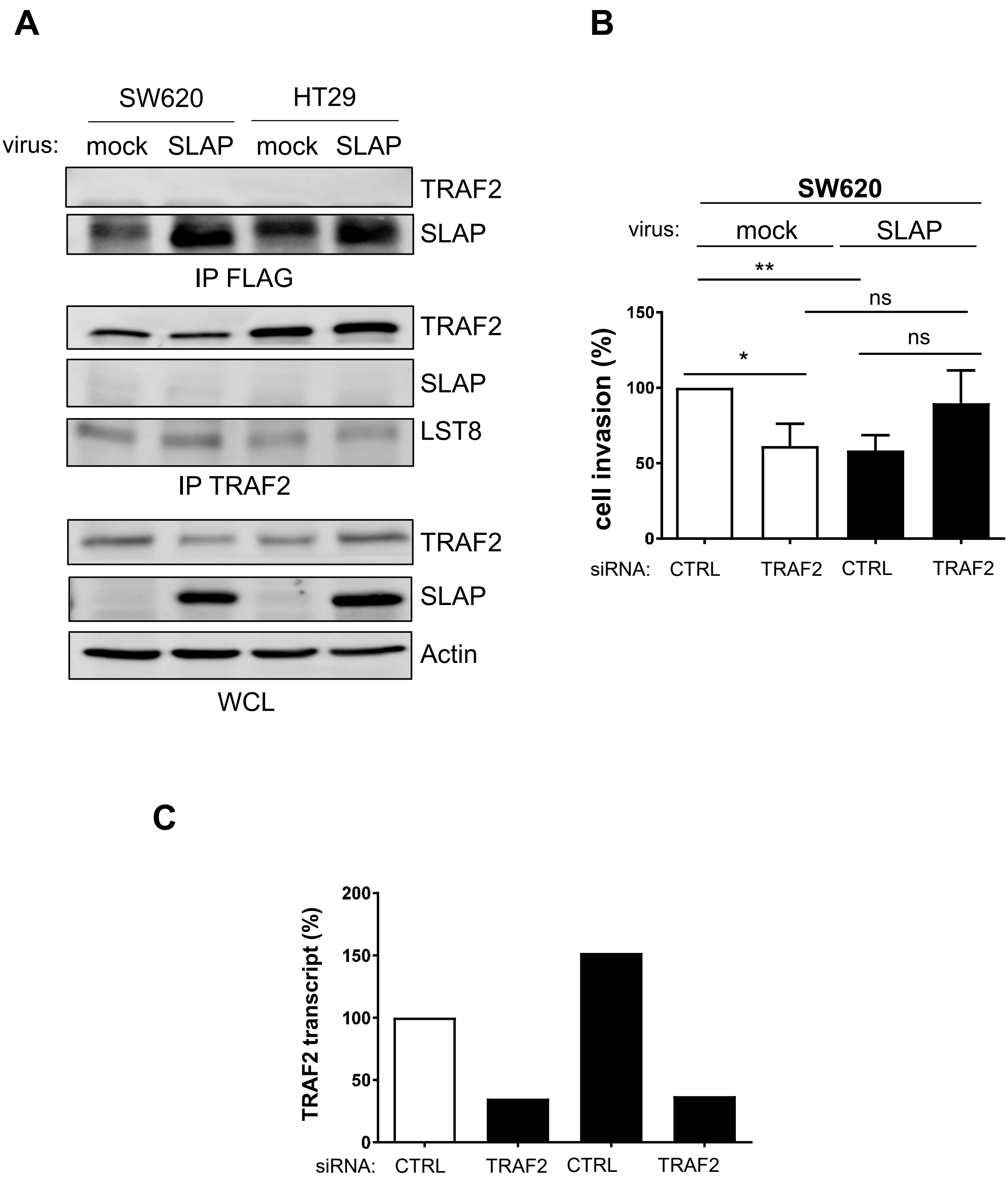


**Figure S8: TRAF2 is not involved in SLAP signaling. A**: SLAP does not interact with TRAF2 in CRC cells. **B.** TRAF2 depletion does not affect SLAP anti-invasive properties in CRC cells. Is shown the mean ± SEM, n=5; ns $p>0.05$; *$p<0.05$, **$p<0.01$; Student's t test. **C.** siRNA silencing of TRAF2 expression by qPCR (n=2).

**A**

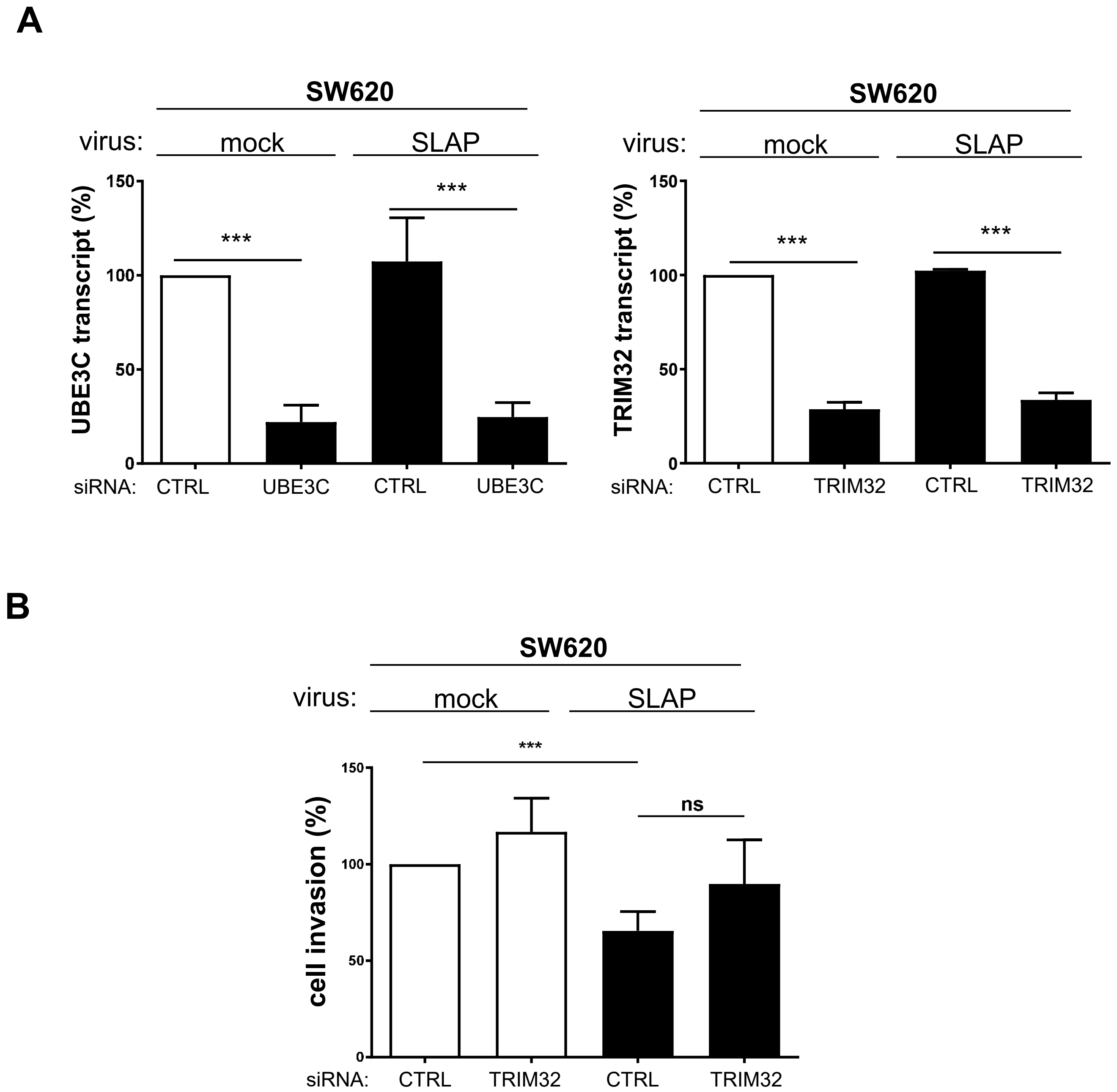


**Figure S9: UBE3C is implicated in SLAP signaling.** **A** qPCR analysis of siRNA UBE3C (left) and TRIM32 (right) silencing (n=3). **B**: TRIM32 depletion does not significantly affect SLAP anti-invasive properties (mean ± SEM, n=5); ns p>0.05; ***p<0.001; Student's t test.

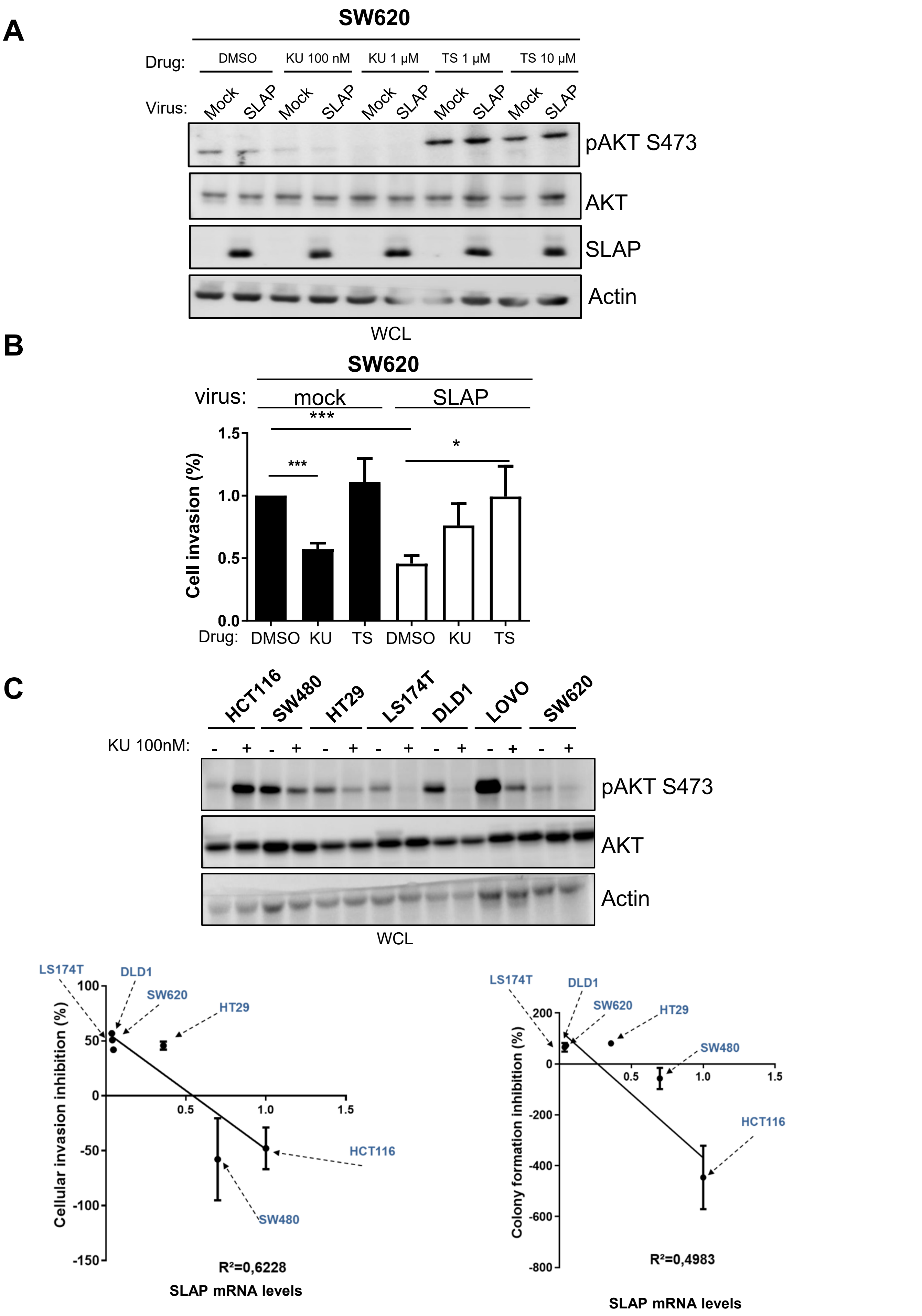


**Figure S10: SLAP overexpression reduces mTORCi activity in SW620 cells.** pAKT level (**A**) and cell invasion (**B**) of cells treated with indicated mTORC inhibitors. **C**: Correlation between endogenous SLAP expression and CRC cell responses to mTORCi (mean ± SEM, n=5; ns p>0.05; ***p<0.001; t-test).

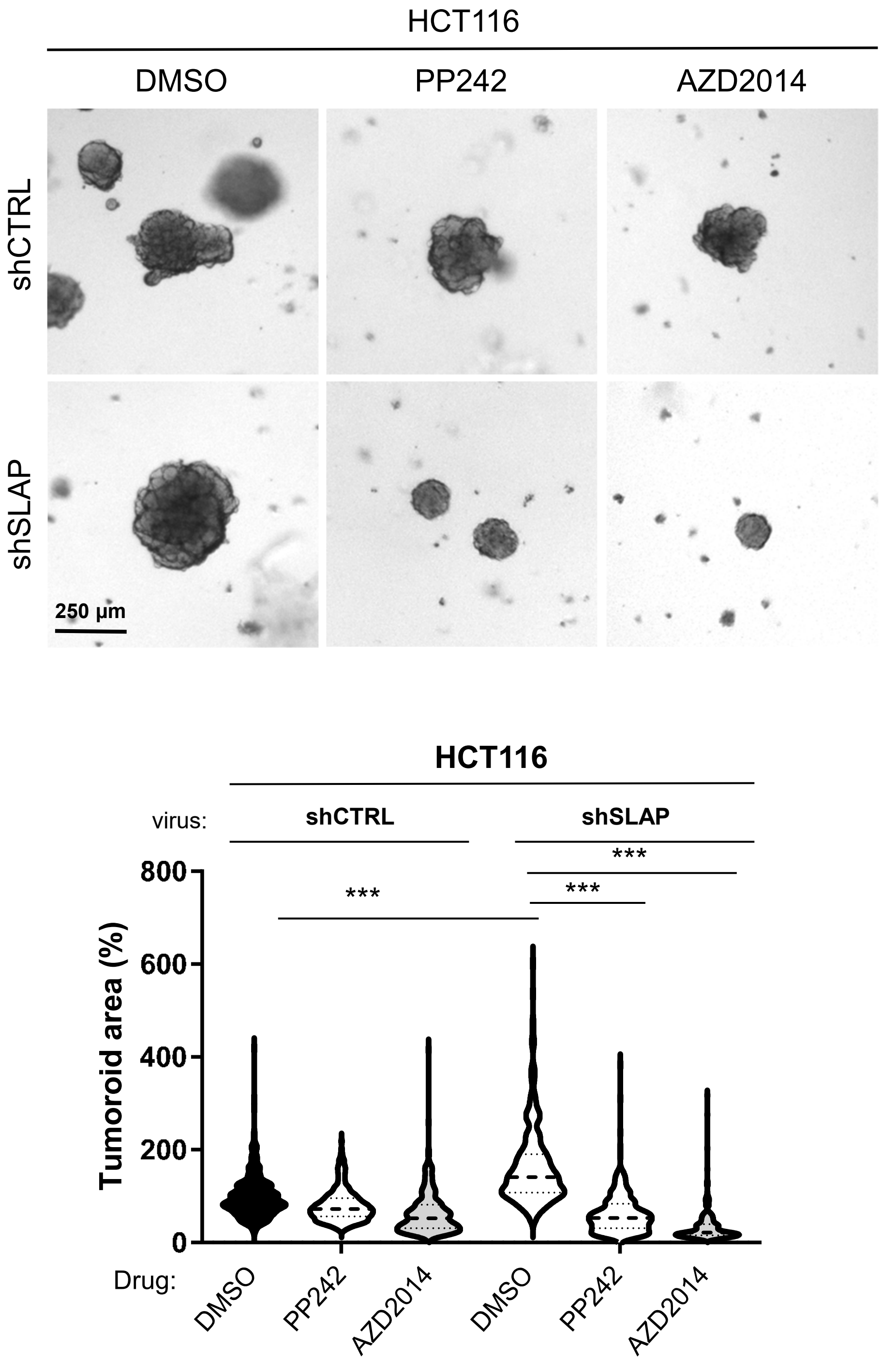


**Figure S11: SLAP depletion enhances the mTORCi activity in tumoroids derived from HCT116 cells.** A representative example (top) and its quantification (bottom) of indicated mTORCi (PP242/Torkinib: AZD2014) effects on tumoroid growth. Is shown a violin representation of the mean, 80-100 tumoroids analyzed/replicate, n=3; ***p<0.001; Student's t test.

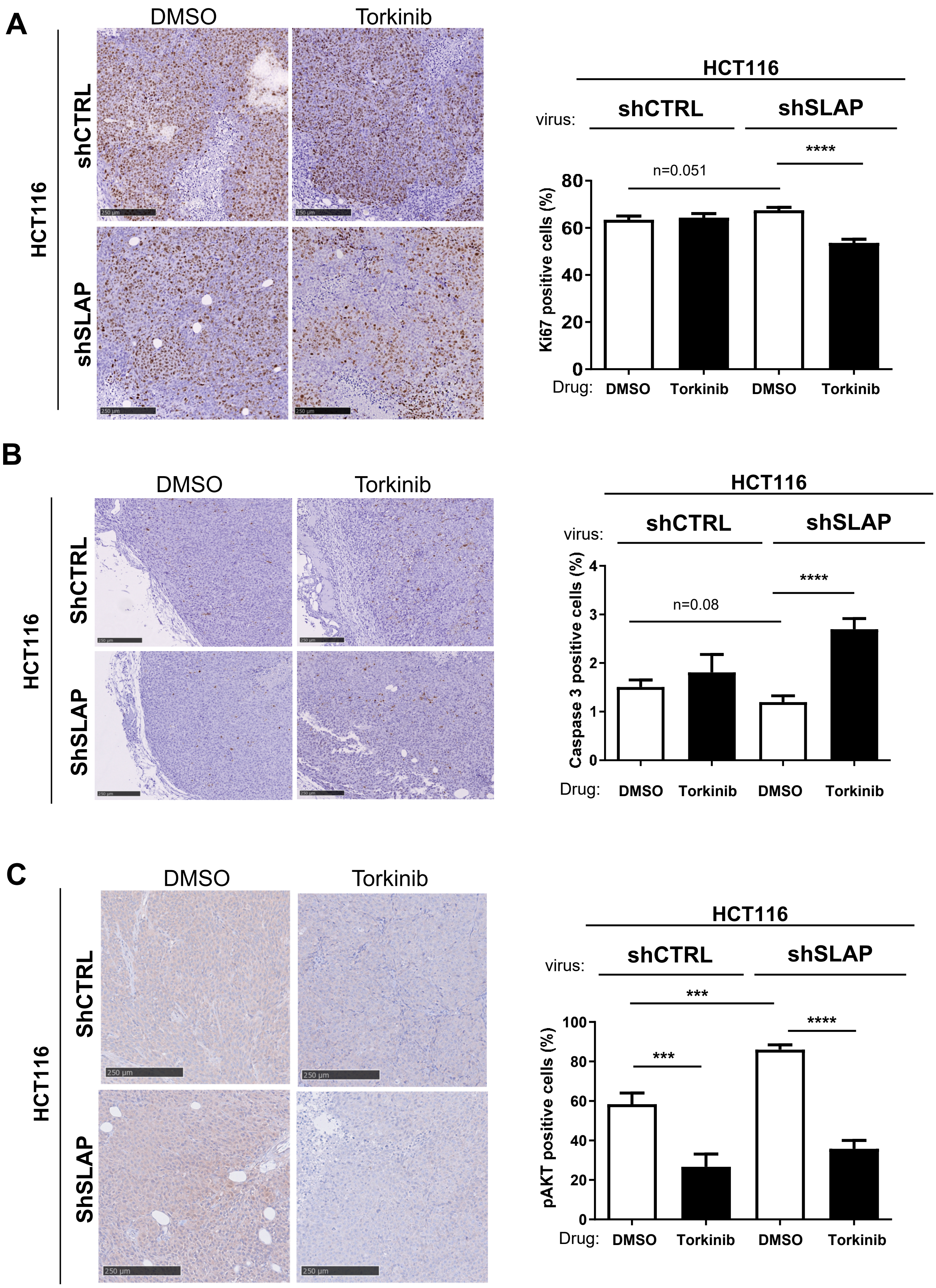


**Figure S12: SLAP depletion sensistizes CRC cells reponse to mTORCi *in vivo*.** IHC analysis of (**A**) cell proliferation (Ki67), (**B**) apoptosis (Caspase 3) and (**C**) pS473 AKT level in indicated tumors treated or not with Torkinib (mean ± SEM, n=5 mice; p close to 0.05 is indicated; ***p<0.005, ****p<0.001; Mann-Withney t-test).

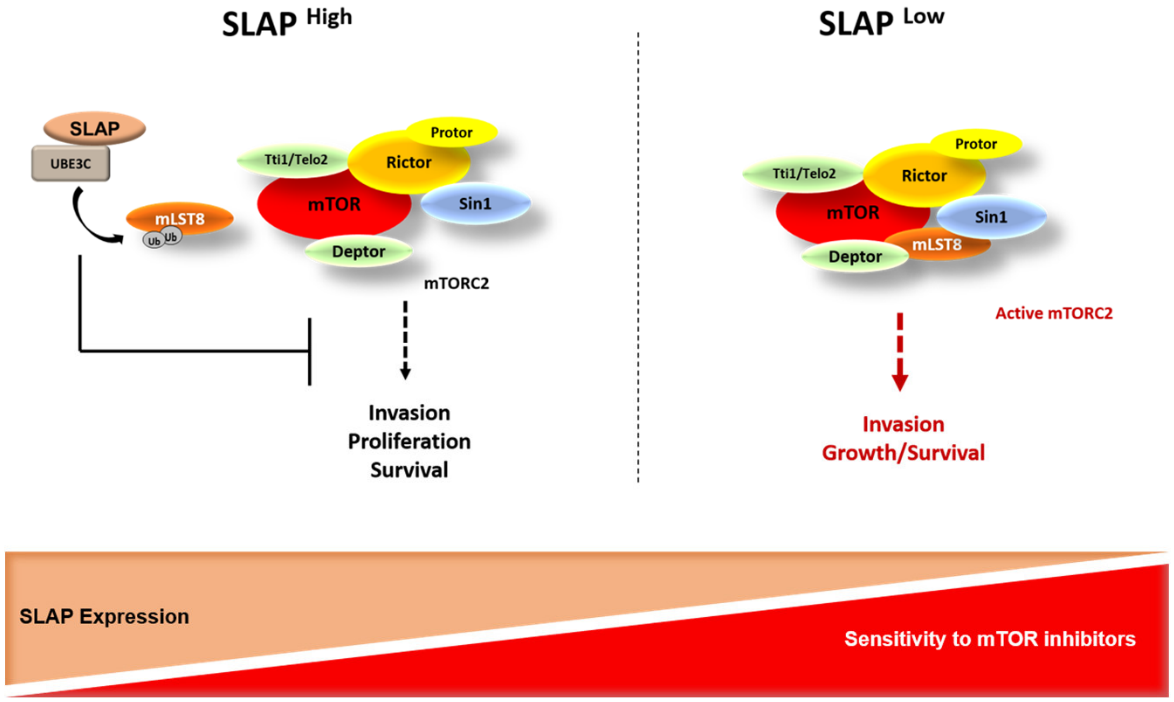


**Figure S13: Proposed model depicting the mechanism by which SLAP controls mTORC2 signaling and tumor cell response to mTORCi in CRC.**

Table S1

| Gene name | Log2 mean H/L normalized | Log2 mean iQBAQ | Log2 mean intensity |
|---|---|---|---|
| GEMIN4 | 4,309993449 | 18,8713518 | 26,14941967 |
| SLA | 4,296702096 | 26,90610553 | 31,96384623 |
| PRKDC | 3,387652029 | 19,63053969 | 28,95008684 |
| ARFGEF2 | 3,264941755 | 16,98804112 | 25,35872215 |
| IPO7 | 3,208933345 | 21,61483084 | 28,54371953 |
| GCN1 | 3,124488083 | 18,95903406 | 27,95638319 |
| ARHGEF28 | 2,952165812 | 14,70060227 | 22,6757512 |
| IPO9 | 2,915221307 | 18,98027813 | 26,09402032 |
| TIMM23 | 2,840251754 | 17,07169279 | 21,44187733 |
| LTN1 | 2,830992202 | 15,9189331 | 23,9174788 |
| ATAD3A | 2,812285942 | 19,47984735 | 26,24024131 |
| XPOT | 2,756461121 | 19,02554333 | 26,03028849 |
| IMMT | 2,745373613 | 18,69707616 | 26,25191484 |
| ERLIN2 | 2,685372076 | 18,83915457 | 24,67938023 |
| AKAP8L | 2,657098795 | 18,84570443 | 25,63110817 |
| TFRC | 2,622344729 | 16,87055737 | 23,44329382 |
| SLC25A13 | 2,608959053 | 20,76266481 | 27,37450654 |
| FGD5 | 2,552991764 | 13,06259148 | 20,48889533 |
| IPO4 | 2,546725756 | 18,87066974 | 26,11626542 |
| FASTKD5 | 2,543231917 | 17,12686846 | 24,4139888 |
| MTOR | 2,528196202 | 14,38924369 | 23,32368904 |
| SMN1 | 2,525926877 | 18,38498606 | 22,19233697 |
| IRAK1 | 2,522821613 | 13,93623304 | 20,41189574 |
| XPO1 | 2,492656498 | 19,88783839 | 27,41855372 |
| ERLIN1 | 2,47255277 | 16,70034489 | 22,3094882 |
| SQSTM1 | 2,47173372 | 18,02756048 | 23,82996075 |
| CIP2A | 2,463334749 | 13,46815732 | 20,62398684 |
| MON2 | 2,455114551 | 13,7824898 | 21,57320091 |
| UBE3C | 2,452147316 | 16,58837012 | 24,23452851 |
| SLC25A3 | 2,438807395 | 21,60357141 | 26,91567937 |
| SLC25A11 | 2,436916968 | 19,60069212 | 24,94975762 |
| PPP6R3 | 2,434855049 | 18,46945104 | 25,38762778 |
| AKAP8 | 2,424903322 | 14,2126357 | 20,64811763 |
| SLC25A12 | 2,365776481 | 17,55310177 | 24,28941377 |
| DDX20 | 2,362853256 | 17,05803049 | 24,45784818 |
| ATAD3B | 2,346999442 | 17,07302229 | 24,22596631 |
| TELO2 | 2,339593515 | 17,09296872 | 24,2800536 |
| SLC25A6 | 2,334415694 | 20,14161657 | 25,53842531 |
| SMPD4 | 2,333977055 | 15,77241768 | 23,18639941 |
| ZWILCH | 2,315493793 | 15,35426086 | 21,44310408 |
| DCAF1 | 2,304063161 | 15,6353289 | 23,22886761 |
| TTC27 | 2,30091437 | 17,65878097 | 24,2304248 |
| ANAPC5 | 2,286482042 | 12,54028951 | 21,42453265 |
| ARFGEF1 | 2,27604004 | 14,28035998 | 22,95092059 |
| TECR | 2,26856413 | 16,02512514 | 20,90493983 |
| SLC25A5 | 2,218171498 | 22,83249654 | 28,10784147 |
| IPO5 | 2,207476424 | 19,4311058 | 27,20708785 |

| | | | |
|---|---|---|---|
| **KNTC1** | 2,198075038 | 14,5091915 | 22,68147464 |
| **ABCD3** | 2,18516883 | 16,65483179 | 23,76955553 |
| **TNPO3** | 2,182734662 | 17,06741335 | 24,70771292 |
| **SLC25A4** | 2,177152408 | 16,07689908 | 21,39881674 |
| **PPP6C** | 2,176971665 | 16,26610328 | 22,02883101 |
| **USP34** | 2,169908969 | 13,21266863 | 22,59089873 |
| **UBR5** | **2,166233382** | **15,86971512** | **25,00392166** |
| **SMARCAD1** | 2,151642392 | 14,59204818 | 22,88850495 |
| **CSE1L** | 2,148988297 | 19,27480582 | 26,48687859 |
| **GBF1** | 2,147648588 | 12,933832 | 21,02394968 |
| **EXOC3** | 2,139109292 | 15,21285099 | 23,12169349 |
| **RICTOR** | 2,133289512 | 15,10680828 | 24,05003351 |
| **USP9X** | **2,131797994** | **13,91877133** | **23,02297504** |
| **SLC25A1** | 2,13116143 | 16,27877612 | 22,01528119 |
| **SLC25A10** | 2,124306037 | 16,93832378 | 21,98525919 |
| **SEC16A** | 2,121463087 | 14,9460229 | 23,96378993 |
| **ECPAS** | 2,120683728 | 15,25583877 | 23,79821284 |
| **DNAJB12** | 2,113967094 | 16,11583617 | 22,81425074 |
| **TMEM33** | 2,101560441 | 18,30201708 | 22,76143166 |
| **XPO7** | 2,092207417 | 16,46868714 | 23,68794787 |
| **EMD** | 2,086942281 | 18,57860365 | 23,89105127 |
| **ANAPC1** | 2,079827496 | 13,73258084 | 22,35381951 |
| **DHCR7** | 2,075566848 | 17,29732829 | 22,26911793 |
| **XPO4** | 2,073168996 | 14,5025625 | 22,0348312 |
| **NCAPD3** | 2,067076488 | 13,88645234 | 22,54081134 |
| **SLC12A2** | 2,066651801 | 13,53392587 | 20,41704349 |
| **GAK** | 2,057993168 | 14,20138371 | 22,08375199 |
| **HACD3** | 2,053644293 | 18,56982718 | 24,46288348 |
| **XPO5** | 2,050234667 | 16,98393994 | 24,3340179 |
| **HSD17B12** | 2,049096341 | 18,75330599 | 24,61940565 |
| **NDUFS2** | 2,047224281 | 15,61448843 | 22,19547434 |
| **SFXN4** | 2,032641581 | 15,10545841 | 20,89218407 |
| **TTI1** | 2,028834283 | 15,04440835 | 22,24276068 |
| **NUP205** | 2,024331488 | 16,20192607 | 24,35700549 |
| **PPP6R1** | 2,022190224 | 15,88396218 | 24,65178318 |
| **DNAJA3** | 2,013950496 | 18,01315181 | 24,75127251 |
| **SAV1** | 2,012586584 | 16,92431767 | 23,83402167 |
| **SQOR** | 1,999963926 | 16,62590381 | 22,93874613 |
| **PDS5A** | 1,994362501 | 14,7841236 | 22,69882786 |
| **VAC14** | 1,990640276 | 16,70725127 | 23,60589362 |
| **EXOC2** | 1,967353117 | 16,30400247 | 23,42076038 |
| **HSPB1** | 1,965544212 | 18,85088068 | 24,59237172 |
| **SLC16A1** | 1,962635434 | 17,66238095 | 24,26525229 |
| **NCAPG2** | 1,957766111 | 14,71881112 | 22,1027754 |
| **DNAAF5** | 1,936006589 | 16,21679637 | 23,25261635 |
| **TNPO1** | 1,930232737 | 17,29157543 | 24,5551721 |
| **ATP1A1** | 1,928616713 | 19,24855612 | 27,45459726 |
| **ZW10** | 1,922692642 | 17,22527649 | 23,77865348 |
| **RIC8A** | 1,905838657 | 16,70526834 | 23,49960878 |
| **ITGB4** | 1,901417201 | 16,13423913 | 24,02613965 |

| | | | |
|---|---|---|---|
| **NUP93** | 1,900039181 | 17,83239352 | 25,20833816 |
| **AGK** | 1,875347588 | 18,06666034 | 25,17429809 |
| **SEC61A1** | 1,866367355 | 16,80040744 | 21,80041334 |
| **EXOC5** | 1,863165355 | 15,88959665 | 22,91025234 |
| **BTAF1** | 1,862418322 | 13,92555135 | 22,17510564 |
| **TBC1D5** | 1,855571179 | 14,68540583 | 21,75286381 |
| **IPO11** | 1,854773669 | 15,95714437 | 23,37926677 |
| **RAF1** | 1,85459416 | 15,12587083 | 21,81268498 |
| **CAD** | 1,852851183 | 20,27682489 | 28,95136313 |
| **SLC25A22** | 1,851545821 | 17,67654481 | 23,89569607 |
| **NOM1** | 1,842455996 | 13,61488235 | 20,04116984 |
| **TUBB2B** | 1,839731205 | 19,3201953 | 24,64212707 |
| **ATP2A2** | 1,837082081 | 18,64592107 | 25,82068513 |
| **ABCC1** | 1,829504564 | 13,069198 | 21,09236401 |
| **TBL2** | 1,82836755 | 16,7630351 | 23,40232667 |
| **UQCRC2** | 1,824183997 | 18,39668282 | 24,43682014 |
| **TIMM50** | 1,808858642 | 17,83992548 | 23,70290485 |
| **GCDH** | 1,808817483 | 15,62470028 | 21,90623248 |
| **UBAC2** | 1,79363238 | 16,76650734 | 21,85395673 |
| **MYBBP1A** | 1,790202046 | 16,50830546 | 24,35474556 |
| **CDC16** | 1,788059237 | 14,98116588 | 21,22235665 |
| **EXOC7** | 1,787892067 | 16,51676188 | 23,45355315 |
| **SFXN3** | 1,787139878 | 17,07660686 | 23,06666012 |
| **SMC4** | 1,780925977 | 16,92920831 | 24,43796528 |
| **HSPA1A** | 1,776047811 | 18,79217388 | 26,3377594 |
| **DHRS7B** | 1,771505287 | 15,0266243 | 20,0266333 |
| **PNPLA6** | 1,762207067 | 13,39544488 | 21,94215747 |
| **PPP6R2** | 1,754673718 | 14,39459667 | 21,92421617 |
| **DDB1** | 1,754545484 | 16,33854315 | 24,41734315 |
| **VDAC3** | 1,746470447 | 17,90542422 | 23,26668495 |
| **TOMM22** | 1,742459014 | 16,81890667 | 22,48874172 |
| **MSH2** | 1,739660967 | 18,27590169 | 25,47383694 |
| **CAMK2D** | 1,736994453 | 16,24948386 | 21,94994162 |
| **KPNB1** | 1,725275859 | 18,94007458 | 26,19759297 |
| **FANCD2** | 1,709356842 | 14,11856262 | 21,58914913 |
| **CHTF18** | 1,702325056 | 14,04277086 | 21,254733 |
| **FANCI** | 1,701475123 | 15,64800173 | 23,49936548 |
| **ACSL3** | 1,698337035 | 16,31769289 | 22,74988743 |
| **SFXN1** | 1,696884066 | 18,64745023 | 24,07477149 |
| **DNAJA2** | 1,69223379 | 19,33091834 | 26,10386272 |
| **RPN2** | 1,689589886 | 15,83947446 | 21,47395871 |
| **MYO1B** | 1,682633195 | 18,97119363 | 26,27032575 |
| **HSPA6** | 1,676959415 | 16,8141944 | 23,0236463 |
| **ANKRD28** | 1,66830068 | 17,41077871 | 24,84570252 |
| **DNAJA1** | 1,666968621 | 18,47901182 | 26,20882807 |
| **MMS19** | 1,666165804 | 16,36670583 | 24,04092077 |
| **HSPA8** | 1,660822148 | 23,17626792 | 29,78958066 |
| **SMC2** | 1,6599398 | 18,22453874 | 25,77533472 |
| **PIK3R4** | 1,653289337 | 14,01445642 | 21,33667508 |
| **HLA-C** | 1,648925577 | 16,29863541 | 22,45202434 |

| | | | |
|---|---|---|---|
| **RCN2** | 1,645909082 | 17,20056979 | 22,00793383 |
| **VPS53** | 1,636705815 | 13,25301523 | 21,41736804 |
| **EXOC4** | 1,63217515 | 16,13056989 | 23,48279335 |
| **CIAO2B** | 1,622789819 | 17,6042202 | 21,26564099 |
| **TRIM25** | **1,610014907** | **16,40801455** | **23,29697183** |
| **EPHA2** | 1,592046321 | 17,50659427 | 24,62515403 |
| **MAPKAP1** | 1,588516746 | 16,35057091 | 22,4087796 |
| **PHB2** | 1,587765005 | 18,88852724 | 25,03002182 |
| **EXOC1** | 1,587268969 | 14,04410947 | 22,37124489 |
| **ATP5F1C** | 1,580402807 | 19,12557995 | 24,15905552 |
| **NDUFA9** | 1,577956467 | 14,22421275 | 20,73803837 |
| **NDUFS7** | 1,570900074 | 14,84785666 | 21,62473954 |
| **CPT1A** | 1,560356315 | 18,13522497 | 24,6348225 |
| **ESYT2** | 1,536484006 | 17,9359247 | 25,29025786 |
| **UFL1** | 1,527921061 | 15,61436477 | 22,72358401 |
| **TIMMDC1** | 1,526519596 | 13,08594266 | 19,77288693 |
| **AIFM1** | 1,524080423 | 17,99254378 | 24,66867843 |
| **WAPL** | 1,496700998 | 14,22349898 | 22,01692524 |
| **UBR4** | 1,493954676 | 9,840568125 | 21,03447301 |
| **SKP1** | 1,479334061 | 17,08792549 | 21,77561939 |
| **HAX1** | 1,479230575 | 15,04526196 | 21,97365253 |
| **OSBPL8** | 1,475577842 | 15,80402175 | 23,48049423 |
| **NFXL1** | 1,473024906 | 14,63618088 | 21,68257517 |
| **COPG2** | 1,468426915 | 14,67343791 | 21,53161626 |
| **TRIP13** | 1,459833773 | 17,84840084 | 23,96001396 |
| **AFG3L2** | 1,459693947 | 15,20789133 | 22,09845537 |
| **DDOST** | 1,452314272 | 17,66340985 | 24,09168202 |
| **HNRNPM** | 1,450749378 | 20,91883849 | 27,97692781 |
| **RNF213** | 1,446838531 | 12,11814081 | 22,15100456 |
| **SLC3A2** | 1,446291501 | 17,97719233 | 24,43592836 |
| **SLC7A5** | 1,44618566 | 16,10702768 | 21,54022013 |
| **PHB1** | 1,441881794 | 17,35457654 | 22,82917043 |
| **PDXDC1** | 1,424680237 | 15,03411115 | 21,71123453 |
| **INF2** | 1,418243904 | 15,13451991 | 22,32218123 |
| **RPN1** | 1,417686072 | 18,04824916 | 25,29866002 |
| **ABCF2** | 1,414622725 | 17,10572781 | 23,44631365 |
| **IPO8** | 1,409562979 | 17,86770305 | 25,12851652 |
| **CLPX** | 1,40165772 | 14,43193282 | 21,31984997 |
| **AAR2** | 1,399553963 | 16,70787585 | 21,79533526 |
| **MSH6** | 1,399061721 | 18,32831262 | 25,74005659 |
| **UNC45A** | 1,391786068 | 16,59220979 | 23,53673494 |
| **SKIC3** | 1,380526285 | 15,25252377 | 23,25099382 |
| **PELO** | 1,379787316 | 15,84730208 | 23,21973515 |
| **TIMM44** | 1,378789031 | 14,3157864 | 21,20192209 |
| **DICER1** | 1,376123667 | 12,4329681 | 20,46741583 |
| **RFC5** | 1,374306889 | 17,18577558 | 24,39082093 |
| **RFC2** | 1,372562082 | 15,59143469 | 21,38847983 |
| **TFB2M** | 1,366783829 | 15,59054807 | 21,32337294 |
| **BRAT1** | 1,36518856 | 15,36162399 | 22,36337871 |
| **MYCBP2** | 1,363227161 | 11,94834287 | 22,68633495 |

| | | | |
|---|---|---|---|
| **HEATR1** | 1,36056096 | 13,76311206 | 22,15871405 |
| **EXOC6** | 1,34718373 | 15,50404307 | 21,96343424 |
| **WDR11** | 1,334368067 | 15,3274006 | 23,00958627 |
| **FRYL** | 1,324330594 | 17,01574861 | 25,98852775 |
| **POLRMT** | 1,321264296 | 13,101051 | 22,33396332 |
| **SKIC2** | 1,320715727 | 13,87811479 | 21,8785489 |
| **GNAI2** | 1,305707895 | 16,01486319 | 21,14290271 |
| **VPS16** | 1,302718727 | 15,12685502 | 22,38750726 |
| **MT-CO2** | 1,298247688 | 16,48152617 | 21,17255447 |
| **MACC1** | 1,293046676 | 14,87605164 | 21,55822575 |
| **ARHGEF2** | 1,292762145 | 15,73252608 | 23,72353888 |
| **COG2** | 1,286504686 | 16,0974355 | 22,73600079 |
| **CIAO1** | 1,278074005 | 14,96382931 | 21,69781035 |
| **COPB1** | 1,276893641 | 18,03709289 | 25,55572797 |
| **OPA1** | 1,262342963 | 14,84138416 | 22,35008238 |
| **PPM1G** | 1,255440324 | 16,79712954 | 23,50767545 |
| **ZC3HAV1** | 1,247775581 | 14,92927382 | 22,32176849 |
| **AP1G1** | 1,245536217 | 14,88821685 | 21,01750304 |
| **PTPN12** | 1,224904656 | 15,41981757 | 23,01622249 |
| **FAM91A1** | 1,224657742 | 15,83524447 | 22,26941284 |
| **HNRNPF** | 1,214186983 | 21,30927539 | 26,99691469 |
| **HADHA** | 1,21138606 | 17,90550453 | 24,59919582 |
| **SEC13** | 1,209297401 | 15,6915527 | 23,01708954 |
| **RPAP1** | 1,206466131 | 13,10860068 | 20,34333466 |
| **HSD17B11** | 1,192765469 | 15,34627132 | 22,4008381 |
| **DDX5** | 1,171100018 | 19,91605827 | 26,19347543 |
| **NDUFS1** | 1,167186657 | 15,20253965 | 22,83100203 |
| **TUBA1B** | 1,146850694 | 23,40935381 | 29,29492949 |
| **COPA** | 1,132905811 | 18,02072318 | 25,92391404 |
| **TUBB** | 1,131106544 | 23,49120599 | 29,44714358 |
| **RPTOR** | 1,11742909 | 13,33590206 | 21,55009855 |
| **NAMPT** | 1,108736311 | 16,84684929 | 23,7664146 |
| **AKAP11** | 1,107353104 | 11,30514978 | 18,88436929 |
| **CNP** | 1,105678125 | 15,30524239 | 22,12792381 |
| **RPL9** | 1,103329834 | 19,04116538 | 23,62794082 |
| **STAT1** | 1,092613399 | 15,59359537 | 22,20960289 |
| **COPG1** | 1,09254575 | 16,9427932 | 24,08936686 |
| **ARHGAP4** | 1,092072092 | 14,22087707 | 20,91363061 |
| **GEMIN5** | 1,079292791 | 13,28526253 | 21,06688453 |
| **MYOF** | 1,075601108 | 15,27035617 | 24,22518429 |
| **CAND1** | 1,070480902 | 17,69550019 | 25,31739071 |
| **HSP90AB2P** | 1,070217597 | 16,67699925 | 22,84193028 |
| **YTHDF2** | 1,06380208 | 17,01129278 | 22,86096823 |
| **PKP2** | 1,059839779 | 17,29368867 | 24,12228087 |
| **RFC4** | 1,053111328 | 17,58715243 | 23,26708041 |
| **IDH3B** | 1,035975741 | 16,64079097 | 23,62281983 |
| **CEP170** | 1,028852026 | 10,7995784 | 20,10646922 |
| **SLC1A5** | 1,020555534 | 20,40571211 | 25,54454024 |
| **CTNNB1** | 1,016686513 | 15,85140013 | 23,21943975 |
| **TCP1** | 1,009586011 | 18,08423746 | 24,47095488 |

| | | | |
|---|---|---|---|
| **ERAL1** | 1,00666904 | 15,44274915 | 21,84396295 |
| **MCM7** | 0,998325537 | 17,67587665 | 25,01971017 |
| **TARS2** | 0,994326329 | 15,29822967 | 21,92637756 |
| **ATXN10** | 0,981122002 | 15,99031009 | 21,69075282 |
| **LRPPRC** | 0,978969292 | 19,74701284 | 27,84461753 |
| **TUBA1C** | 0,974712817 | 18,00844686 | 23,85760912 |
| **AKAP13** | 0,973872087 | 12,49290469 | 20,54345858 |
| **PKP3** | 0,960427675 | 17,67380594 | 24,61230549 |
| **VDAC2** | 0,958360176 | 18,40228763 | 24,1121912 |
| **HSP90AB1** | 0,954568603 | 22,16357127 | 28,67872096 |
| **TUBB4B** | 0,953451467 | 21,20970125 | 26,69221091 |
| **SLC2A1** | 0,950493306 | 17,79061577 | 23,62920526 |
| **PFKP** | 0,941131329 | 15,58875562 | 22,23189733 |
| **CCT3** | 0,938122671 | 20,44988179 | 27,35592804 |
| **MTREX** | 0,93104004 | 14,47256815 | 21,78348748 |
| **PDCD6IP** | 0,929828848 | 14,98323115 | 22,2158063 |
| **CTNND1** | 0,925885525 | 16,08309145 | 24,04354578 |
| **UPF1** | 0,924378625 | 16,93206093 | 24,82026312 |
| **SAMHD1** | 0,919225661 | 17,63606521 | 23,98779947 |
| **ELP1** | 0,916807821 | 16,03744686 | 24,03066455 |
| **ATP5F1B** | 0,916807791 | 16,08316696 | 21,94110685 |
| **HADHB** | 0,914105199 | 16,59667022 | 23,23577293 |
| **YTHDC2** | 0,912841471 | 11,83749653 | 19,54032736 |
| **RPS27A** | 0,904837298 | 22,20773169 | 25,92947815 |
| **COPB2** | 0,900232475 | 17,71699804 | 25,0085124 |
| **DNM2** | 0,899149817 | 14,74826624 | 22,42011785 |
| **ASPH** | 0,896465635 | 16,74603313 | 23,81570082 |
| **SPIN2B** | 0,894061049 | 14,30784117 | 19,21473967 |
| **ACADM** | 0,883672914 | 18,41260981 | 24,0280469 |
| **MYO1C** | 0,875072338 | 18,59901083 | 25,73384797 |
| **PALD1** | 0,869002735 | 13,83444141 | 21,75458417 |
| **TUBGCP3** | 0,863462752 | 14,65991787 | 22,15996657 |
| **PDCD6** | 0,856149131 | 18,37156397 | 23,20854159 |
| **RPL30** | 0,85451436 | 19,97790596 | 24,01137617 |
| **SKIC8** | 0,853835981 | 14,65653659 | 21,20450746 |
| **POLR2A** | 0,853077452 | 12,67865148 | 20,80331761 |
| **RPL36** | 0,851958872 | 19,37608016 | 22,04934117 |
| **RPL31** | 0,841852382 | 19,75594812 | 23,56330925 |
| **DDX17** | 0,841678007 | 17,35439288 | 24,30187237 |
| **SERPINH1** | 0,833942526 | 17,65255558 | 24,25666711 |
| **PLAA** | 0,831309757 | 14,98250345 | 21,5060738 |
| **CNOT1** | 0,829876614 | 16,31751861 | 24,67857994 |
| **MRPL13** | 0,829635855 | 19,66985103 | 25,23773358 |
| **ACACA** | 0,828712844 | 12,47652682 | 22,28543759 |
| **TNS4** | 0,827466745 | 13,55286519 | 19,99984039 |
| **MYO1D** | 0,824564235 | 14,44803455 | 23,30991372 |
| **MARS1** | 0,820349186 | 19,5701769 | 26,62197635 |
| **MRPS18B** | 0,818646096 | 13,35842227 | 20,56470524 |
| **TRIM28** | 0,815329494 | 17,92833331 | 25,25376113 |
| **RPL7A** | 0,815138232 | 20,9848653 | 25,8336904 |

| | | | |
|---|---|---|---|
| **CTPS1** | 0,811471059 | 19,02640817 | 25,66356636 |
| **HK2** | 0,806860166 | 18,24545274 | 24,84146485 |
| **EIF4G2** | 0,800173032 | 17,12334816 | 24,45132013 |
| **RPLP2** | 0,799971436 | 19,8640128 | 24,04499126 |
| **PPP1CA** | 0,796140834 | 15,11566686 | 21,56526326 |
| **DHX30** | 0,793812675 | 15,78881178 | 23,61334015 |
| **DDX39B** | 0,787237411 | 15,64995407 | 21,86425971 |
| **NSF** | 0,786512731 | 13,50847157 | 20,4224604 |
| **PANK4** | 0,783091983 | 14,48658785 | 21,22874042 |
| **ALDH18A1** | 0,770321612 | 12,73724734 | 19,48818927 |
| **CYB5B** | 0,769898664 | 15,70053003 | 21,48901951 |
| **RPLP0** | 0,76496892 | 22,35444741 | 27,83930049 |
| **RPL7** | 0,764714163 | 20,89922906 | 25,82055529 |
| **HSPD1** | 0,763836471 | 16,79004298 | 23,14630473 |
| **MDN1** | 0,760902187 | 11,76869439 | 23,08063821 |
| **ARF4** | 0,759780827 | 18,24305618 | 23,27289495 |
| **PTPN23** | 0,753814135 | 11,4441521 | 20,09188643 |
| **RPL12** | 0,753290739 | 20,73710317 | 25,36897388 |
| **RPL27** | 0,751186585 | 19,64835758 | 24,35169161 |
| **AP2A2** | 0,75039209 | 13,38607803 | 20,61941461 |
| **MATR3** | 0,741488486 | 15,44424922 | 22,36629909 |
| **CCT4** | 0,735435513 | 16,47561206 | 22,66952928 |
| **RPL5** | 0,733886452 | 20,40089541 | 25,87861126 |
| **TUBG1** | 0,732963843 | 15,44887988 | 23,18504358 |
| **IGF2BP3** | 0,731692992 | 16,99773815 | 23,26405692 |
| **HSPA12A** | 0,728620019 | 15,66930586 | 21,98416995 |
| **STRAP** | 0,726598711 | 24,225199 | 31,19619413 |
| **PYCR1** | 0,722670041 | 15,81899818 | 21,22159971 |
| **CALM1** | 0,719971977 | 16,68680598 | 21,71162665 |
| **CCT6A** | 0,718672155 | 16,93859563 | 23,67614232 |
| **HSPA2** | 0,717473526 | 14,33231633 | 22,55233482 |
| **AARS1** | 0,71677143 | 15,2182605 | 22,21624062 |
| **KIF2A** | 0,711803277 | 15,07014751 | 21,69214438 |
| **EXOSC3** | 0,710525429 | 14,5428908 | 20,23296501 |
| **RPL4** | 0,699587173 | 19,77822903 | 25,91694756 |
| **CCT8** | 0,699344329 | 16,44848739 | 22,92116781 |
| **RPL3** | 0,687986333 | 19,04934516 | 27,03703364 |
| **YWHAB** | 0,686971075 | 15,36651574 | 20,17385603 |
| **RPL18A** | 0,686741074 | 19,91561732 | 24,09725009 |
| **RARS1** | 0,683411982 | 18,61739835 | 25,34863343 |
| **PFKM** | 0,681973912 | 16,17454213 | 23,12983221 |
| **RPL18** | 0,678651875 | 19,833166 | 23,64048047 |
| **SYNRG** | 0,67784645 | 12,73307892 | 21,1580771 |
| **EEF1E1** | 0,673013605 | 17,45875904 | 22,15939178 |
| **SYNCRIP** | 0,672440413 | 18,88471111 | 25,74668284 |
| **RPL23** | 0,666271832 | 20,44285105 | 24,67895597 |
| **RPL24** | 0,665665616 | 21,02439842 | 25,02912319 |
| **DNAJB11** | 0,658417166 | 16,54228627 | 22,77315933 |
| **PCNA** | 0,651606638 | 21,02059207 | 26,58501559 |
| **EXOSC7** | 0,648373382 | 15,99028128 | 21,7431218 |

| | | | |
|---|---|---|---|
| **SNRNP200** | 0,645455741 | 17,75883479 | 26,23325985 |
| **TUFM** | 0,623351872 | 20,93695898 | 27,49566724 |
| **NCAPD2** | 0,621764138 | 14,35520883 | 22,13871177 |
| **ILF2** | 0,620085853 | 18,02629109 | 24,06633713 |
| **LRRC59** | 0,618833774 | 18,79794644 | 24,56778715 |
| **IARS1** | 0,618332661 | 19,24194747 | 26,96426086 |
| **PCBP2** | 0,617549259 | 19,89339437 | 25,81474947 |
| **KRT18** | 0,615981179 | 20,40944964 | 26,95275585 |
| **MRPS22** | 0,614442846 | 16,15279977 | 22,86529856 |
| **FEN1** | 0,604356011 | 16,41186403 | 22,95112816 |
| **POLR2B** | 0,599730404 | 15,85908986 | 23,2101283 |
| **CCT5** | 0,596343655 | 16,8005292 | 23,74299165 |
| **RPL11** | 0,594519895 | 20,72207095 | 25,18364103 |
| **SMC1A** | 0,593258151 | 11,74830472 | 19,69179803 |
| **PRPF8** | 0,592668301 | 16,78979767 | 25,87213471 |
| **MAGED2** | 0,583840011 | 16,07096646 | 22,75589516 |
| **ACOT8** | 0,583839935 | 16,11523335 | 21,55577867 |
| **MRPS27** | 0,583759783 | 16,54089993 | 22,65196379 |
| **MED23** | 0,575360702 | 11,50060956 | 18,62988782 |
| **GMDS** | 0,57347149 | 16,60157736 | 23,70217565 |
| **IGF2BP2** | 0,568080786 | 15,64523623 | 21,49623939 |
| **GARS1** | 0,567415655 | 17,2077579 | 24,31695359 |
| **RPL6** | 0,564410685 | 19,26232582 | 24,19189538 |
| **EEF1G** | 0,560062734 | 19,46837081 | 25,80927193 |
| **AIMP2** | 0,556862636 | 17,74851391 | 24,58699673 |
| **MRTO4** | 0,555178127 | 15,65879724 | 21,73798887 |
| **GTPBP4** | 0,552716407 | 14,54436077 | 21,76940419 |
| **RPL27A** | 0,541498323 | 20,46385893 | 24,30428855 |
| **MTHFD1** | 0,537985153 | 18,65861247 | 25,81847634 |
| **HSP90AA1** | 0,535256745 | 19,93852958 | 26,42660277 |
| **AIMP1** | 0,530836487 | 17,32203376 | 24,99167361 |
| **HNRNPH1** | 0,529071318 | 16,93222244 | 22,74002227 |
| **PTGES3** | 0,526870098 | 17,6526116 | 22,41256606 |
| **ACP1** | 0,515712469 | 16,36179531 | 20,53171143 |
| **RNF40** | 0,515106848 | 14,31887228 | 21,74760843 |
| **SRPK1** | 0,512985321 | 15,93905115 | 22,67210762 |
| **TNS3** | 0,510708763 | 13,39373007 | 22,10833813 |
| **DCTN1** | 0,506297289 | 13,75540226 | 21,46413489 |
| **KARS1** | 0,505857095 | 20,46722343 | 27,43494381 |
| **KEAP1** | 0,504789878 | 16,40224044 | 22,84752282 |
| **EFTUD2** | 0,501753378 | 18,16102032 | 25,67773799 |
| **GIGYF2** | 0,500241169 | 12,94778305 | 20,58190424 |
| **DNAJC10** | 0,499510006 | 15,91259311 | 21,95701085 |
| **STOML2** | 0,499101727 | 17,77678427 | 23,20407954 |
| **LUC7L2** | 0,498781921 | 19,15325513 | 24,6076855 |
| **HNRNPUL1** | 0,497160993 | 15,74325485 | 22,64057976 |
| **FXR2** | 0,495950945 | 14,05518074 | 21,19410195 |
| **CNN2** | 0,494507797 | 18,66792464 | 23,7381023 |
| **CDK6** | 0,491117649 | 16,72314096 | 22,58475091 |
| **C1QBP** | 0,490125122 | 19,90313358 | 25,47525004 |

| | | | |
|---|---|---|---|
| **ILF3** | 0,48583897 | 18,11573628 | 25,032322 |
| **OXA1L** | 0,481980826 | 16,05586141 | 22,38360165 |
| **SSB** | 0,480730428 | 15,93945678 | 21,57425474 |
| **RPL17** | 0,479196071 | 20,53574746 | 25,99351034 |
| **IGF2BP1** | 0,477369932 | 19,54004351 | 25,76057242 |
| **EPRS1** | 0,47643358 | 19,43457801 | 27,59168334 |
| **RPL21** | 0,474728465 | 19,93745237 | 24,59341316 |
| **RRM1** | 0,470458797 | 16,43807708 | 23,75826958 |
| **H2BC4** | 0,469733161 | 19,64716913 | 23,49059124 |
| **MTHFD2** | 0,469729683 | 15,97143877 | 21,55627777 |
| **YWHAG** | 0,469417129 | 17,39403278 | 22,82400493 |
| **DYNC1H1** | 0,468114078 | 13,6673703 | 24,20587993 |
| **PIGT** | 0,464720541 | 15,41330838 | 21,53331262 |
| **LARS1** | 0,462191956 | 18,57028439 | 25,91974576 |
| **CDK1** | 0,461339968 | 20,64994846 | 27,03274938 |
| **PABPC4** | 0,459064342 | 15,73500944 | 22,03603943 |
| **RUVBL2** | 0,45280626 | 16,35905174 | 23,36595216 |
| **CCT2** | 0,45259191 | 15,62686877 | 22,40983287 |
| **PPP2CA** | 0,445249935 | 16,4532492 | 22,32993718 |
| **HNRNPD** | 0,444608781 | 20,26295659 | 25,10176071 |
| **RPL13** | 0,443728662 | 20,70483633 | 24,87477415 |
| **ESYT1** | 0,443642032 | 19,20224182 | 26,65386153 |
| **RNGTT** | 0,441270998 | 16,59991284 | 23,40493122 |
| **TFIP11** | 0,439729512 | 15,19453885 | 22,36235339 |
| **GANAB** | 0,437263232 | 14,50629311 | 22,46335341 |
| **GNAS** | 0,436375102 | 14,16136811 | 20,24882632 |
| **RBM39** | 0,431105905 | 17,63906038 | 23,46213461 |
| **DDX21** | 0,429749811 | 17,39647657 | 23,82473336 |
| **PPFIBP1** | 0,42889275 | 12,36283879 | 21,1219938 |
| **SEPTIN9** | 0,427320134 | 13,2952093 | 19,68747593 |
| **DIMT1** | 0,418398679 | 15,5984015 | 21,22132436 |
| **MCM4** | 0,417142342 | 13,79060644 | 21,47862349 |
| **OBI1** | 0,413904527 | 19,3598432 | 26,43000561 |
| **RPL38** | 0,412420281 | 20,22342748 | 24,06030188 |
| **TCERG1** | 0,411588955 | 13,81521649 | 20,38398126 |
| **VAPA** | 0,401408408 | 15,5137276 | 21,12051802 |
| **HSPA5** | 0,395062775 | 20,37288206 | 26,81667107 |
| **DHX9** | 0,390553976 | 17,57175264 | 25,49054023 |
| **CAPRIN1** | 0,390062263 | 16,92179833 | 23,16063859 |
| **PSMD6** | 0,386535095 | 17,05498227 | 23,41965935 |
| **RNH1** | 0,385154891 | 17,01518998 | 23,0222075 |
| **FRG1** | 0,384392487 | 15,09384393 | 20,09276832 |
| **EEF1A1** | 0,381984699 | 23,28434592 | 30,15612904 |
| **QPCTL** | 0,381479032 | 17,93885977 | 24,5956721 |
| **YWHAQ** | 0,375734554 | 18,35720404 | 23,5646316 |
| **FARSB** | 0,373453302 | 17,71501014 | 23,99446739 |
| **YWHAE** | 0,372692127 | 19,7461272 | 24,99446739 |
| **CDK5** | 0,367930125 | 16,27739638 | 22,27615092 |
| **YWHAZ** | 0,367613378 | 17,96027727 | 23,12027016 |
| **SMCHD1** | 0,366625305 | 13,46797483 | 22,08643257 |

| | | | |
|---|---|---|---|
| **NUP155** | 0,364721792 | 16,29630086 | 23,30979347 |
| **EIF2AK3** | 0,362785962 | 14,43278111 | 21,93793841 |
| **HSPA4** | 0,361038382 | 16,47028816 | 23,74163115 |
| **TRIO** | 0,354909073 | 11,11115743 | 19,68119723 |
| **PRPF6** | 0,349327534 | 16,8774519 | 24,11362518 |
| **CBX3** | 0,348402435 | 16,2960917 | 20,90949565 |
| **CLTC** | 0,347552301 | 16,4338071 | 24,96557881 |
| **HSPBP1** | 0,342674173 | 15,85858814 | 22,14662585 |
| **POLD1** | 0,336656636 | 13,64876578 | 21,10640106 |
| **SUCLG1** | 0,335307931 | 16,84564522 | 21,73014448 |
| **RECQL** | 0,334911493 | 16,16335747 | 23,23648841 |
| **EIF4A1** | 0,328798217 | 20,94477639 | 27,19522901 |
| **PDLIM5** | 0,327687359 | 14,39178061 | 20,43616862 |
| **EXOSC10** | 0,324637818 | 15,00266854 | 23,03847519 |
| **DDX23** | 0,322735774 | 15,01809873 | 21,82078249 |
| **MCM3** | 0,317960048 | 18,47168712 | 25,81588971 |
| **SNRPD3** | 0,312084285 | 22,06733873 | 26,84455767 |
| **ANKRD17** | 0,310398279 | 12,14436081 | 20,53469044 |
| **NSUN2** | 0,303926826 | 17,34901624 | 25,04234939 |
| **NASP** | 0,303517743 | 15,07313738 | 21,7607416 |
| **EDC4** | 0,301002295 | 15,6808265 | 24,34629044 |
| **CSDE1** | 0,296369294 | 15,3330039 | 21,95663281 |
| **MAP4** | 0,295429143 | 16,44140964 | 24,26658964 |
| **CCDC88A** | 0,284557546 | 15,66478059 | 24,05133973 |
| **RPL23A** | 0,273957451 | 21,37983347 | 25,07238812 |
| **LANCL2** | 0,270648502 | 17,61994292 | 24,01736144 |
| **MAPK6** | 0,266216857 | 16,28794079 | 22,89933356 |
| **NPM1** | 0,260495438 | 19,04376514 | 25,13053411 |
| **GFPT1** | 0,260025644 | 16,13201768 | 23,05678984 |
| **HSPH1** | 0,258659621 | 17,23467088 | 24,66200927 |
| **NCL** | 0,258153086 | 21,52347773 | 28,13646571 |
| **CHD4** | 0,25631024 | 13,12589773 | 21,51058777 |
| **FXR1** | 0,255847159 | 14,95648053 | 23,21398366 |
| **NCBP1** | 0,249384611 | 14,99423556 | 21,90351365 |
| **DNAJC13** | 0,248146186 | 16,02079913 | 24,51169395 |
| **PYGL** | 0,246306756 | 13,77347579 | 20,80162417 |
| **SNRPB** | 0,246063442 | 18,88285795 | 24,09297259 |
| **ERP44** | 0,243851892 | 16,27809852 | 21,60002056 |
| **ATXN2L** | 0,241933704 | 15,1053793 | 22,39916977 |
| **MRPL14** | 0,241046888 | 15,23954881 | 19,91043779 |
| **PABPC1** | 0,240627754 | 19,52269263 | 26,07002811 |
| **SFPQ** | 0,235849609 | 16,94835963 | 24,07139275 |
| **GART** | 0,235604524 | 17,40999028 | 25,27457361 |
| **FAM120A** | 0,235257323 | 14,77579887 | 22,45000299 |
| **PSMD3** | 0,235171557 | 16,98773403 | 23,26710423 |
| **VARS1** | 0,234865099 | 14,76707403 | 22,44652401 |
| **PSMD13** | 0,233356365 | 17,36248039 | 23,5949853 |
| **HTATSF1** | 0,230756362 | 11,82898269 | 21,31115257 |
| **EIF2S3** | 0,230510448 | 16,53267553 | 23,35460882 |
| **ADAP1** | 0,222371808 | 15,72166886 | 22,44239549 |

| | | | |
|---|---|---|---|
| **CAPZA2** | 0,221444043 | 16,49174873 | 22,59773433 |
| **VIL1** | 0,221155287 | 18,80956928 | 25,60140795 |
| **DHX36** | 0,221010961 | 14,74408049 | 23,2256158 |
| **DLAT** | 0,220082246 | 14,54226611 | 21,08390344 |
| **JAK1** | 0,216816991 | 19,0428136 | 26,6170369 |
| **CYFIP1** | 0,215782159 | 15,56354487 | 26,50075076 |
| **SUGT1** | 0,214311338 | 15,18338985 | 21,48202602 |
| **ASCC3** | 0,213751617 | 14,58642934 | 23,30803956 |
| **G3BP1** | 0,213502824 | 16,65774178 | 23,18597113 |
| **NCAPG** | 0,212569336 | 15,01680829 | 21,57138726 |
| **CKAP5** | 0,207826255 | 15,21516262 | 23,53824767 |
| **CCAR2** | 0,207193176 | 17,09195528 | 24,63158281 |
| **YBX1** | 0,206601549 | 20,09324346 | 26,21437283 |
| **HSP90B1** | 0,206393182 | 17,1310531 | 24,36963959 |
| **ARPC2** | 0,20497537 | 16,1791186 | 21,77128634 |
| **FUS** | 0,204725009 | 18,31136251 | 24,18055873 |
| **HNRNPAB** | 0,204591442 | 17,89719551 | 22,96049717 |
| **MAT2A** | 0,201809492 | 16,1398311 | 22,28571978 |
| **EEF1B2** | 0,198163063 | 18,53339818 | 23,46146862 |
| **LARP4** | 0,191373136 | 14,75994018 | 22,59900659 |
| **WWP2** | 0,190994057 | 13,34177361 | 19,86533255 |
| **PRAG1** | 0,190572708 | 13,51886212 | 21,64376813 |
| **ARPC1B** | 0,189645372 | 15,36038988 | 21,99863358 |
| **QARS1** | 0,187869084 | 17,90243989 | 24,93850984 |
| **MTHFD1L** | 0,184005171 | 18,10486489 | 25,1315026 |
| **ANKFY1** | 0,180843816 | 17,00171605 | 25,25000593 |
| **SUGP1** | 0,17636112 | 14,50783593 | 21,10684831 |
| **DNMT1** | 0,174150726 | 12,82919418 | 20,66211433 |
| **HSPA9** | 0,173067712 | 19,09404503 | 25,41496518 |
| **ZC3HAV1L** | 0,172877939 | 18,09760696 | 22,90498392 |
| **NQO1** | 0,169411969 | 14,48104257 | 22,12559257 |
| **HNRNPU** | 0,164309977 | 20,77458493 | 27,91159387 |
| **AP2B1** | 0,163820709 | 14,80057976 | 22,15367242 |
| **PCBP1** | 0,163640387 | 18,61466312 | 24,70102297 |
| **DSTN** | 0,162725684 | 18,33504718 | 22,91012422 |
| **PSMC6** | 0,160576016 | 17,03413981 | 23,75110051 |
| **PSMD2** | 0,156991947 | 17,18668816 | 25,15143406 |
| **FLII** | 0,156461332 | 14,92585111 | 22,17072217 |
| **VDAC1** | 0,155069199 | 16,46525816 | 22,1972955 |
| **DSP** | 0,152313239 | 12,13035605 | 20,99895488 |
| **TNIK** | 0,151014525 | 13,49760239 | 20,98361342 |
| **TRAP1** | 0,144655494 | 17,50289395 | 24,40227896 |
| **HNRNPA1** | 0,143297596 | 18,85157216 | 24,58924668 |
| **XRN2** | 0,141777216 | 16,80081018 | 23,81487331 |
| **CAPZB** | 0,140929171 | 17,23671807 | 23,12763524 |
| **CSNK2A1** | 0,13547795 | 17,57690255 | 24,49301691 |
| **HDLBP** | 0,135206462 | 14,57740916 | 22,78864718 |
| **FASN** | 0,130601433 | 17,35026587 | 26,80981162 |
| **RB1CC1** | 0,13029832 | 13,70944286 | 22,43632454 |
| **EWSR1** | 0,127721343 | 17,76724045 | 24,20349219 |

| | | | |
|---|---|---|---|
| **TAF4** | 0,125254379 | 15,49846489 | 22,05682291 |
| **DDX41** | 0,12235467 | 15,64491716 | 22,62088264 |
| **DDX1** | 0,121926076 | 17,50555986 | 25,18857919 |
| **SNRPD2** | 0,114629094 | 20,73880505 | 25,42263064 |
| **DUT** | 0,113900466 | 17,27218021 | 22,78022656 |
| **YBX3** | 0,113333741 | 16,20131351 | 22,26115146 |
| **AGO2** | 0,112786833 | 13,73025429 | 20,77507009 |
| **USP7** | 0,112322042 | 15,64728415 | 24,36899608 |
| **JUP** | 0,112106219 | 12,51719366 | 20,760904 |
| **PSMC5** | 0,111632348 | 17,14800646 | 23,74672725 |
| **SF3B3** | 0,10997119 | 20,53642114 | 27,86116233 |
| **PFN1** | 0,107272718 | 19,66309941 | 23,94389886 |
| **MYL6** | 0,101024739 | 16,77021651 | 21,17564376 |
| **NUDC** | 0,099490441 | 17,43556175 | 23,83640587 |
| **ANXA3** | 0,097300663 | 16,85614991 | 22,33772424 |
| **PSMD1** | 0,095055708 | 16,55094214 | 24,29525363 |
| **SNRPA1** | 0,09342498 | 18,15383733 | 22,48602645 |
| **SRRT** | 0,092297654 | 15,30104004 | 23,65623703 |
| **KHSRP** | 0,091598251 | 14,88397967 | 22,37665958 |
| **LASP1** | 0,090650183 | 16,77520302 | 22,23276598 |
| **SCYL2** | 0,086675108 | 19,68194068 | 27,26862547 |
| **IQGAP1** | 0,085347591 | 18,20824865 | 26,44682686 |
| **ACTC1** | 0,083928109 | 19,07429132 | 27,51713207 |
| **CANX** | 0,071735204 | 16,44049071 | 23,35507951 |
| **CPPED1** | 0,070403076 | 15,5365609 | 21,68719682 |
| **PSMD8** | 0,068945893 | 15,57379509 | 21,52244999 |
| **PSMD11** | 0,067771798 | 18,13255328 | 24,34854339 |
| **ARPC4** | 0,067156792 | 15,84256314 | 20,01247632 |
| **RPS23** | 0,065926077 | 20,51030767 | 24,75033393 |
| **CAPN1** | 0,061310785 | 14,13290131 | 21,28021647 |
| **SNRPE** | 0,060863622 | 19,35527223 | 23,47944609 |
| **EEF1D** | 0,056893314 | 18,4290021 | 24,08826181 |
| **LARP1** | 0,055565883 | 16,50669808 | 24,37517522 |
| **ACTR1A** | 0,054994283 | 16,53484999 | 23,0010588 |
| **UGGT1** | 0,054633131 | 13,08364553 | 20,7191682 |
| **PAXBP1** | 0,052568921 | 14,55776284 | 21,60035335 |
| **ACTR2** | 0,051752606 | 17,84125925 | 23,33409066 |
| **DCTPP1** | 0,049421668 | 15,71384292 | 20,82384301 |
| **MDH2** | 0,048436124 | 17,64342407 | 23,19491853 |
| **LANCL1** | 0,046854229 | 15,53803733 | 21,06955291 |
| **TTLL12** | 0,046141738 | 14,69855998 | 22,18107459 |
| **ACAT1** | 0,042737716 | 17,87061356 | 23,84086166 |
| **PSMA4** | 0,042401525 | 19,89754612 | 25,57282557 |
| **DDX3X** | 0,038422129 | 20,71879205 | 27,54603304 |
| **RPS25** | 0,037311898 | 21,47735577 | 24,81178063 |
| **PSMC3** | 0,037171309 | 15,51605907 | 22,26862309 |
| **GOT2** | 0,036608833 | 17,14605152 | 23,28913465 |
| **TAGLN2** | 0,035356481 | 14,3244853 | 19,02493042 |
| **LDHA** | 0,034600706 | 18,79884303 | 24,37178994 |
| **HNRNPK** | 0,031945614 | 20,36066136 | 26,66761415 |

| | | | |
|---|---|---|---|
| **CIRBP** | 0,031917423 | 16,31733606 | 21,33276212 |
| **PSMC2** | 0,02840413 | 17,03459086 | 22,84193028 |
| **PRMT1** | 0,027054996 | 19,03434404 | 25,25731475 |
| **ACTB** | 0,026020971 | 23,1674794 | 29,70746961 |
| **MYL12B** | 0,025170576 | 16,88010961 | 21,0500169 |
| **BUB3** | 0,02453953 | 17,93307588 | 23,04416863 |
| **ELMO2** | 0,022658986 | 19,90970561 | 27,10729275 |
| **USO1** | 0,022325182 | 14,10344627 | 21,38991557 |
| **PSMD7** | 0,02179712 | 16,61790265 | 22,78047355 |
| **STK38L** | 0,020105083 | 18,91768286 | 25,73360665 |
| **BCLAF1** | 0,01915395 | 16,71500611 | 23,77722588 |
| **SEC24C** | 0,018684019 | 14,7108961 | 21,71394917 |
| **SF3B1** | 0,015721157 | 19,6514208 | 28,15465319 |
| **PFAS** | 0,014983688 | 14,81187638 | 22,74208022 |
| **PYM1** | 0,014833729 | 16,37865047 | 21,92711339 |
| **CTTN** | 0,013698076 | 18,55743752 | 25,0667477 |
| **PA2G4** | 0,013555125 | 18,84098071 | 25,28001347 |
| **GDI2** | 0,011190252 | 17,54805425 | 24,10874224 |
| **LDHB** | 0,00792739 | 18,90971538 | 24,10985828 |
| **UBAP2** | 0,006807797 | 14,73365092 | 22,02551179 |
| **PYGB** | 0,006630779 | 18,29950616 | 25,85888037 |
| **PPIB** | 0,006513478 | 19,14363565 | 23,91503346 |
| **SF3B2** | 0,005773641 | 19,25075486 | 27,1279002 |
| **XRCC6** | 0,004163391 | 18,63036277 | 25,54004272 |
| **CAPZA1** | 0,004096224 | 18,54719203 | 24,56821252 |
| **EEF2** | 0,002488877 | 21,03962237 | 28,20634272 |
| **AHCY** | 0,000778828 | 18,08079324 | 23,90039661 |
| **XRCC5** | -0,002021222 | 17,23158996 | 25,32202991 |
| **ABCF1** | -0,002505209 | 16,4607705 | 23,22543929 |
| **GLRX3** | -0,003085829 | 18,19929532 | 24,10224144 |
| **TNRC6B** | -0,003444967 | 15,1621815 | 23,94321329 |
| **PAICS** | -0,005565071 | 20,17960947 | 25,80834743 |
| **SART1** | -0,005849939 | 16,3530618 | 22,97827498 |
| **PLCG1** | -0,006130041 | 15,69721194 | 23,57212395 |
| **RPS12** | -0,008587881 | 19,83785888 | 25,29854349 |
| **GTPBP10** | -0,008849149 | 15,48250337 | 22,51082919 |
| **STK38** | -0,009693763 | 15,90865922 | 23,57597096 |
| **MYH14** | -0,01015858 | 13,84408457 | 21,5305689 |
| **GAPDH** | -0,010560629 | 21,60461385 | 27,55063916 |
| **MRPL44** | -0,011883739 | 19,74347522 | 25,46519421 |
| **LMO7** | -0,012470517 | 15,55666471 | 24,19630515 |
| **CRKL** | -0,013094022 | 17,73276586 | 24,14647048 |
| **SEPTIN2** | -0,013494416 | 13,93510365 | 20,75895404 |
| **THRAP3** | -0,014484992 | 18,34148118 | 25,55753062 |
| **RPS15A** | -0,017329446 | 20,53910642 | 25,49495045 |
| **FAM98B** | -0,017395199 | 16,40274007 | 22,30292298 |
| **DOCK9** | -0,017935523 | 12,03288376 | 22,44675956 |
| **SBF1** | -0,020077133 | 11,61771231 | 19,8100245 |
| **CACYBP** | -0,020640444 | 15,81419858 | 22,99528433 |
| **HNRNPA2B1** | -0,020852648 | 18,10940559 | 24,07452652 |

| | | | |
|---|---|---|---|
| ENO1 | -0,021233253 | 19,48941811 | 25,26496845 |
| GADD45GIP1 | -0,021716527 | 16,89593023 | 21,68495917 |
| PKM | -0,025404628 | 16,80846986 | 23,84638819 |
| SF1 | -0,027189675 | 15,71811363 | 21,8371762 |
| RPS13 | -0,030329447 | 20,80156891 | 25,94152778 |
| DOCK4 | -0,031272688 | 19,92841042 | 28,59422508 |
| ADISSP | -0,03182814 | 17,12168527 | 22,42770705 |
| CFL1 | -0,033215184 | 20,38039906 | 25,51387366 |
| HSD17B4 | -0,033215184 | 14,61872962 | 21,01102064 |
| DHX29 | -0,034367178 | 14,86529831 | 23,22412948 |
| TMOD3 | -0,034953336 | 16,83166187 | 23,37731734 |
| ARF1 | -0,034965656 | 17,25684752 | 21,57880632 |
| CS | -0,036895827 | 17,9061018 | 23,15404858 |
| CAPN2 | -0,037369596 | 15,1449901 | 21,93663315 |
| PSMA1 | -0,040237233 | 16,1871394 | 21,93598606 |
| CARS1 | -0,040430142 | 15,8259277 | 22,18345931 |
| SRSF5 | -0,040838166 | 18,63523631 | 23,04968461 |
| PGD | -0,04126123 | 16,33405792 | 22,05490368 |
| CHID1 | -0,043943305 | 17,71768535 | 23,26042896 |
| GMPS | -0,044962529 | 16,43731861 | 23,67952162 |
| RPS24 | -0,046091643 | 21,30838496 | 25,03661801 |
| TARS1 | -0,046392057 | 14,8388264 | 21,68528719 |
| RPS19 | -0,047646538 | 21,18036043 | 25,68985398 |
| LCP1 | -0,047890116 | 15,2145721 | 22,07665905 |
| PDIA4 | -0,047974626 | 16,7964892 | 23,51700388 |
| RPS6 | -0,04865092 | 22,39168203 | 26,46731569 |
| U2AF1 | -0,049989534 | 18,24301898 | 24,23432628 |
| TSR1 | -0,050248436 | 17,85151929 | 24,56445717 |
| CHD1 | -0,051414073 | 14,60655173 | 22,21954315 |
| ACTN1 | -0,051668285 | 16,14080315 | 24,02304274 |
| PMVK | -0,052311389 | 15,88152825 | 20,87375149 |
| RPS17 | -0,053049624 | 21,38807424 | 25,82539995 |
| RPS4X | -0,054706919 | 21,95684547 | 27,07670253 |
| MRPL45 | -0,055761248 | 18,31926344 | 25,34032571 |
| RBM17 | -0,056871298 | 16,43566266 | 22,12086602 |
| WDR26 | -0,060362143 | 16,637573 | 24,49106038 |
| HDGF | -0,062263947 | 17,39877152 | 22,64817908 |
| RPS16 | -0,062545117 | 20,78961059 | 25,48072431 |
| MRPL15 | -0,064007284 | 19,52593876 | 24,93024549 |
| ARHGAP21 | -0,064683583 | 12,98150782 | 21,3697372 |
| RPS5 | -0,065128782 | 20,78609623 | 25,37238792 |
| UBE2N | -0,066034672 | 17,05566727 | 22,87028093 |
| TPP2 | -0,066057267 | 17,43009859 | 25,50223388 |
| EIF3K | -0,06607239 | 19,21897152 | 24,06874417 |
| SPIN3 | -0,072467555 | 15,77121863 | 21,87555239 |
| EIF4A3 | -0,073423577 | 15,0635903 | 21,07478782 |
| MRPS30 | -0,074129683 | 18,54798775 | 24,16834179 |
| SSRP1 | -0,075129881 | 20,0352712 | 26,20871646 |
| DOCK7 | -0,076366546 | 16,58152916 | 24,81571059 |
| RPS3A | -0,077274365 | 20,63654379 | 26,49840909 |

| | | | |
|---|---|---|---|
| **TXN** | -0,077786965 | 20,83163352 | 24,7524453 |
| **RPS2** | -0,079594894 | 20,97943264 | 26,58969911 |
| **SNRPA** | -0,080352303 | 16,37827792 | 22,69246368 |
| **UBA1** | -0,080759076 | 16,7867631 | 24,19870481 |
| **AK2** | -0,08204894 | 16,08821543 | 22,27598995 |
| **RPS8** | -0,082871254 | 20,80064822 | 25,21965639 |
| **PRDX1** | -0,084777416 | 22,15746763 | 27,35514394 |
| **RPS9** | -0,086967018 | 20,76511448 | 25,38354891 |
| **GID8** | -0,089170132 | 18,00388021 | 22,93192319 |
| **TXNDC12** | -0,089858379 | 15,91398796 | 20,8464392 |
| **MTMR14** | -0,091284455 | 17,97772414 | 24,37263147 |
| **MYH10** | -0,093094041 | 13,11374217 | 22,11410197 |
| **EIF5B** | -0,093455658 | 16,85817531 | 25,29540546 |
| **CBR1** | -0,097571336 | 16,41880793 | 21,5062514 |
| **MYH9** | -0,098309891 | 18,37088603 | 26,96585766 |
| **SLAIN2** | -0,101636875 | 16,77214701 | 23,39762137 |
| **MRPL47** | -0,102829283 | 18,86350875 | 24,02027555 |
| **SWAP70** | -0,103022988 | 16,1176431 | 24,41536285 |
| **GGH** | -0,103286413 | 16,95868864 | 22,70473504 |
| **EZR** | -0,10495595 | 17,57557864 | 23,85709334 |
| **SF3A1** | -0,105452501 | 18,68539215 | 26,42864331 |
| **U2SURP** | -0,105690524 | 19,03743792 | 26,1794771 |
| **PFN2** | -0,107507905 | 17,57092547 | 21,48105703 |
| **SRSF1** | -0,108238635 | 17,98683828 | 23,10488792 |
| **NACA** | -0,108884141 | 13,74062404 | 21,4685498 |
| **EIF2S1** | -0,10891785 | 19,71175882 | 25,82107456 |
| **RPS21** | -0,109327656 | 18,44995599 | 22,17031467 |
| **MRPL27** | -0,110993817 | 17,16458944 | 22,03586049 |
| **LRCH1** | -0,111544445 | 16,42963558 | 23,32425692 |
| **OTUD4** | -0,114488095 | 18,32377607 | 26,0734441 |
| **DDX46** | -0,114972386 | 15,98709373 | 22,87181051 |
| **RPS18** | -0,115706861 | 21,50258192 | 25,6839336 |
| **DDX42** | -0,119040276 | 18,2477792 | 26,2257114 |
| **MRPL4** | -0,119212587 | 20,04762763 | 26,64670691 |
| **ACTBL2** | -0,119311842 | 21,99241146 | 27,38470199 |
| **PPM1B** | -0,119988457 | 17,79945449 | 23,65486864 |
| **RPS11** | -0,120137422 | 19,72325208 | 24,33858134 |
| **MRPL49** | -0,120709877 | 18,6759134 | 23,23004605 |
| **CAMSAP3** | -0,120743863 | 18,05726728 | 26,49188612 |
| **GTF2I** | -0,122538363 | 16,27932643 | 24,03316262 |
| **SUPT16H** | -0,124078423 | 20,19897652 | 27,52882659 |
| **MRPL22** | -0,124345736 | 18,8637608 | 23,54510808 |
| **GPX4** | -0,125171561 | 15,69477502 | 21,61925638 |
| **PRPS1** | -0,126711729 | 18,90031061 | 25,18114032 |
| **RPS10** | -0,127095083 | 21,15130605 | 25,7327876 |
| **RACK1** | -0,127609867 | 20,97256851 | 27,32130745 |
| **EIF5A** | -0,127880467 | 20,61406186 | 26,89079777 |
| **PRRC2C** | -0,130728716 | 13,94343056 | 23,48212452 |
| **POLDIP2** | -0,132016567 | 15,61758225 | 21,47248856 |
| **SF3B6** | -0,132103545 | 17,74364231 | 22,25781827 |

| | | | |
|---|---|---|---|
| **MRPL23** | -0,13347181 | 19,09198971 | 23,69173273 |
| **PLPBP** | -0,134793835 | 16,56091493 | 21,73142953 |
| **USP15** | -0,13558073 | 17,59542747 | 24,95712897 |
| **KIF5B** | -0,135696941 | 15,59525748 | 23,12575545 |
| **ABCE1** | -0,137489103 | 16,60146613 | 22,6918321 |
| **MRPL18** | -0,138417751 | 19,05890249 | 23,24667102 |
| **MRPL1** | -0,139341724 | 19,37377286 | 24,87242439 |
| **VPS25** | -0,139792067 | 14,43860778 | 20,3745386 |
| **CPSF6** | -0,140555167 | 18,84604125 | 24,57604018 |
| **RANBP10** | -0,141202006 | 18,06815547 | 23,72928715 |
| **MRPL39** | -0,142172888 | 20,0131659 | 26,03951444 |
| **WDR77** | -0,143399353 | 26,29488568 | 31,76240064 |
| **RPS3** | -0,145009619 | 22,88531183 | 28,54803165 |
| **SNRPD1** | -0,14547765 | 22,80231138 | 26,42734367 |
| **ACLY** | -0,145517551 | 14,58643325 | 22,40776877 |
| **PRDX6** | -0,145850057 | 19,09632825 | 24,60542647 |
| **PPIL4** | -0,146653668 | 18,98027441 | 25,56770979 |
| **ANXA2** | -0,147010398 | 20,30054174 | 26,38561282 |
| **FLNB** | -0,151121703 | 15,60822594 | 24,88988541 |
| **CMAS** | -0,155914346 | 17,76092132 | 24,79813042 |
| **PGAM5** | -0,156134044 | 20,25909115 | 25,96900936 |
| **PRPS2** | -0,156166208 | 18,27140527 | 24,94443517 |
| **EIF3H** | -0,158316651 | 21,31672531 | 28,16844381 |
| **PRMT5** | -0,158316683 | 25,7659191 | 33,37450775 |
| **SLIRP** | -0,158421352 | 16,74432143 | 21,33977318 |
| **EIF1AX** | -0,15877558 | 16,90618598 | 21,21286127 |
| **PRDX2** | -0,161483802 | 16,22656272 | 23,48772409 |
| **DHX15** | -0,16281026 | 20,73490859 | 27,57900301 |
| **LGALS3** | -0,162842541 | 18,51514542 | 24,62581733 |
| **GSPT1** | -0,162985215 | 16,72089441 | 23,34461648 |
| **WARS1** | -0,164248363 | 16,28818719 | 22,93324473 |
| **ZRANB2** | -0,167166711 | 18,1927744 | 24,68362133 |
| **TRIM21** | -0,167858005 | 21,52022605 | 27,93620104 |
| **MRPL21** | -0,16874703 | 18,84097047 | 23,99897783 |
| **CLNS1A** | -0,170226207 | 23,48754133 | 28,76914394 |
| **RPSA** | -0,170978522 | 21,78719954 | 27,52454877 |
| **RPS14** | -0,171579649 | 21,9668605 | 26,03537391 |
| **MRPL43** | -0,172847618 | 18,33453809 | 23,36088101 |
| **NME2** | -0,17322984 | 15,36472745 | 22,31096773 |
| **PTPN11** | -0,173262371 | 15,36815402 | 22,55185781 |
| **MRPL28** | -0,175598726 | 18,91168651 | 24,54329179 |
| **PSAT1** | -0,175940925 | 16,12803461 | 21,88529321 |
| **ROCK1** | -0,177995958 | 18,7257601 | 26,92711339 |
| **CHERP** | -0,178605435 | 17,89071552 | 25,45400102 |
| **GSPT2** | -0,180969523 | 15,65550277 | 21,462871 |
| **MRPL58** | -0,18174658 | 20,15503231 | 24,83849122 |
| **RPS7** | -0,182447678 | 20,36558557 | 26,87751565 |
| **H4C1** | -0,184763405 | 18,98171035 | 22,56667285 |
| **MRPL41** | -0,184982067 | 19,03094906 | 23,97622497 |
| **RIOK1** | -0,185747628 | 21,33153441 | 28,17955936 |

| | | | |
|---|---|---|---|
| **ACTR3** | -0,18640416 | 15,60793131 | 21,90502066 |
| **MRPL50** | -0,186828328 | 18,03930107 | 23,12594985 |
| **APEX1** | -0,187099286 | 15,70734116 | 22,41450288 |
| **RBM10** | -0,188857819 | 22,07036011 | 29,21482673 |
| **ARHGEF10** | -0,190149313 | 14,80005727 | 23,15261557 |
| **EIF3D** | -0,190733734 | 21,95409855 | 29,32184502 |
| **MRPL3** | -0,191326656 | 16,89992725 | 24,5788447 |
| **UPP1** | -0,195161707 | 15,36070994 | 21,23397536 |
| **EIF4G1** | -0,197327008 | 18,15313636 | 26,05355097 |
| **FKBP1A** | -0,198228795 | 17,8447977 | 21,21860738 |
| **MRPL11** | -0,199810343 | 19,79485158 | 24,64214544 |
| **MRPL42** | -0,203925576 | 16,48571927 | 22,11882957 |
| **RBM8A** | -0,204482378 | 16,06437789 | 21,56699794 |
| **PCBD1** | -0,205430244 | 16,13264699 | 20,63564373 |
| **ACAP2** | -0,205685315 | 20,34183006 | 27,72216686 |
| **TKT** | -0,205957127 | 15,80057976 | 23,60594636 |
| **LRRFIP1** | -0,20787762 | 15,08234312 | 22,38341707 |
| **EIF3F** | -0,208733258 | 22,74859894 | 28,34112922 |
| **MRPL48** | -0,209536554 | 19,49780247 | 24,28234867 |
| **EIF3J** | -0,211726342 | 20,5345543 | 25,87995138 |
| **MRPL19** | -0,212606537 | 18,96922817 | 24,94079629 |
| **BOLA2** | -0,217390145 | 19,75568195 | 24,67537074 |
| **SND1** | -0,218178721 | 18,93203977 | 26,44581184 |
| **PRPSAP2** | -0,218654295 | 19,73636174 | 26,1799769 |
| **MRPL32** | -0,21870743 | 16,90080393 | 21,60905426 |
| **MRPL37** | -0,225947423 | 20,24889772 | 26,88403028 |
| **SPIN1** | -0,227033358 | 19,94590482 | 25,88612163 |
| **MRPL53** | -0,230084442 | 18,00899389 | 22,49122353 |
| **NME1** | -0,234202154 | 19,725617 | 25,01917284 |
| **MRPL38** | -0,237597346 | 19,75003851 | 25,80441374 |
| **PGK1** | -0,238436696 | 19,37849243 | 26,23753909 |
| **EIF3L** | -0,238992824 | 21,69706937 | 28,16598686 |
| **DHX38** | -0,242043721 | 17,05285811 | 24,64141059 |
| **ROCK2** | -0,242353682 | 11,83594512 | 19,71619539 |
| **EIF3M** | -0,243079186 | 22,19818722 | 28,35765158 |
| **ALDOA** | -0,247023118 | 17,59233537 | 24,16627425 |
| **PPA1** | -0,251495817 | 17,92624966 | 24,71715399 |
| **COL12A1** | -0,251667626 | 12,97265078 | 21,73245122 |
| **EIF3B** | -0,252128553 | 23,06415473 | 30,15024481 |
| **MRPL2** | -0,252561038 | 16,61827078 | 24,383483 |
| **VCP** | -0,253578316 | 18,10848702 | 25,08894919 |
| **EIF3E** | -0,254335283 | 22,1973345 | 28,84296056 |
| **CLIC1** | -0,255459966 | 16,67260029 | 21,98355544 |
| **EIF3I** | -0,260307414 | 21,50381636 | 28,28642033 |
| **YARS2** | -0,262027713 | 16,566832 | 23,4340774 |
| **GOLGA3** | -0,263120984 | 16,30002777 | 24,5127792 |
| **SRSF11** | -0,263542326 | 16,92964369 | 22,67633963 |
| **MCM5** | -0,265124924 | 17,50472331 | 24,80306727 |
| **EIF3A** | -0,266003679 | 22,87430638 | 30,6772709 |
| **EIF3C** | -0,269152849 | 22,34886302 | 29,69731325 |

| | | | |
|---|---|---|---|
| **PRPSAP1** | -0,269854236 | 17,88745467 | 24,13729992 |
| **CSTB** | -0,274363517 | 17,5695738 | 23,42734047 |
| **LGALS1** | -0,275733888 | 18,88985447 | 23,73190249 |
| **GNPNAT1** | -0,276520072 | 17,01599573 | 22,21187166 |
| **IVNS1ABP** | -0,283497739 | 17,92037806 | 25,21750091 |
| **TMPO** | -0,283860706 | 16,03543633 | 23,20349717 |
| **MRPL46** | -0,287932555 | 17,8962851 | 24,39283518 |
| **PRRC2A** | -0,288390576 | 15,697583 | 25,13176425 |
| **PSMB5** | -0,298069513 | 14,32596125 | 19,23285965 |
| **DYNC1I2** | -0,301514799 | 14,45226269 | 20,3697088 |
| **TJP1** | -0,304338721 | 15,42373091 | 23,50495522 |
| **MRPL17** | -0,305743829 | 17,67381054 | 22,8087893 |
| **CCDC88C** | -0,306930159 | 18,38269438 | 27,22747522 |
| **LYAR** | -0,309413054 | 14,90917431 | 21,63978524 |
| **TPM4** | -0,314167468 | 17,09805265 | 22,27793928 |
| **SERPINB1** | -0,317975152 | 16,77987749 | 22,68416018 |
| **RANBP9** | -0,322168583 | 17,29124566 | 23,370642 |
| **EIF4B** | -0,326395317 | 22,65589141 | 29,56700254 |
| **EIF3G** | -0,328217481 | 21,39667219 | 27,70949233 |
| **TPM3** | -0,329721627 | 16,45670734 | 21,45671871 |
| **SPTAN1** | -0,332325795 | 14,86972235 | 24,65567309 |
| **MRPL40** | -0,343137546 | 18,87554739 | 24,75428172 |
| **ACTN4** | -0,343640898 | 18,47029213 | 25,96643721 |
| **GGCT** | -0,344538199 | 14,97733956 | 20,54673372 |
| **U2AF2** | -0,348647485 | 19,2552038 | 24,8357315 |
| **POF1B** | -0,352860507 | 15,76647715 | 21,76533575 |
| **QPCT** | -0,360977816 | 18,22887348 | 23,91922293 |
| **CCAR1** | -0,362361922 | 15,85286271 | 22,66970948 |
| **SPTBN1** | -0,363233901 | 15,28170066 | 24,6094714 |
| **FLNA** | -0,365381946 | 20,74455165 | 29,83926545 |
| **R3HDM1** | -0,380286552 | 14,85569583 | 21,95314305 |
| **PRRC2B** | -0,381563973 | 12,52737928 | 21,64681767 |
| **ZYX** | -0,385817853 | 15,62440519 | 24,4373325 |
| **EXOSC8** | -0,389709407 | 19,11268385 | 24,20875986 |
| **PRPF31** | -0,391638625 | 16,11321039 | 22,13636661 |
| **RTF2** | -0,394818645 | 15,61177985 | 22,10773174 |
| **TJP2** | -0,412615801 | 15,31238666 | 24,30449746 |
| **ARHGAP23** | -0,436246376 | 12,32158993 | 21,4896729 |
| **CDV3** | -0,459357679 | 15,85161079 | 23,95103624 |
| **E2F7** | -0,469299938 | 13,77312203 | 21,70269352 |
| **PLS3** | -0,473680763 | 13,63602411 | 21,17842252 |
| **MRPL24** | -0,474702871 | 18,51776164 | 24,18359056 |
| **MSN** | -0,483056979 | 13,74167764 | 21,27976321 |
| **SF3A2** | -0,497577762 | 16,55952149 | 22,68331792 |
| **MKLN1** | -0,502556283 | 15,01148498 | 21,89929053 |
| **ANXA1** | -0,505748235 | 17,74237304 | 23,59520542 |
| **EXOSC6** | -0,525690252 | 18,9640174 | 24,16566108 |
| **PUF60** | -0,548722974 | 15,1649462 | 22,84113689 |
| **SVIL** | -0,551197696 | 14,82914131 | 22,9562133 |
| **KIF11** | -0,557454515 | 24,20309385 | 31,83830888 |

| | | | |
|---|---|---|---|
| **PPIA** | -0,557461623 | 21,92046044 | 26,65194281 |
| **EIF4A2** | -0,565876205 | 15,88543423 | 22,43715467 |
| **SPINDOC** | -0,622420877 | 17,78303872 | 24,4967189 |
